# Neotaphonomic characteristics of vertebrate site formation in underwater caves

**DOI:** 10.64898/2026.02.13.705854

**Authors:** Meg M Walker, Joanne E Wilkinson, Mathew Stewart, Geraldine E Jacobsen, Shwaron Kumar, Vladimir Levchenko, Stewart Fallon, Rebecca Esmay, Rachel Wood, Justyna J Miszkiewicz, Gilbert Price, Elizabeth Reed, Joeseph Monks, Julien Louys

## Abstract

Recovering well-preserved vertebrate remains from underwater caves has provided critical insights into archaeological and palaeontological records worldwide. However, understanding how bone assemblages form and are modified in underwater environments remains limited due to stable low energy burial conditions that produce time-averaged deposits, and underwater settings that hinder traditional recording and recovery methods. This study applies an actualistic taphonomic framework to three assemblages of domesticate animal bones (N = 231) from two underwater caves, Green Waterhole and Gouldens Sinkhole, near Mount Gambier, South Australia, encompassing known submerged (wet; N = 134) and dry (N = 97) burial conditions. The assemblages were examined to assess how wet and dry cave environments impact bone distribution, surface and microstructural modification. Radiocarbon dating of 41 specimens indicates that domesticate fauna were deposited over decadal and centennial timescales, allowing taphonomic signatures to be contextualised through time. Statistically significant differences were identified between wet and dry burial contexts. Bones recovered from wet contexts exhibit mostly better preservation, including skeletal elemental completeness, surface, and microstructure, than those from dry caves. However, some of the submerged specimens also have elevated frequencies of bone surface corrosion with macroscopic evidence for heterogenous black biological staining, algal or biofilm attack, and a distinctive form of circular etching. Histotaphonomy further reveals patterns of peripheral cyanobacterial tunnelling across most bones recovered from submerged contexts. Bones from dry environments were dominated by terrestrially linked tunnelling across all regions of the bone cortex. These findings can be explained by variation in light availability across different cave zones which influences biological activity and, in turn, the expression of taphonomic markers on bone externally and at the microstructural level. This is the first study to provide a benchmark bone dataset for reconstructing depositional histories and post-depositional reworking in underwater cave environments under a taphonomic framework.

## Introduction

Bones recovered from aquatic landscapes are often exceptionally well-preserved, providing some of the best known assemblages for revealing the lives of past people and ecological communities [1–5]. This has been particularly evident in underwater caves in near shore karst environments [6]. However, the origin and site formation processes of these sites have rarely been studied, hindering archaeological and palaeontological investigations [7].

Underwater caves can be environmentally stable, with low water flow rates, light exposure limited to cave entrances, and have thermally and chemically distinct water across different areas [7, 8]. Sediment accumulation is generally limited to allochthonous depositions at cave entrances and locally slow autochthonous accumulations throughout. These conditions can result in unstratified, comingled, and time-averaged deposits of exceptionally preserved remains across the surface of cave floors.

A lack of stratification in underwater cave deposits further hinders comprehensive investigations of changes in faunal and archaeological change through time. Key to near shore phreatic (underwater) caves environments are shifting water tables, from relatively small seasonal fluctuations to extreme eustatic changes that occur with glacial-interglacial cycles [9]. The presence or absence of water in these caves alters site formation processes and taphonomic histories, thus identifying wet or dry deposition in such sites could aid in narrowing the timing of depositional windows [7].

Determining when and how a bone is modified under different conditions is informed by actualistic studies that establish chronologically controlled modern comparative benchmarks [10, 11]. Early depositional changes can be preserved through deep time, captured by the fossilisation process that records changes in the initial burial environments [12, 13]. However, to date no actualistic studies have directly identified the effects of different underwater cave landscapes, such as variations in physical structures, hydrology and chemistry, biological communities, and light exposure on the taphonomy of vertebrate remains [7].

Actualistic taphonomy in either cave or aquatic settings has traditionally focused on taxonomic composition (e.g. [3, 4, 14]) and decomposition patterns (e.g. [15, 16, 17]), with only a few investigations identifying modifications to bone surfaces and microstructure [18–20]. Examining bone surface modifications (BSMs) and histotaphonomy, the taphonomic study of tissue microstructure, may be essential in underwater cave settings to fill the gaps created by working in environments prone to ‘silting out’, and thus limiting traditional excavation and recording methods (6). For example, BSMs preserve layers of alterations associated with the death of animals, movement or disturbance of remains, and interactions with local burial agents such as sediment, chemistry, and microbiota [21]. Linking BSM morphology and expressions of modification to taphonomic agents creating these changes has shown differences between permanently wet, dry, or changing hydrological and burial conditions in a temperate landscape [19]. Across a thirty-year actualistic study in Neuadd, Wales, submerged bones exhibited no rodent gnawing but did show corroded surfaces, and biotic attack by moss, algae, and lichen which resulted in different corrosive patterns and in black stains across bone surfaces [19].

Aquatic-specific taphonomic agents have also been identified across experimental and observational studies with different hydrological and site conditions [7], but many of these are not applicable to underwater cave systems. Low water flow in many submerged conduits and sinkholes will limit sediment abrasion and polishing across bones that is typically reflective of high energy environments [22–24]. Compared to marine and fluvial landscapes, flora and fauna are limited in terrestrial underwater caves, further restricting the degree of degradation to bone surfaces [25–28]. Floral communities that can produce pitting, staining, and general corrosion [19, 21] are likely to occur only in the entrance zones of cave systems due to light availability [8].

In histotaphonomy, measures of bone histological integrity have been used to study burial environments [29–31]. Aquatic diagenesis, the structural alteration of biomineralised tissues [30, 32, 33], differs from terrestrial, dry diagenesis. Whilst bacterial diagenesis has been shown to be decoupled from burial conditions in terrestrial settings [34], the DNA of aquatic cyanobacteria has been linked to a distinct form of microstructural bone tunnelling (Wedl-Tunnelling) [35]. Proliferation of radial microcracks across the secondary osteon cement line, a highly mineralised boundary of a fundamental cortical bone building unit, has also been linked to decay of bone collagen underwater [36], and used to infer aquatic submersion in palaeontological deposits [37, 38]. Embedded foraminifera were also identified in bone microstructures from aquatic settings, but these are limited to marine environments and sea caves [39].

Biotic modifications and loss of the organic content of bone are also linked to early stages of diagenesis in aquatic environments [13, 36, 40]. However, the timing and expression of diagenesis is inconsistent in submerged and terrestrial forensic settings [34, 41, 42]. For example, in an experimental study of submerged domesticate sheep bones under different chemical conditions, wet bones experienced greater collagen loss and increased porosity after 12 months compared to those left in dry settings [43]. Cyanobacteria Wedl-tunnelling occurred in waterlogged sand as early as four weeks [43], although its expression tends to vary across environmental conditions as reported by others [42]. Identifying when in the early diagenetic period histotaphonomic features occur in underwater caves is likely dependant on the site type (sinkhole, conduit) and local conditions.

Thus, submersion will affect the spatial distribution, surface modification, and microstructural levels of bones deposited in such contexts [7]. Here, we examine the effects of the underwater conditions on bone preservation at two submerged cave systems in southeast South Australia, Green Waterhole (cave) and Gouldens Sinkhole (cenote), using an actualistic approach. We focus on the taphonomic diagenesis of bones to document patterns across bone surfaces and microstructure at different underwater cave sites, and within a cave site. Our aim is to identify patterns that distinguish early wet from dry bone modifications in caves, in the expectation that it will provide resolution of time-averaged fossil deposits in caves that experience fluctuating water levels due glacial-interglacial conditions.

## Materials and methods

### Study sites

Green Waterhole (5L81) and Gouldens Sinkhole (5L8) are two caves formed through phreatic karst dissolution and collapse processes in the Gambier Limestone Formation of southeast South Australia (Fig 1). Green Waterhole (5L81) is a doline collapse cave with two submerged conduits on the northwestern and southeastern sides of a dry silt talus cone (Fig 2; [44]). The dry, central doline is exposed to aerial conditions, without an overhead environment, whilst the margin of the doline is protected by the entrance zone ceiling. The cave is situated adjacent to a highway, surrounded by pastoral fields and pine forests, with the perimeter of the site fenced off. A large assemblage of vertebrate remains associated with extinct, extant and domestic taxa was collected from the southeastern underwater cave [4, 45, 46].

**Fig 1.**
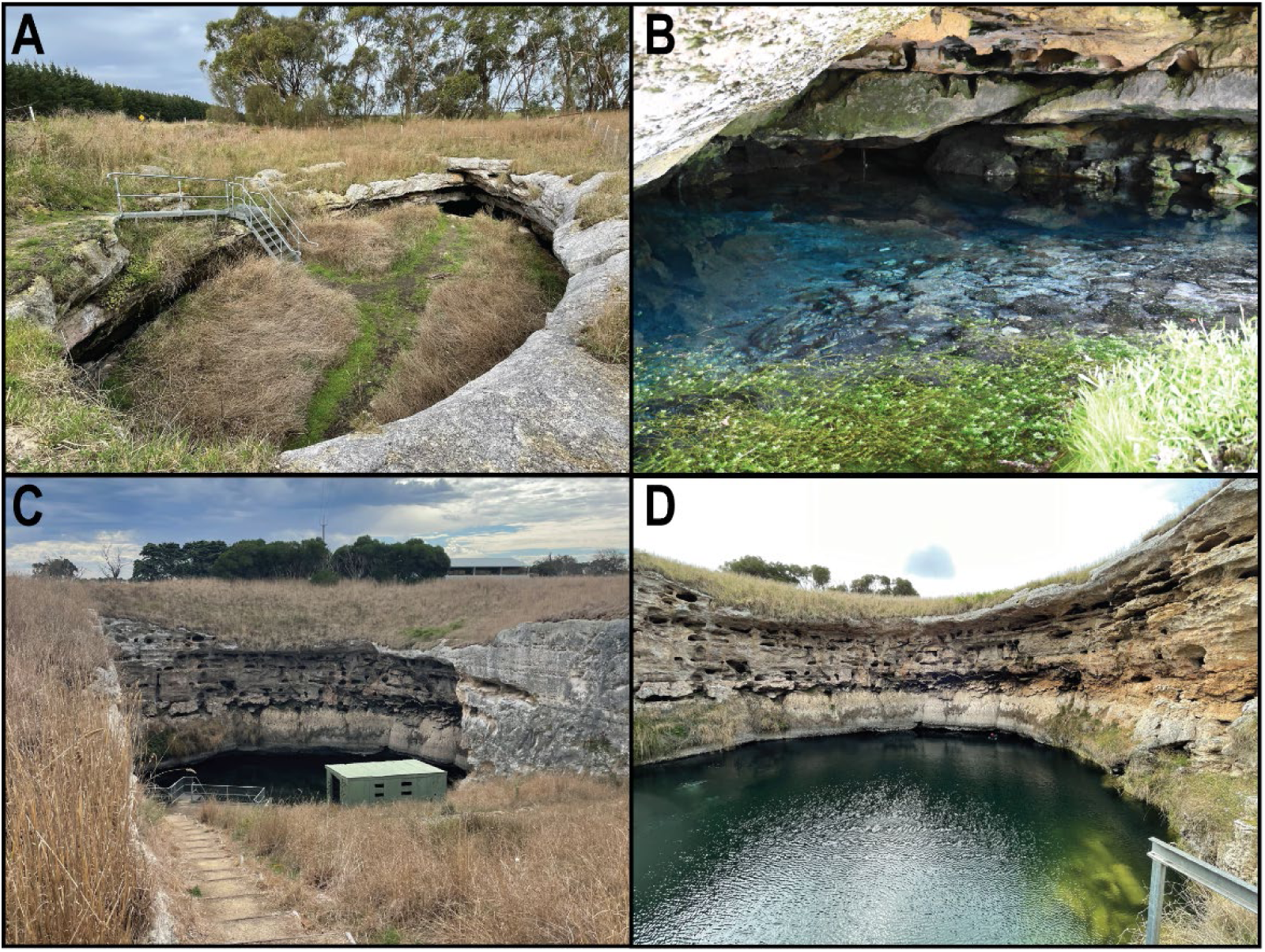
A: Green Waterhole (5L81) collapsed doline looking east; B: Green Waterhole northwest lake with aquatic flora across margins; C: Gouldens Hole (5L8) looking north looking towards sinkhole across artificial ramp access path and pumphouse; D) Gouldens Sinkhole looking north across exposed lake towards vertical walls pocketed with phreatic, vertical tunnels.

**Fig 2.**
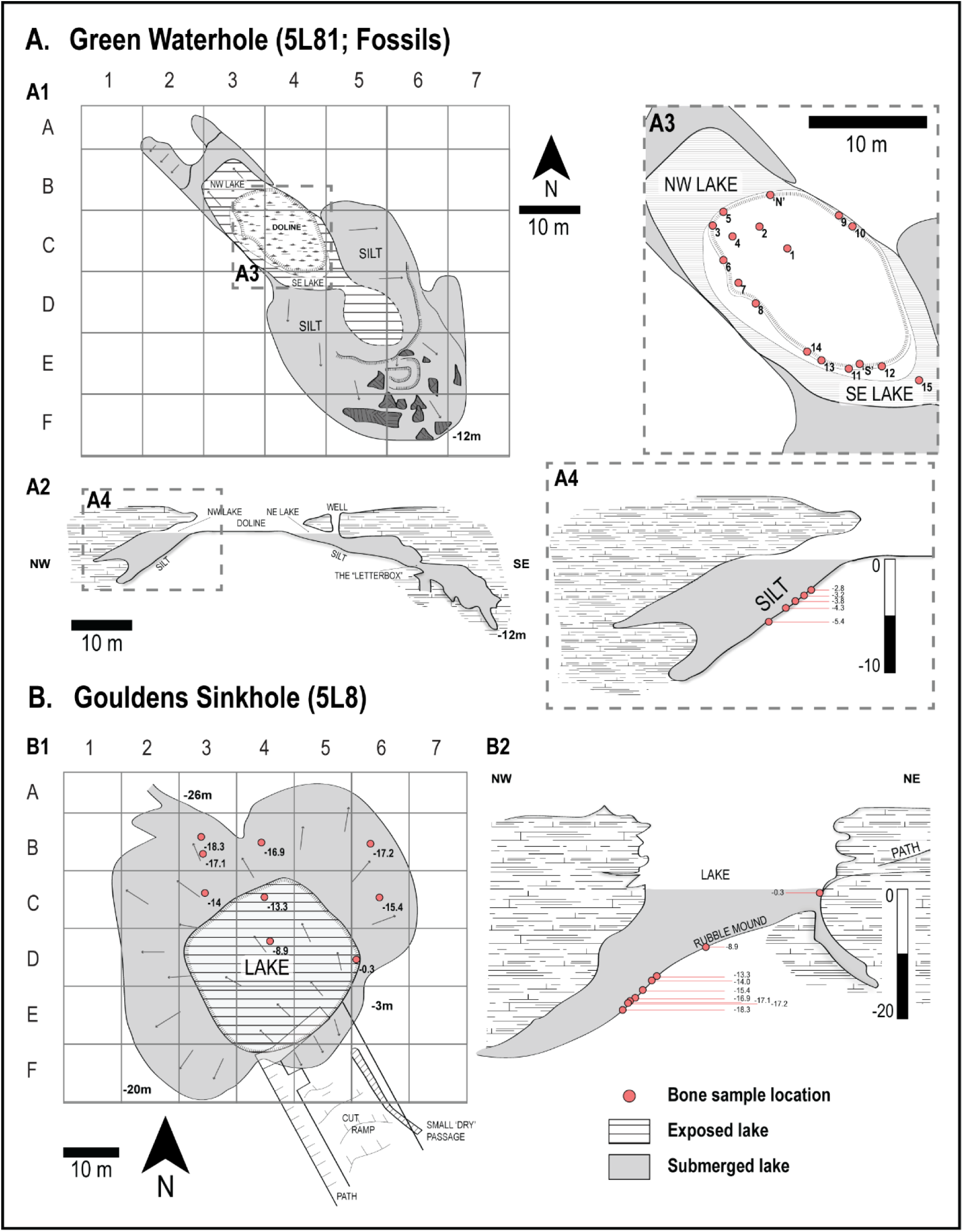
Site maps of Green Waterhole (A) and Gouldens Sinkhole (B), and associated recovery locations. A1) map of gridded Green Waterhole, plan view; A2) cross section view of Green Waterhole; A3) inset of A1, surface collection map across dry doline, plan view; A4) inset of A2, collection depths across Green Waterhole West Lake at Grid B3, cross section view; B1) gridded map of Gouldens Sinkhole with approximate locations and depths of collection points, plan view; B2) collection depths across Gouldens Sinkhole, cross section.

Gouldens Sinkhole is a round sinkhole, 29 metres in diameter at water surface and 63 metres below surface (Fig 1). The sheer vertical sides of the open sinkhole leads to a submerged overhead environment (Fig 2). The site, currently fenced off as a cave reserve, sits in a pastoral setting. A large access ramp was built into the side of the sinkhole (date unknown), likely to provide stock access to water. An abandoned historic pump house sits at the bottom of the ramp (1940s - 50s). Approximately 2,500 – 3,000 cubic meters of excavated rubble was pushed into the sinkhole, increasing the size of the central talus cone [47]. Material found on top of the talus cone postdates the path construction while deeper assemblages include bones deposited prior to the path’s construction.

Permissions to conduct scientific diver investigations were provided by the Department for Environment and Water (DEW) and the South Australian Heritage Council (Permit No. 0001/23) and supported by the Cave Divers Association of Australia.

### Assemblage

Recent (<250 years old) assemblages of skeletal elements were collected from Green Waterhole (GW; 5L81) and Gouldens Sinkhole (GH; 5L8) in April 2023 from submerged (wet) and surface (dry) burial contexts. Three distinct assemblages were collected, varying in hydrological context and depth (Table 1). Specimens were collected on top of the submerged cave floor sediment at different depths below the water’s surface (wet), and across the surface floor of the doline collapse area at GW (dry) (Fig 2). For each submerged site, bones associated with shallow depths were recovered near the entrance of the cave, while those at deeper points were under an overhead environment. Location and photographic context were recorded for each specimen prior to collection, however, silting out after collection limited visibility and thus the possibility of further in situ analyses. With reduced visibility and inability to use classic terrestrial field recording techniques, it was not possible to determine if bones from the same depths were articulated and belong to the same individual.

**Table 1.**
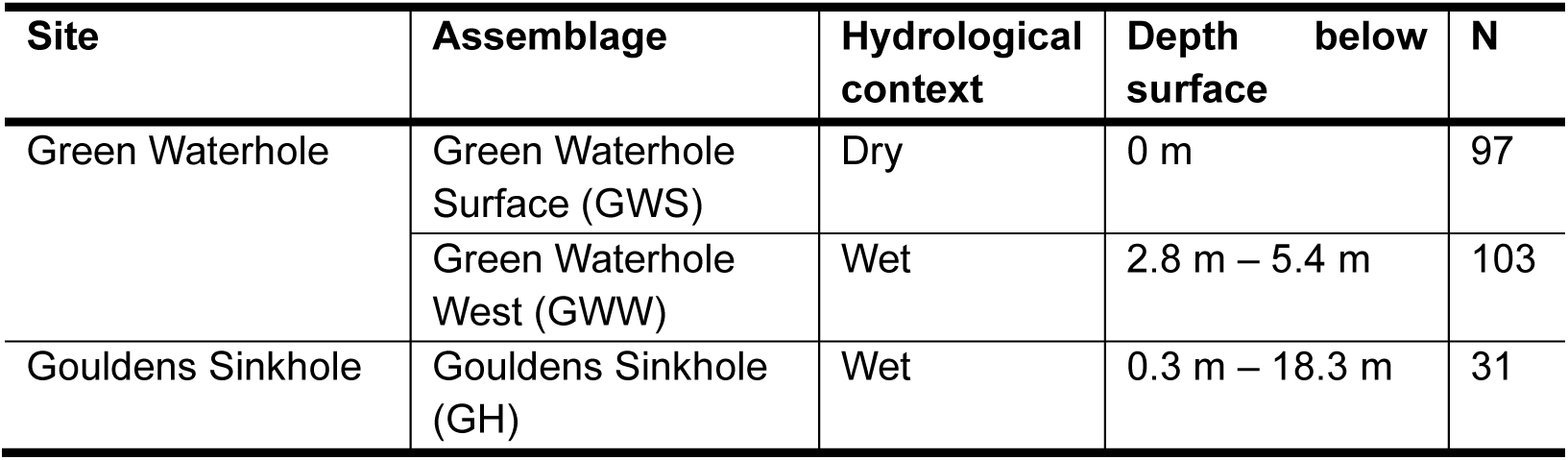
Site assemblages and associated hydrological context and specimen frequencies.

| Site | Assemblage | Hydrological context | Depth below surface | N |
| --- | --- | --- | --- | --- |
| Green Waterhole | Green Waterhole Surface (GWS) | Dry | 0 m | 97 |
|  | Green Waterhole West (GWW) | Wet | 2.8 m – 5.4 m | 103 |
| Gouldens Sinkhole | Gouldens Sinkhole (GH) | Wet | 0.3 m – 18.3 m | 31 |

At Green Waterhole, bones were collected from the submerged, wet ‘Green Waterhole West’ lake (GWW) and dry ‘Green Waterhole Surface’ (GWS) (Fig 2). The western lake (GWW) was targeted to minimise the effects of frequent diving activities, common in the eastern lake, and it was less disturbed by previous palaeontological research programs [4]. In the submerged lake, skeletal specimens were collected from five depths across either an allochthonous silty humic clay sediment near the entrance, or an autochthonous fine, powdery clayey carbonate sediment within the overhead cave environment [4]. Specimens collected from deeper regions of the cave experienced less light exposure than those higher and towards the entrance. The dry surface assemblage was collected across seventeen discrete locations from or on top of humic sediment. Seasonal water level fluctuations, and historically higher ground water levels than found presently suggest that bones collected from the perimeter of the doline at the water’s edge experienced wet conditions in the past [48]. Dumped refuse around the margins of the doline contained fragments of glass, plastic, PVC pipe, metal, and ceramics intermixed with bone.

At Gouldens Sinkhole (GH), bones were collected from nine locations on sediments resembling those at Green Waterhole (Fig 2). Specimens collected from within the sinkhole are presumed to have been underwater since deposition. Those found deeper, towards the back of the cave in the overhead environment, experienced decreased light exposure. One ‘dry’ specimen was collected from the artificially constructed ramp, emphasising its association with a historic or modern period. A specimen collected from a shallow underwater shelf (-0.3m) likely experienced seasonal hydrological flux. Historic refuse, including metal fragments and a dead tree, are spread across the sinkhole talus cone.

Bones from wet environments were conserved as much as possible during collection, transport, drying and cleaning processes. Waterlogged bones were transported in sealed and insulated containers and slowly dried out in laboratory conditions. When bones were completely dry, debris were removed using dry brushes to limit exfoliation of bone surfaces, and ethanol, acetone or water was applied to spot clean localised areas of bones where required. Delamination was monitored across the drying and cleaning processes.

Forty-two skeletal samples (Table 4 in S1 Appendix) were submitted for radiocarbon analysis to the Australian Nuclear Science and Technology Organisation (ANSTO) (49) (12 samples) and the Australian National University (ANU) Radiocarbon Laboratory [50, 51] (30 samples). Historic assemblages are rarely analysed because calibrated ages have large and/or multiple age ranges, due to rapid fluctuations in atmospheric radiocarbon content, in part due to the impact of industrialisation on the radiocarbon calibration curve (“the Suess effect”; [52, 53]). To offset this problem, this work identifies bones deposited across two time periods: decadal (<50 years) and centennial (>50, <185 years for domesticates and <1000 for native fauna). Each group was defined and examined to determine if the accumulation of taphonomic modifications on bone occurs at different temporal resolutions. Age groups are defined using historic documentation of European occupation starting in the Mount Gambier region from 1841 [54–56], and the period of intense atomic bomb testing that markedly increased atmospheric radiocarbon content [57, 58]. Radiocarbon methods are outlined in S1 Appendix.

### Skeletal analysis

Bones were taxonomically identified to the lowest taxonomic unit and size class [59]. Skeletal element, epiphyseal fusion, and side were recorded [60]. Fragmentation (breakage index) and completeness [18, 61], shape and size of specimens [60, 62, 63], and butchery and burning patterns and portions [64] were measured to assess site formation processes (details in S1 Appendix). Spatial analyses were limited to changes across depth. To quantify the assemblages [65, 66], the number of identifiable species (NISP), and minimum number of individuals (MNI) were calculated based on the zonation method proposed by Dobney and Rielly [67], and taking into consideration side, fusion stages, and refitting analysis. Fifteen specimens were selected for taxonomic identification through Zooarchaeology by Mass Spectrometry (ZooMS) analysis (details in S1 Appendix).

### Bone surface modifications

Bone modifications [21] were recorded as follows: Bone surface alterations included linear marks, pits and perforations, deposition of sediments, and discolouration and staining [21]; modifications to shape included scale of abrasion and rounding [68]; modifications associated with penetration into bone tissue included flaking and cracking, corrosion expression, digestion, and bone mineral modifications [18, 21]; and finally modifications relating to a removal or separation of bone tissue or skeletal elements included breakage, deformation, and crushing [69].

The presence/absence, description, and location of changes were recorded for taphonomic features that were then categorised into agent type (physical, chemical, biological). Quantitative methods were used to do this where possible (Table 2 in S1 Appendix). For physical abrasion features, we included rounding, polishing, scratches and scuff marks, crushing, collection damage, and impacts. For physical distortion and deformation features, we included periosteal bone shrinkage, delamination, plastic deformation, and distortion. Chemical corrosion included the degree of damage (general, >50% of the specimen, or isolated), expression (pitting, bone loss, surface), depth of penetration (superficial or deep), and spread type (continuous or discontinuous). In instances of chemical mineral modification, we used chalky texture, mineralisation, cementation of sediments, chalky mineral deposition [4, 7]. The measure of mineralisation is broad, and determined by weight, colour and texture changes. Biological anthropic changes included burning, butchery marks, and percussion marks, and for fauna and flora, root pitting, floral/biological etching, insect boring, fungal/agal growth, gnawing, puncture marks (location, opposing marks, shape), and animal scratch marks. Except for rodent gnawing and predation marks, biological agents were only recorded where they were observed to directly create a modification. See S1 Appendix for further information.

**Table 2.** Histological definitions previously associated with aquatic biodegegredation.

| Aquatic feature | Definition | Reference |
| --- | --- | --- |
| Wedl-type 1 or cyanobacterial tunnelling | Random, singular or bifurcating tunnels between 5 – 19 μm, without a hypermineralised rim. Extending up to 200 - 300 μm from the peripheral surface. Round grains may be identified at the end of tunnel structures. | [35, 76-80] |
| Lacustrine microboring | See Wedl-type 1, between 7 - 18 μm in diameter with a hypermineralised boundary viewed under bSEM | [76, 81, 82] |
| Radial microfractures | Small radial fractures, spanning across the secondary osteon cement line at a perpendicular angle. | [36, 83] |

Staining was identified by the colour, location (general or isolated), margin type (sharp or diffuse), and spread (mottled or continuous). Non-invasive elemental composition of black staining was conducted using portable X-ray florescence (pXRF) [70, 71] (details in S1 Appendix).

Weathering stages were recorded alongside the presence and absence of each feature (shallow split lines, deep longitudinal cracking, bleaching, flaking) associated with its scale to determine potential intricacies of ‘weathering’ across dry and wet environments [72]. Flaking was distinguished from delamination. Delamination is the separation of a single external bone layer (approximately >1mm thick) beginning with initial separation of periosteal lamellate from the bone cortex, followed by longitudinal cracking, and finally removal of bone. Each stage was included in the definition of delamination. Flaking and exfoliation ranged between the light, superficial removal of bone surfaces in either continuous or irregular patches not defined by bone structural orientation, to removal of bone surfaces first preceded by linear cracking as observed during the weathering process. Flaking produces multiple layers of flakes, that penetrate the bone matrix at different depths compared to the single delamination event.

### Histological preparation and analysis

Twenty-four samples were selected for histological analysis (GH N=5, GWW N=8, GWS N=10). Weightbearing long bones and ribs were selected to target Haversian bone systems that produce secondary osteons (Table 4 in S1 Appendix), following taxonomic identification (see further below). Representative samples from large (e.g. kangaroo and cow) and medium (e.g. sheep) animals were selected for each context where possible based on zooarchaeological classifications [59]. Standard methods for undecalcified, unstained bone histology preparation were followed to create specimen blocks for scanning electron microscopy (SEM) analysis, and thin sections for histological analysis [34, 73] (details in S1 Appendix). Blocks at least 3 mm thick, and thin sections of approximate 100 µm thickness, were examined for markers of bioerosion, and radial microfractures across secondary osteons under backscatter SEM and transmitted light microscopy, respectively, to identify size, form, demineralisation, hypermineraliation, and potential associations with histological features [74]. Most blocks and thin sections represented a complete cross-section (n=21) through a bone shaft, unless only a portion was available (n=3). The location of biodegradation was identified as either peripheral modification (localised or more regional at the outer bone pocket within a block/thin section) or general destruction (widespread impact across the block/think section). To quantify the peripheral degradation, a maximum depth of penetration was measured starting at the periosteal border using the straight-line tool in ImageJ 1.54g.

The type of bioerosion marker observed was grouped into either Wedl-Tunnelling Type 1 (possible associated with cyanobacterial attack) or other Microscopic Focal Destruction (MFD) [32, 35, 75, 76]. Inconsistent use of terminology in the histotaphonomy literature presents issues in evaluating features associated with aquatic environments, with some authors identifying cyanobacterial tunnelling as synonymous with Wedl-tunnelling Type 1 [35, 76], and others separating them into a unique form [29]. For simplicity, cyanobacteria are here identified as Wedl-tunnelling type 1 based on original descriptions (Table 2).

Quantitative measures of the Oxford Histological Index (OHI), bacterial attack (Bacterial-Attack-Index; BAI) and cyanobacterial attack (Cyanobacterial-Attack-Index; CAI) were measured using a scale of 0 to 5 [29, 84], with 0 indicating less than 5% preservation and 5 indicating over 95% preservation (Table 3 in S1 Appendix). All samples were also assessed for birefringence under polarised light. Finally, we measured total cortical area, and the area of regions modified by the different biodegradation markers using the polygon tool in ImageJ 1.54g. Percentages were then calculated for each biodegradation type to estimate OHI, BAI and CAI from total cross sections.

**Table 3.** Frequency of specimens associated with age periods across sites and burial conditions.

| Site | Condition | Age | Age Period | n= |
| --- | --- | --- | --- | --- |
| Gouldens Sinkhole | Wet | >1955 | Decadal | 0 |
|  | Wet | 1841 - 1955 | Centennial | 13 |
| Green Waterhole West | Wet | >1955 | Decadal | 9 |
|  | Wet | 1841 - 1955 | Centennial | 1 |
| Green Waterhole Surface | Dry | >1955 | Decadal | 1 |
|  | Dry | 1841 - 1955 | Centennial | 12 |
| Total | Wet | >1955 | Decadal | 9 |
|  | Wet | 1841 - 1955 | Centennial | 15 |
|  | Dry | >1955 | Decadal | 1 |
|  | Dry | 1841 - 1955 | Centennial | 12 |

**Table 4.** Number of identified species (NISP) and minimum number of individuals (MNI) for taxon across the wet and dry assemblages.

|  |  | GH (wet) |  | GWW (wet) |  | GWS (dry) |  |
| --- | --- | --- | --- | --- | --- | --- | --- |
| Taxa |  | NISP | MNI | NISP | MNI | NISP | MNI |
| Introduced | *Bos taurus* | - | - | 4 | 1 | - | - |
|  | *Bos* sp. | 1 | 1 | 41 | 2 | 2 | 1 |
|  | cf. *Bos* sp. | 3 | 1 | 12 | 1 | 2 | 1 |
|  | *Ovis aries* | 1 | 1 | - | - | 4 | 2 |
|  | Ovicaprid | 6 | 3 | 20 | 3 | 55 | 2 |
|  | cf. ovicaprid | - | - | 1 | 1 | - | - |
|  | *Sus domesticus* | - | - | 3 | 1 | - | - |
|  | *Sus* sp. | - | - | 7 | 1 | - | - |
|  | *Oryctolagus* | - | - | - | - | 12 | 2 |
| Native | Macropodidae | 3 | 1 | - | - | - | - |
|  | Macropodinae | - | - | 1 | - | - | - |
|  | *Macropus* sp. | 6 | 1 | 1 | - | - | - |
|  | cf. *Macropus* | 2 | 1 | 6 | - | 1 | 1 |
|  | cf. *Notamacropus* | - | - | - | - | 1 | 1 |
|  | *Dromaius novaehollandiae* | - | - | 1 | 1 | - | - |
|  | *Canis lupus dingo* | 2 | 1 | - | - | - | - |
|  | *Dasyurus* sp. |  | - | - | - | 2 | 1 |
|  | *Trichosurus vulpecula* | 2 | 1 | - | - | - | - |
|  | *Rattus* sp. cf. *R. lutreolus* | 1 | 1 | - | - | - | - |
|  | Unidentified | 4 | - | 6 | - | 17 | - |
| Total |  | 31 | - | 103 | - | 97 | - |

### Statistical analyses

Statistical comparisons between wet and dry conditions were undertaken using IBMM SPSS 29 with significance tested at 95% and 99% confidence intervals. Differences between the presence and absence of bone modifications and histological data were tested using the 2-sided asymptotic Pearson chi-squared analysis. Although all sample sizes were adequate across all variables (>100), some variables presented in groups at low frequencies (<6), and in these instances, Fisher’s Exact 2-sided test was performed. We acknowledge that larger sample sizes will be necessary to validate our findings in these cases. Ordinal bone surface and histological data were tested across burial conditions using the non-parametric, independent samples Mann-Whitney U test to identify significant differences in distributions. The OHI, BAI and CAI scales, and peripheral degradation were further compared across depth in aquatic contexts. Temporal categories were used to test statistical differences in taphonomic indicators across the wet assemblage through time, and for the total pooled wet and dry assemblage to understand if differences are a result of time or burial condition.

## Results

### Site chronologies

Of the 42 specimens submitted for radiometric carbon dating, 41 passed pretreatment screening (S1 Table). Two groups were identified associated with domesticates: decadal, representing the modern period younger than 1955; and centennial, representing deposition between 1841 and 1955 (Table 3). Analysis of bone modifications through time can only be conducted across wet assemblages as too few decadal specimens are represented from dry conditions (n=1).

### Skeletal assemblage

A total of 231 bone specimens were analysed from GWW (NISP=103), GWS (NISP=97), and GH (NISP=31) (Table 4). Large taxa include *Bos taurus* (cow), Macropodinae and *Macropus* (kangaroo), *Dromaius novaehollandiae* (emu); medium taxa: ovicaprid (sheep/goat), *Sus scrofa* (pig), *Canis lupus* (dog/dingo); and small taxa: *Oryctolagus* (rabbit), *Trichosurus vulpecula* (possum), *Dasyurus* (quoll) and *Rattus* sp. cf. *R. lutreolus* (swamp rat). All specimens analysed by ZooMS were identified to the genus or species level: *Macropus* sp. (n=3), *Canis* sp. (n=1), *B. taurus* (n=3), *O. aeries* (n=7), and *S. scrofa* (n=1) (S1 Table). Ancient DNA techniques identified decadal sheep from the underwater assemblages as belonging to the Merino breed [85].

For domesticates, 70.4% of bones are associated with the forelimb and hindlimb (GH= 45.5%, GWW=68.2%, GWS=77.8%), followed by axial elements (rib/vertebrate: GH= 45.5%, GWW=19.3%, GWS=17.5%) and cranial elements (GH= 9.2%, GWW=11.4%, GWS=4.8%).

Only a single *in situ* articulation was identified across the three sites, a rabbit skeleton consisting of 12 elements from GWS, located on the surface near the margins of the lake. It is possible that other articulated elements were present but missed due to limited visibility underwater during recovery. Specimens from underwater contexts have likely been reworked by divers prior to collection.

#### Quantitative distributions

Bone fragments were generally large across sites, with the highest proportion falling into size class 6 (32 - 128mm: GH=71.4%, GWW=36.8%, GWS=48.5%). Specimens from GWW trended larger than those from GWS; 84.5% of the GWS assemblage fall between size classes 5 and 6 and 90.8% of bones at GWW fall between size classes 6 and 8. No identifiable size class sorting by depth was observed across samples from wet contexts, or by site across dry burial sites (S1 Fig).

Bone specimen shape (maximum breadth and length ratio) indicated bone sorting based on depth, where an increase in depth was associated with a more even shape (Fig 3). A significant correlation between the breadth to length ratio and depth was identified across both wet sites; however, the correlation coefficient at GH is higher (R=0.730, *p*=0.007) than GWW (R=0.311, *p*=0.005), suggesting depth is a better predictor of shape of the bone at the former site.

**Fig 3.**
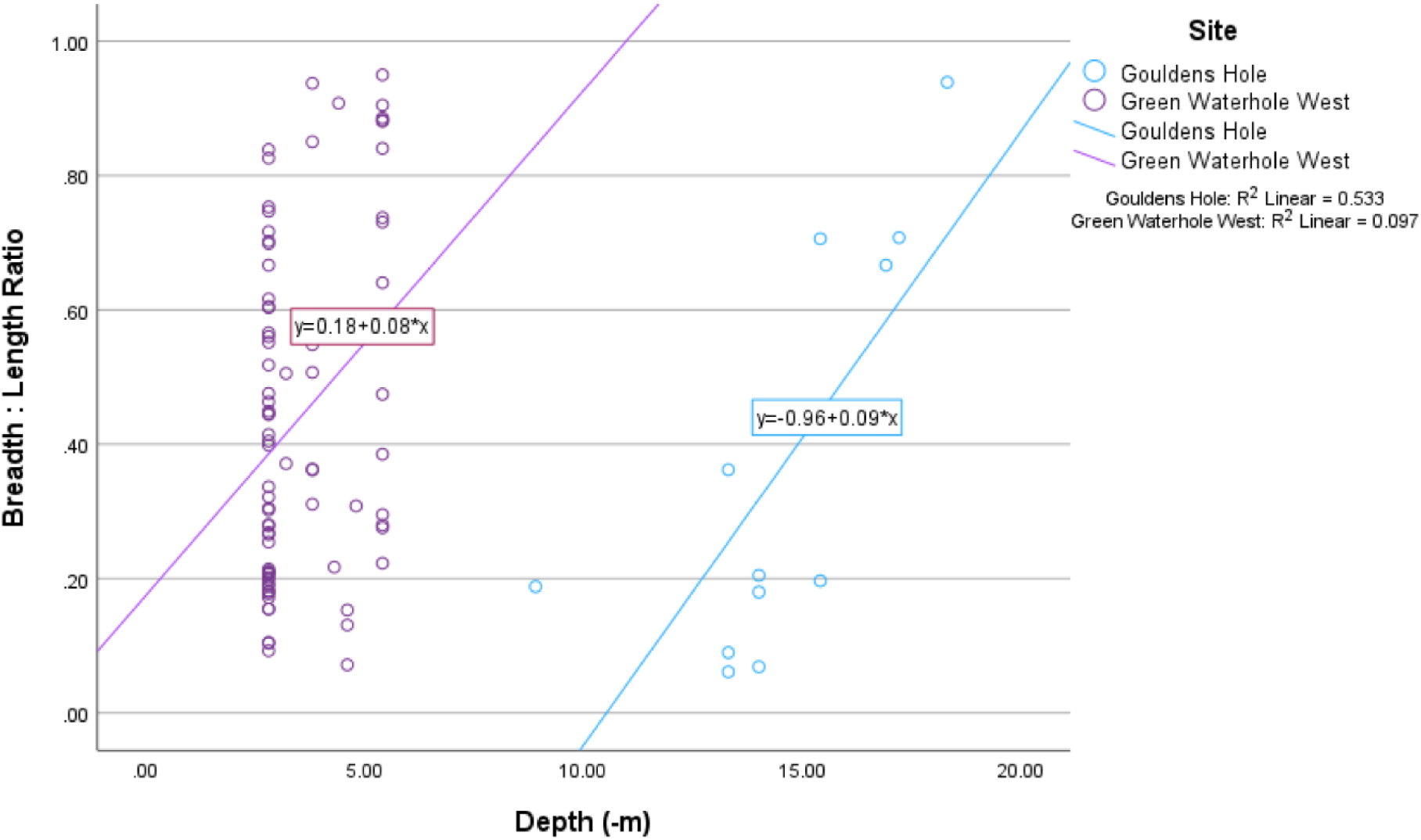
Shape variation (60) across dry (0m) and wet (-0.3 to -18.3m) contexts.

### Bone Surface modifications

Bone surface modifications (BSMs) were identified across wet (GH and GWW) and dry (GWS) burial environments (Table 5). Data are presented according to associated agent types: physical, chemical and biological. Raw frequency BSM data across wet and dry contexts are provided in S1 Dataset.

**Table 5.**
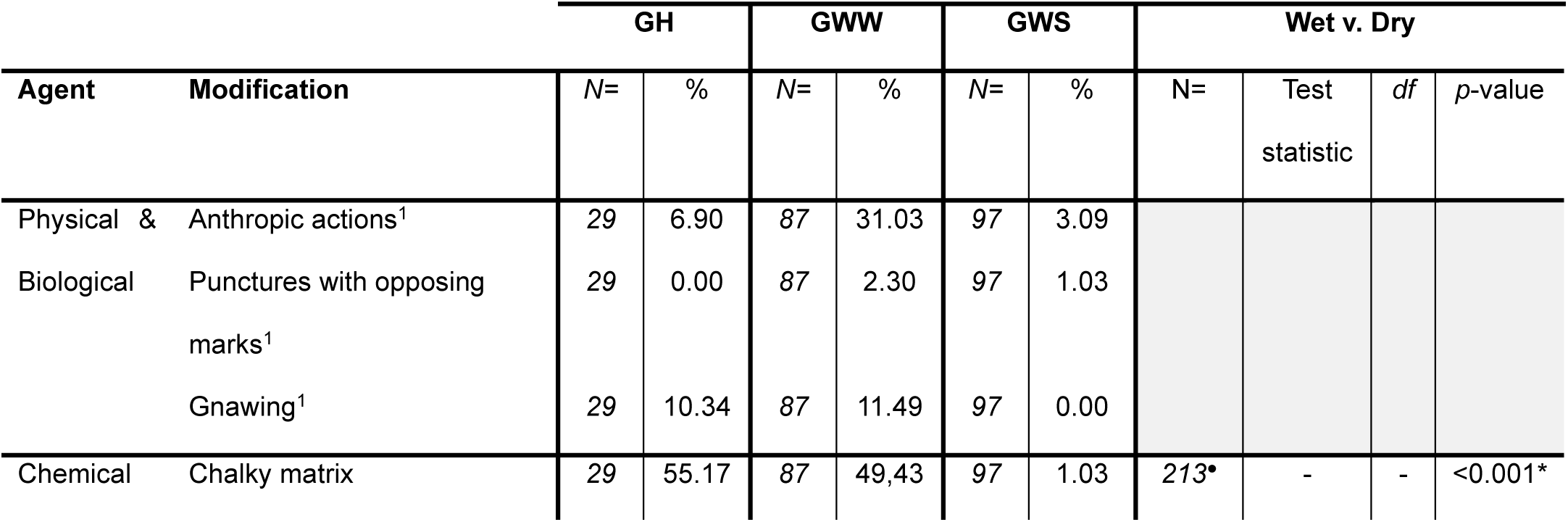
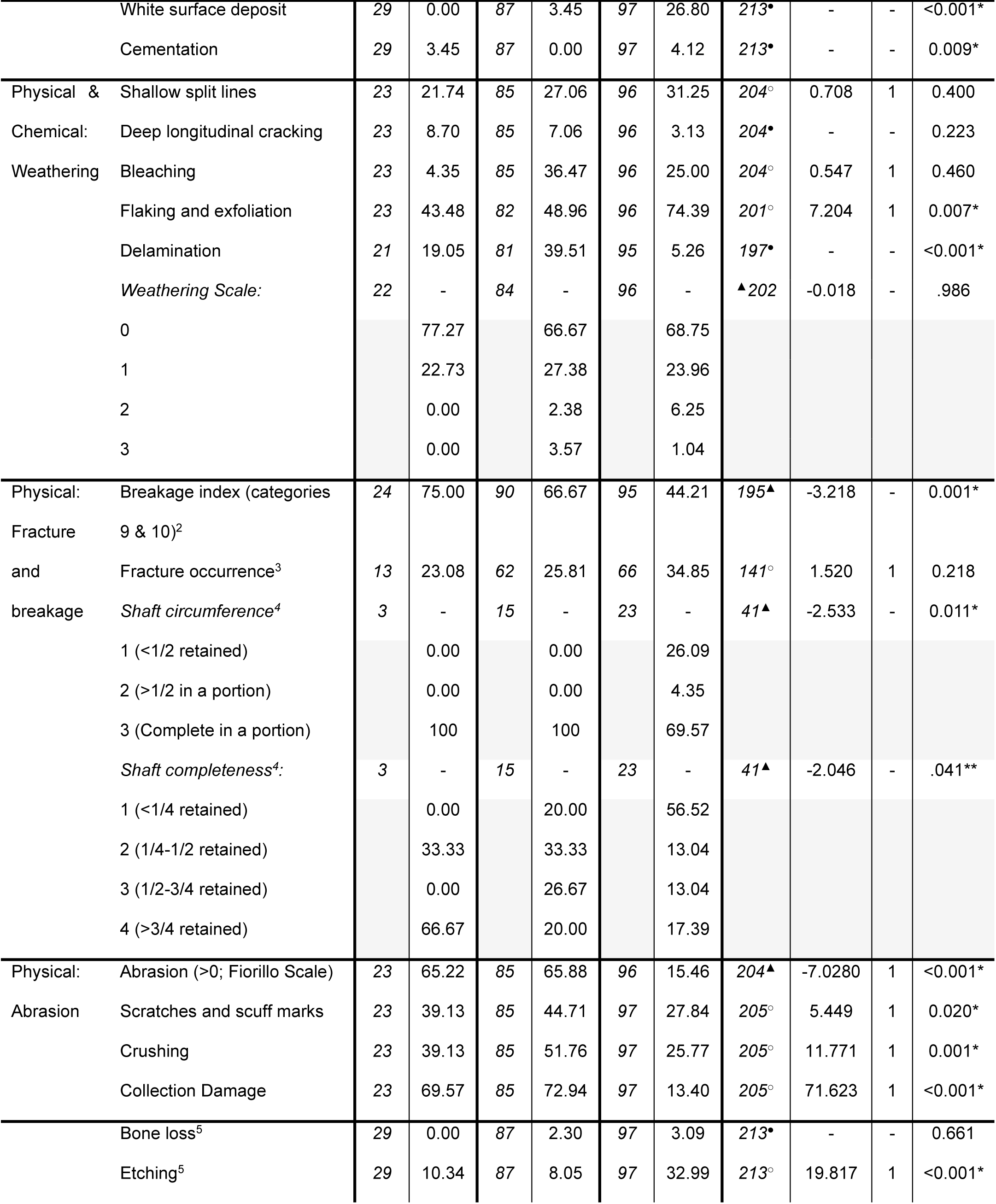
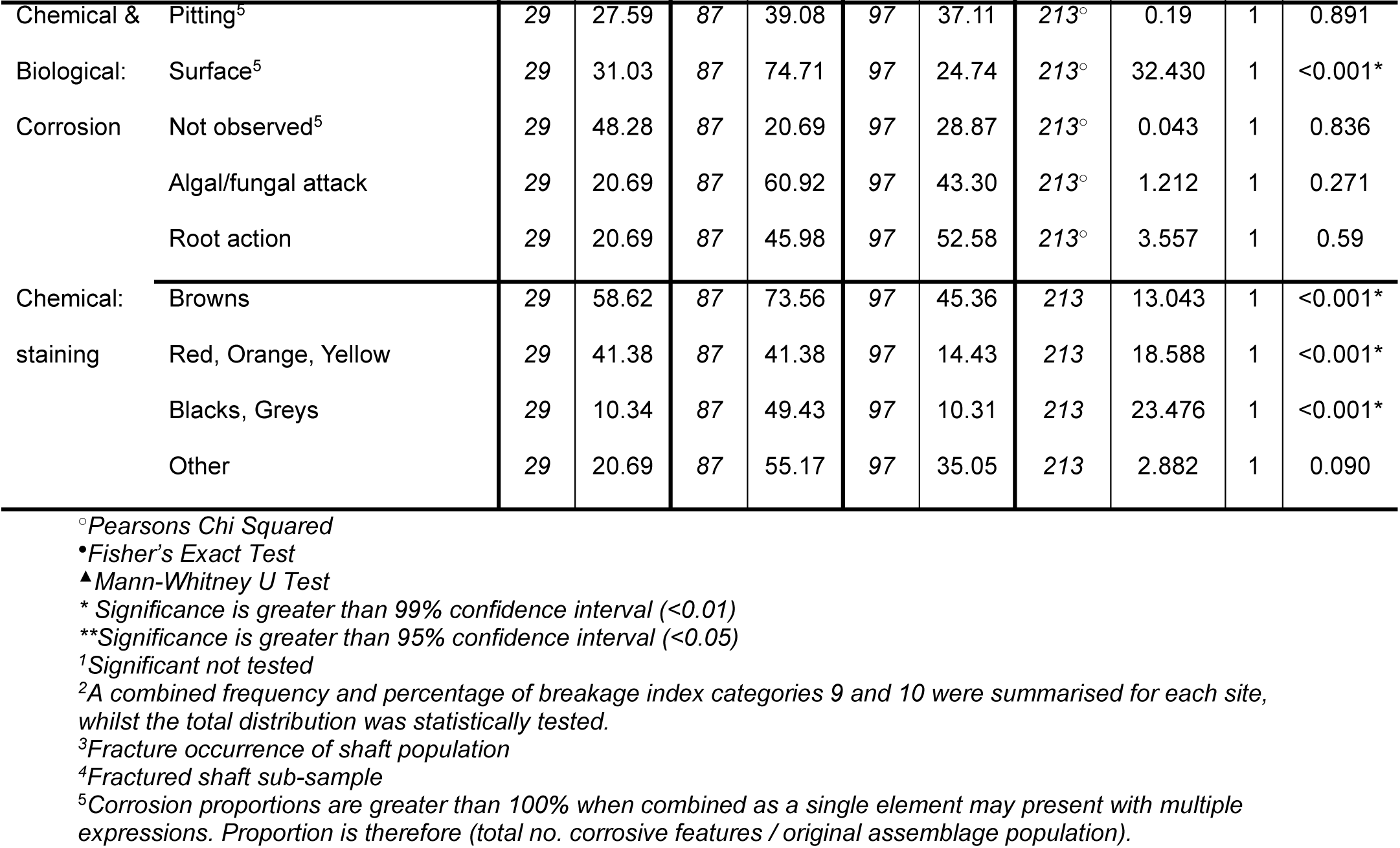
Summary of bone surface modifications across Gouldens Sinkhole (GH), Green Waterhole West (GWW), and Green Waterhole Surface (GWS) assemblages. Columns reflect the total (N=) number of specimens analysed from the assemblage, and the relative proportion (%) of the assemblage to present with that modification. Number of specimens in the assemblage for each modification may vary depending on availability of data.

#### Anthropic actions and predation

Anthropic modifications (n=33) were only identified in domesticated taxa. A single specimen from GWS showed evidence of burning (Stage 2) but did not exhibit cut marks. Butchery marks (n=32) include saw marks across full cross sections, sawn bone, and thin linear cuts with V-shaped cross sections (Fig 4a-b). Modified elements included ribs, scapulae, innominate, femora and vertebrae. One bone presented with thick abraded lines and crushed margins across the cortex, identified as the effect of rubbing against cave diving line (2 - 6mm braided nylon line) when used a secondary tie-off point (Fig 4c).

**Fig 4.**
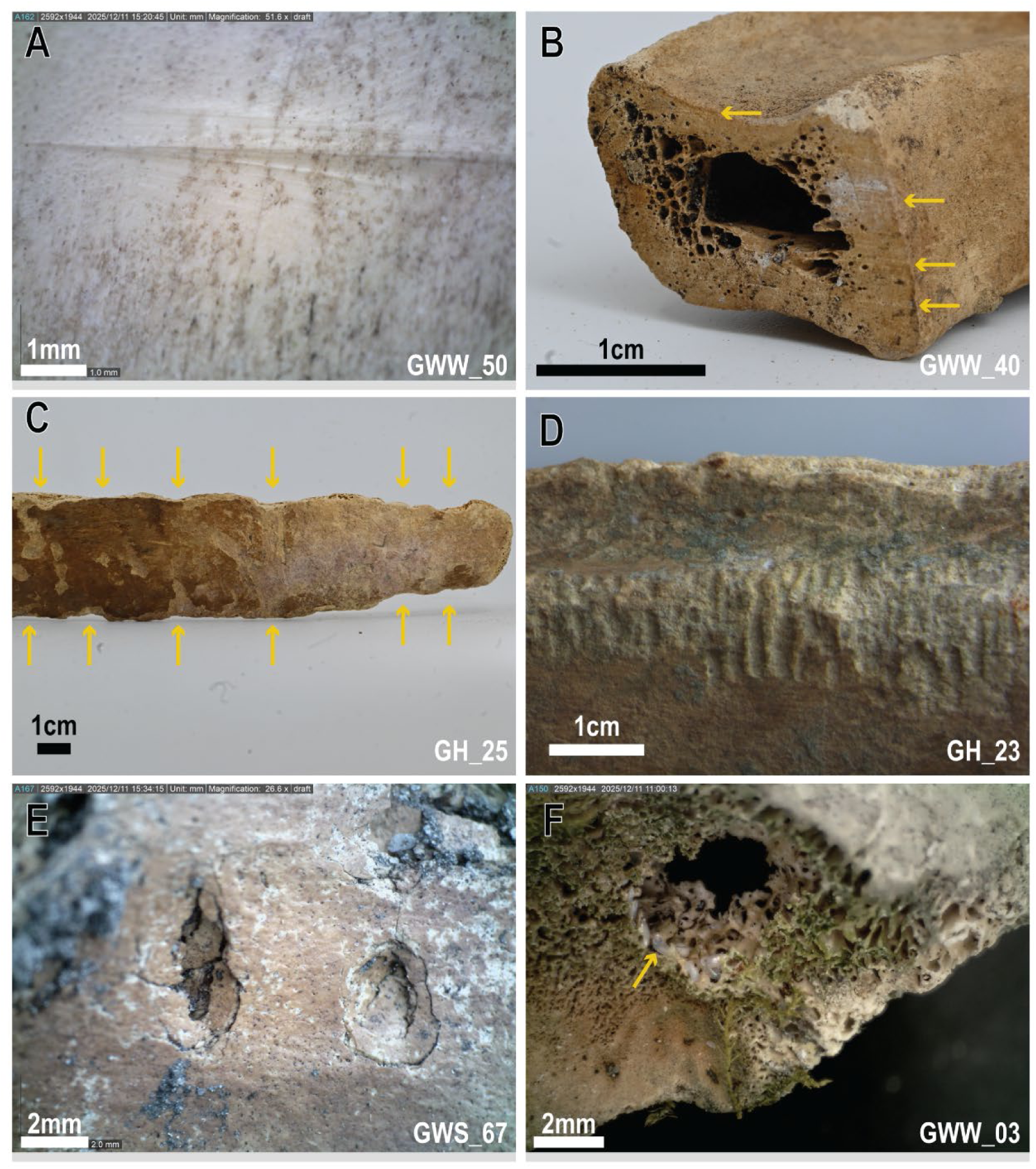
Anthropic actions and predation across wet (A-D, F) and dry (E) cave assemblages. A: Slice marks on cow humerus from knife with V-shaped profile; B: cow ventral rib end sawn through, yellow arrows indicate examples of saw striations; C: cow distal rib deformed by cave line (2-6mm braided nylon line) on caudal and cranial margins indicated by paired yellow arrows; D: gnawing across cow rib margin; E: paired predation puncture marks with conical profile; F: insect boring through trabecular bone and embedded ant eggs.

Low frequencies of gnawing marks were identified across GH (n=3) and GWW (n=10), no gnawing was identified from GWS (Table 5; Fig 4d). Carnivore predation, specifically paired dental punctures with opposing marks, were observed in low frequencies in wet and dry conditions (GWW n=2, GWS n=1, Fig 4e). Possible insect boring was observed on four bones from underwater caves (GWW n=3, GH n=1, Fig 4f).

#### Bone quality and cementation

All but three samples from GWW were unmineralised, a generalisation based on weight and texture of the element. Compared to wet burial sites, dry contexts were defined by a significant increase in isolated expressions of cementation, and white chalky mineral deposits expressed as both isolated specks and broad coverage (Table 5). A significant proportion of specimens recovered from underwater contexts exhibited an increase in chalky texture (Table 5, Fig 5A-B). The chalky texture is limited to the sub-periosteal pocket, sandwiched between a thin solid exterior bone and the hard cortical bone beneath (Fig 4B). It was only observed when bone surfaces had been modified to expose underlying layers, and thus we likely underestimate the frequency of this feature. Time was not found to influence the deposition of white mineral, or alteration of the bone matrix (Table 6).

**Fig 5.**
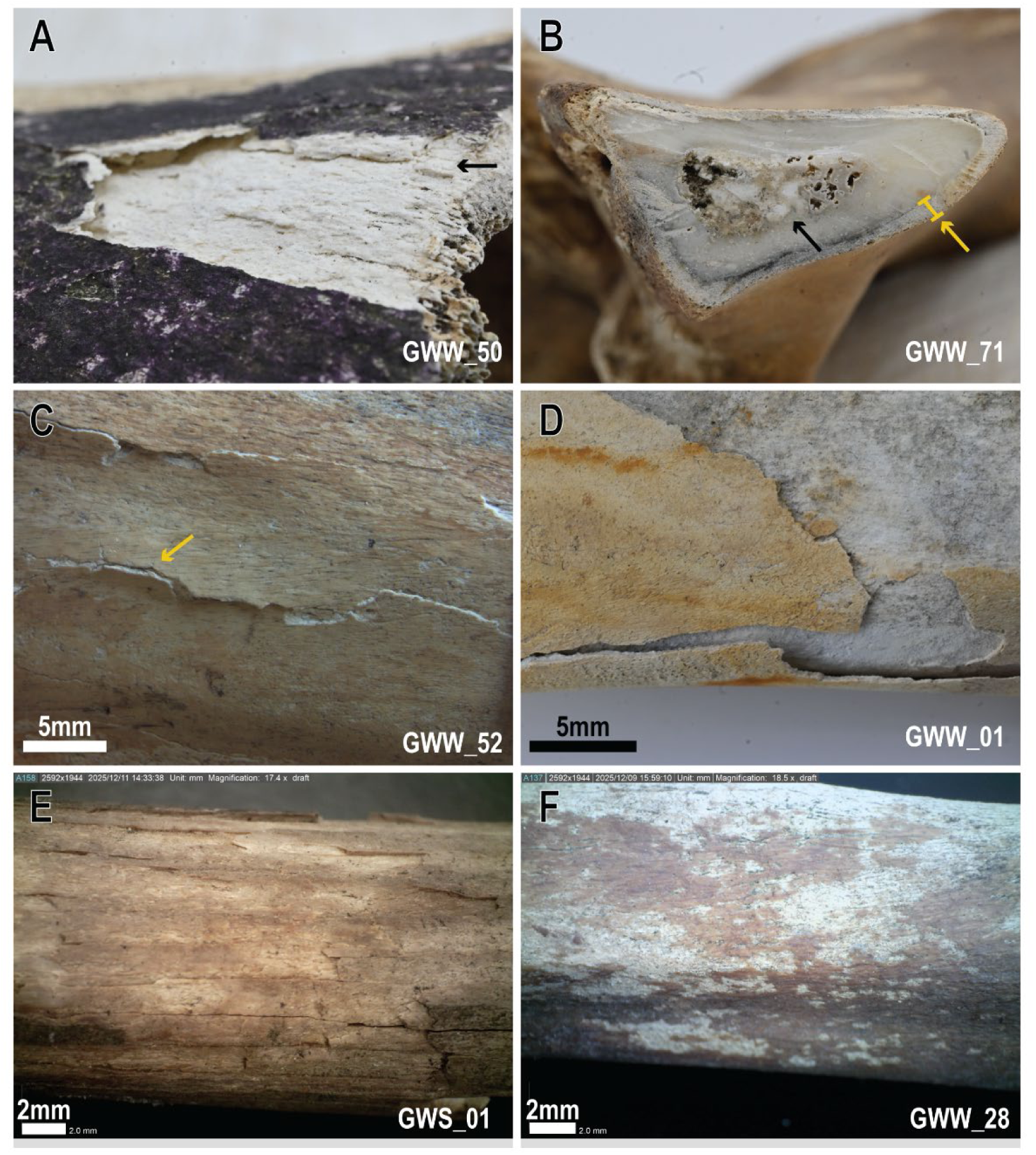
Bone quality and weathering modifications. A: chalky white bone surface (black arrow) exposed through delamination and beneath biotic staining and corrosion; B: subperiosteal pocket of chalky bone modification (yellow indicators) beneath an intact bone surface, and preserved bone marrow (black arrow); C: early stages of delamination with bone surface layers separated but not removed, and ‘popped’ fracture margins (yellow arrow); D: late stages of delamination with bone separated surface; E: weathering flaking with irregular, multi-layered flakes removed from surface; F: the chemical process of desquamation/exfoliation resulting in the light removal of surface bone tissue.

**Fig 6.**
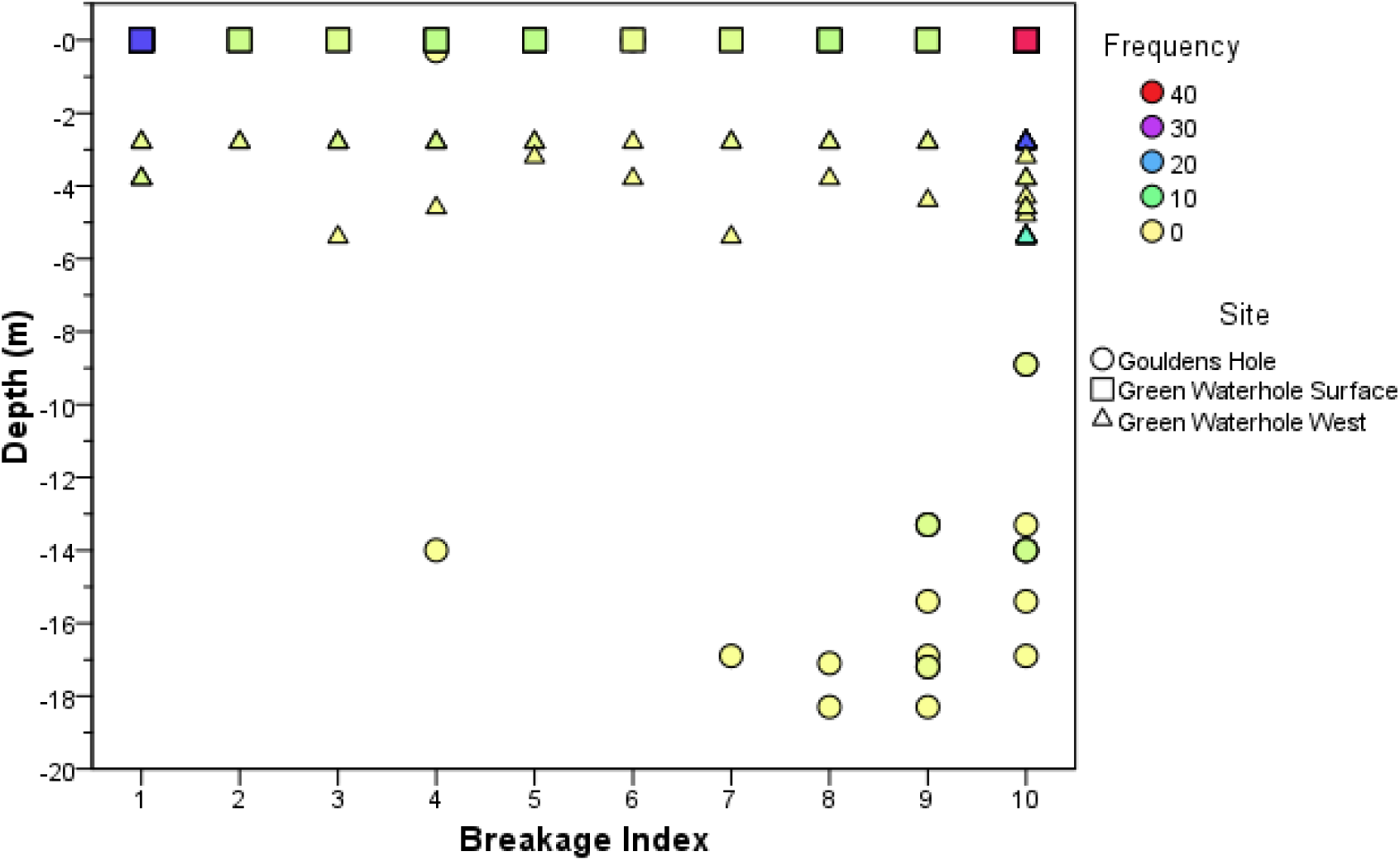
Distribution of Breakage Index (BI) frequencies across collection depth below water surface (m), grouped by site.

**Table 6.**
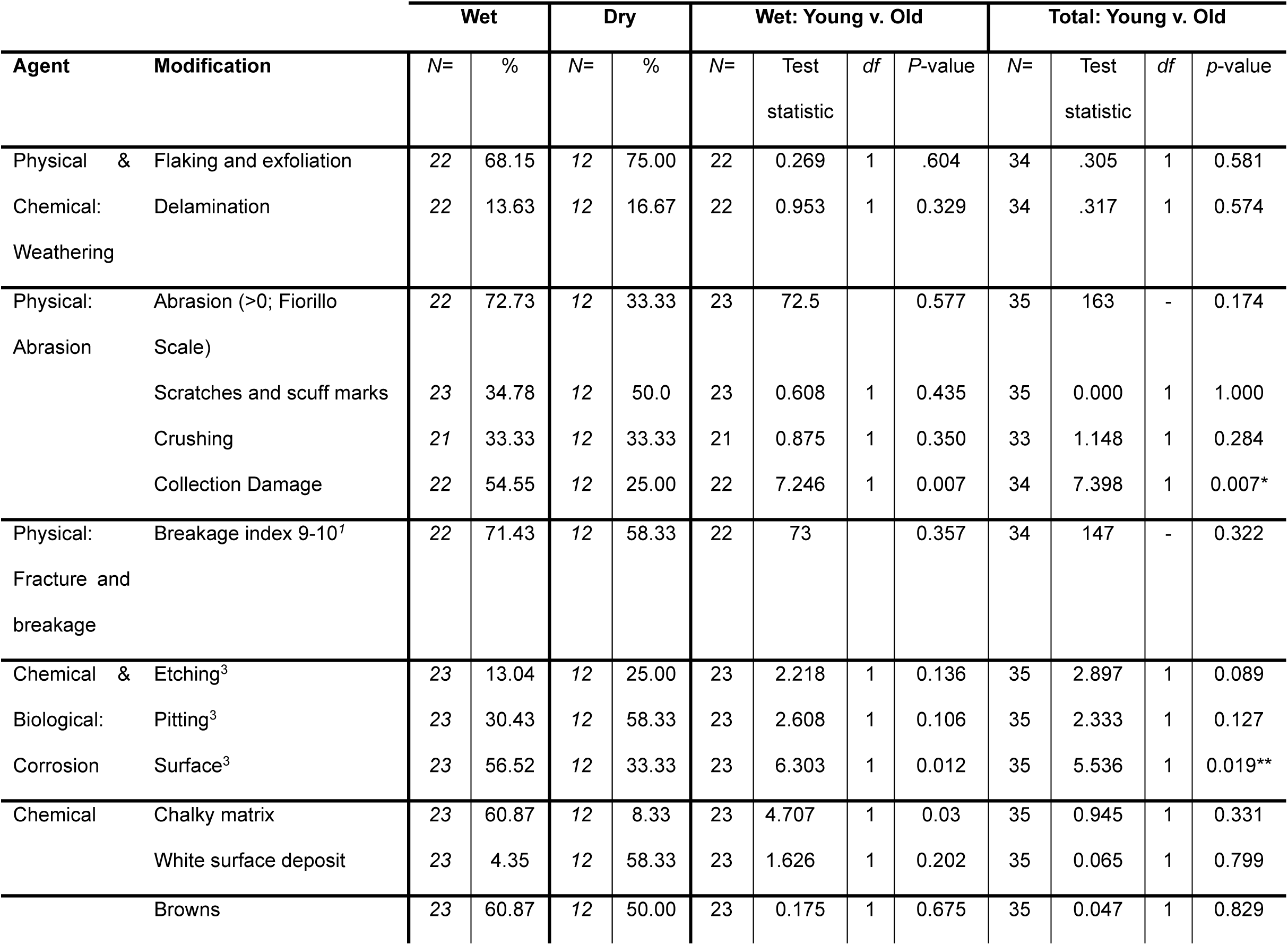
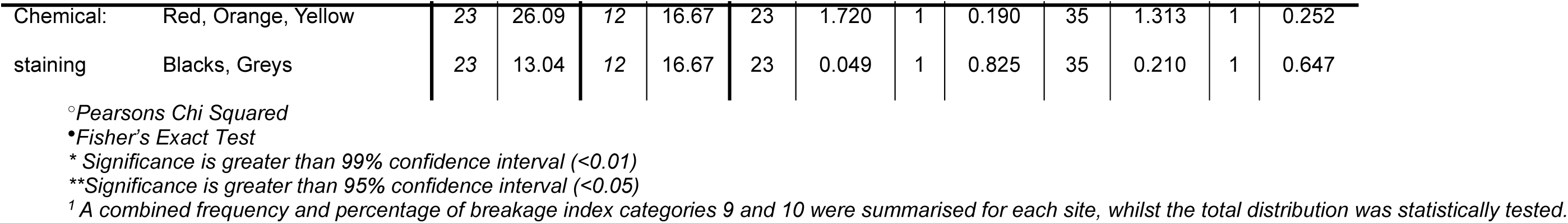
Chronological analysis of statistically significant features identified across the wet and dry assemblages.

#### Weathering

Weathering across the three assemblages was low with all specimens exhibiting weathering stage 3 and below (Table 5). Although no differences were identified between weathering stages across wet and dry environments, components of the weathering scale varied (Table 5). Wet bone presented significantly lower proportions of flaking and exfoliation (Fig 4E-F), and more bone surface delamination (Table 5, Fig 4C-D). Differences in weathering sub-categories across wet bones were not influenced by timescales (Table 6).

Despite controlling the drying process, delamination occurred post-collection and highlights the damage caused by wetting then drying (Fig 4C-D). Whilst delamination on bone from dry assemblages occurred solely on long bones of medium sized animals, all bone types recovered from submerged sites were impacted (Long n=26, 38.8%; Flat: n=5, 35.7%; Short: n=1, 50.0%; Irregular: n=14, 23.5%). In wet contexts, delamination was most frequent across bone shafts, areas not associated with spongiform bone structures (e.g. epiphyses). Large animal bones were more likely to be modified in wet assemblages (47.5%) followed by medium sized animals (20.7%). Too few small animals were recovered for calculations of proportion of bones with delaminated surfaces.

#### Physical agents

Breakage Index (BI) was significantly different between wet and dry assemblages, but with low levels of breakage observed across the three sites (Table 5). Bones either recorded high (BI 9 &10) or low (BI 1) levels of completeness, but submerged assemblages recorded the highest rates of completeness (Table 5). The proportion of bones with very high completeness at GWW increases when anthropogenic butchery fragmentation is excluded (69.1%). Breakage was not impacted by depth or time since deposition (Table 6, Fig 5).

Fracture frequencies were consistent across wet and dry contexts, however wet assemblages exhibited significantly greater preservation of long bone shaft circumferences, and higher levels of shaft completeness (Table 5). Over half of the fractured specimens at GWS retained less than a quarter of their original shaft whilst specimens from underwater contexts showed variable shaft fragmentation (Table 5). Fracture pattern frequencies in wet contexts were not tested for differences across time scales due to small sample size (decadal n=1).

Saturated wet bones were soft, both before and after drying, and prone to damage, such as plastic deformation, during collection, transport, and handling (Table 5). This influenced data attributed to physical modifications by adding bone surface modifications or removing evidence of past events. After excluding post-collection bone surface modifications, a single specimen at GH presented with plastic deformation at -14 metres below the water surface, whereas thirteen specimens were impacted at GWW across various depths (-2.8m n=14, -3.2m n=1, -4.3m n=1, -5.4m n=3). Physical damage was more common across the wet assemblages, and when compared to dry bones, presented with significantly greater levels of general abrasion, scratches and scuff marks, and crushing (Table 5). Collection damage could not be excluded from this data due to similarities between pre- and post-collection events. No bones were identified as rounded or polished because of physical, non-anthropic agents in each burial context. However, time significantly contributed to the likelihood that bones may be modified for the wet and pooled total assemblage (Table 6).

#### Corrosive and floral agents

High levels of pitting, general surface corrosion, forms of etching, and bone loss were identified across the assemblages (GH= 62.5%, GWW=79.3%, GWS=71.1%, frequently associated with an observed biological agent: flora and/or biofilms (Fig 7). Seventy specimens presented with more than one type of corrosion modification (GH n=5, GWW n=39, GWS n=26).

**Fig 7.**
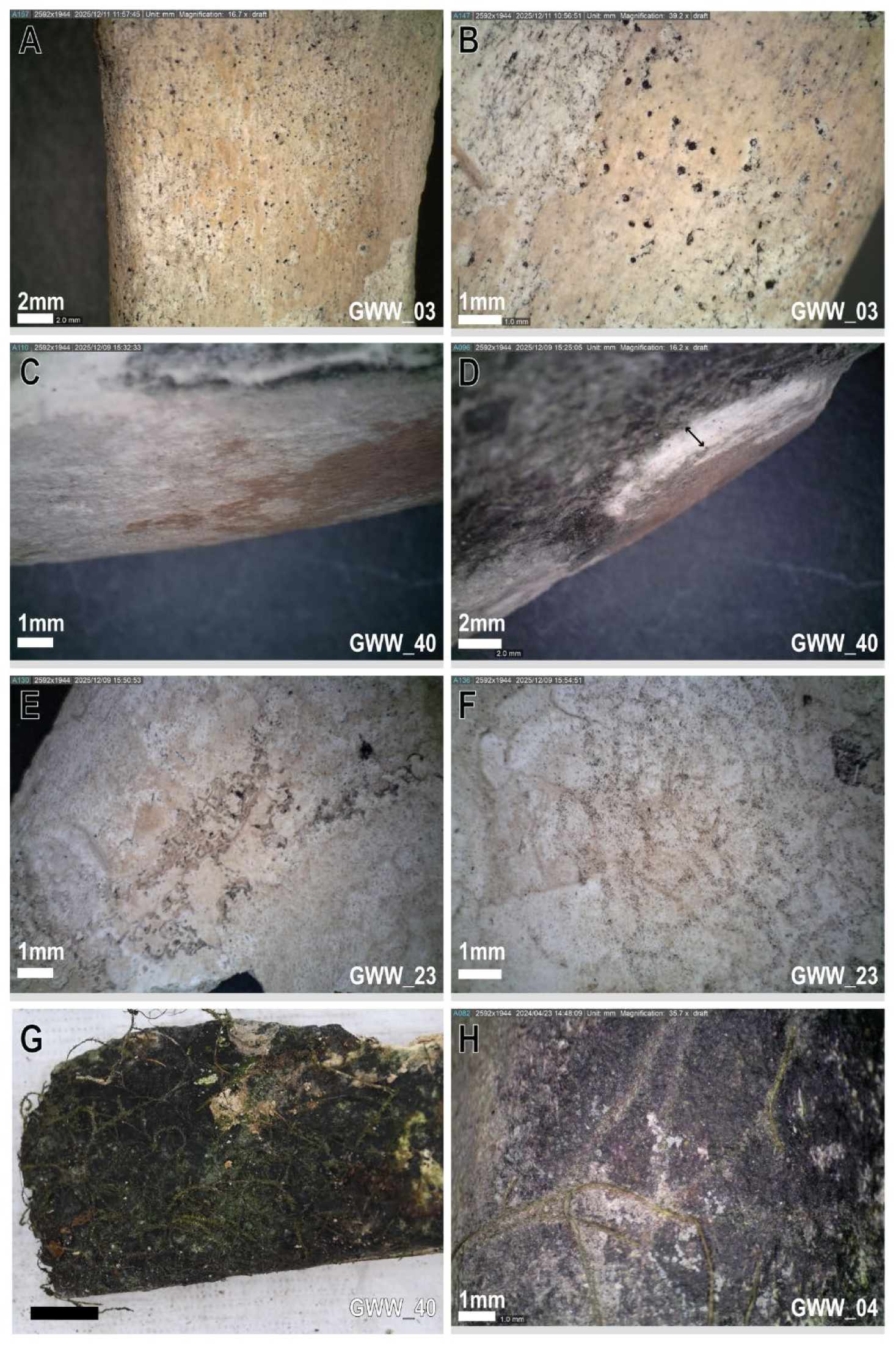
Corrosive features across wet environments. A-B: broad pitting across bone adjacent to surface removal (B); C: shallow, continuous surface corrosion; D: deep continuous surface corrosion with depth indicated by arrow; E-F: localised (E) and broad (F) continuous etching; G-H: broad surface corrosion with plant root attachments before cleaning (G) and a close up after cleaning (H).

Whilst the presence of etching was significantly more prominent across dry environments than wet (*p*<0.001), a distinct circular etching was identified as unique to the wet landscape, specifically at GWW (n=5, 4.1%; Fig 7). These are characterised by a superficial surface expression of corrosion featuring concentric rings (Fig 8), measuring approximately 5mm in diameter, complete circles or semicircular, and feature either a single or double ring. Isolated and clustered patterns of the circular target etching were observed. No identifiable agent was associated with these modifications. Linear etching was identified on bone from wet and dry contexts (GH= 10.3%, GWW=3.5%, GWS=29.9%). Flora was not identified alongside all etching, and some bones with adhering flora did not express linear etching, indicating a degree of attack that warrants further investigation.

**Fig 8.**
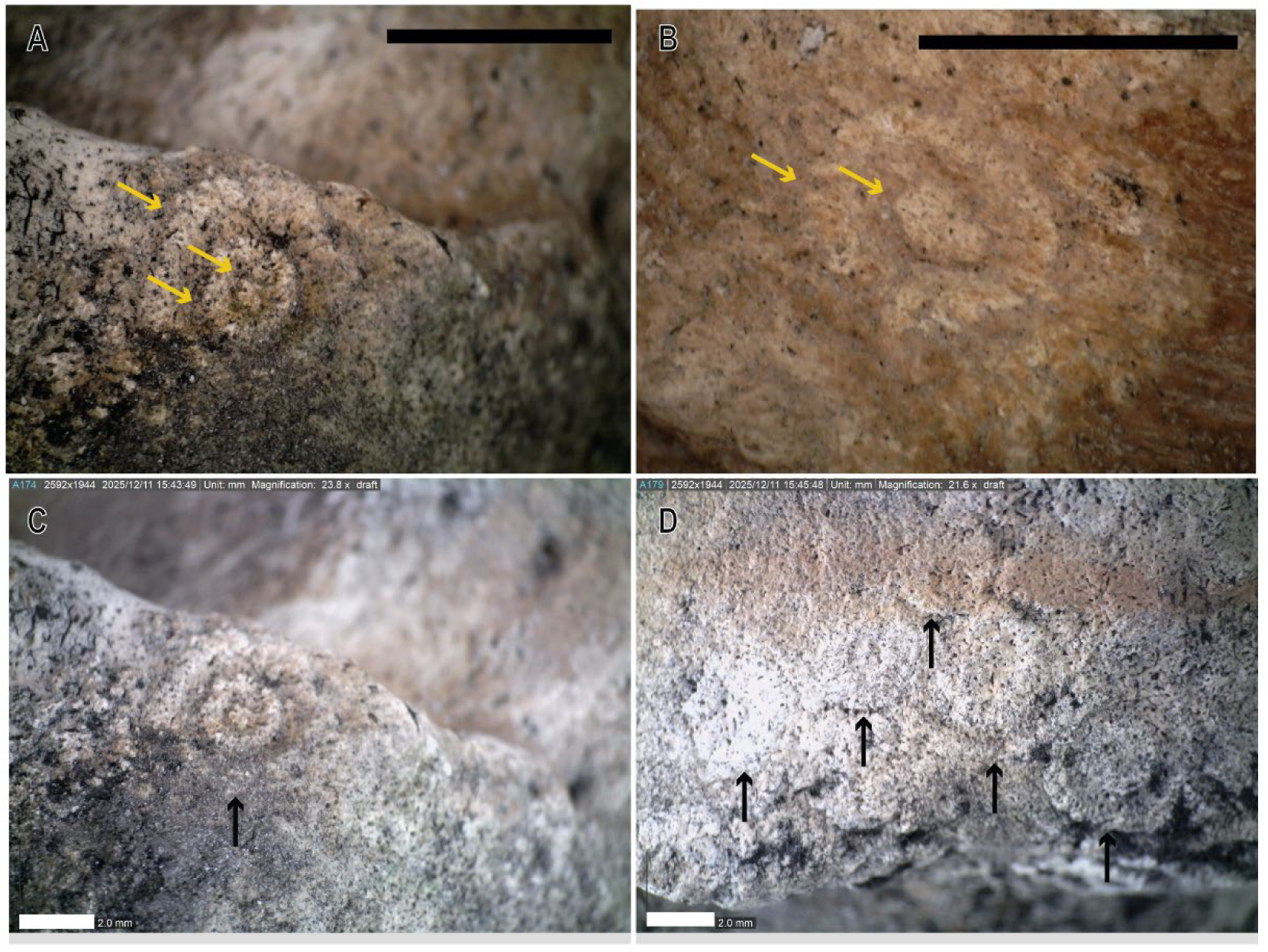
Circular target etching across bone surfaces. Yellow arrows indicate etched rings, and black arrows indicate the presence of features. Isolated features shown in A-C, and multiple, partially overlain features in D.

Pitting was common and not significantly different between wet and dry burial contexts. It was the most common corrosive feature at GWS. Extensive pitting in underwater contexts culminated in general surface loss where the isolated pitted features became continuous, destroying large regions of bone (Fig 7). Underwater, corrosive pitting and floral pitting were not congruent. General pitting without any signs of floral agent were present across a wider range of depths below water surface (-2.8m to -17.2m), whilst pitting associated with floral agents was restricted to the shallow underwater regions of GWW (-2.8 to -5.4m).

Surface corrosion was significantly greater on bones from underwater, and the only pre-collection bone surface modification to significantly increase in frequency through time (Table 5-6). Expression of corrosion varied across sites, likely a product of different agents. Superficial surface corrosion was most common in both the wet and dry corrosion sub-assemblages (GWW=83.1%, GH=77.8%, GWS=91.7%). Underwater, a continuous expression (70.3%) was associated with shallow depths at GWW whereas the deeper waters at GH were dominated by discontinuous corrosion features (66.7%; Fig 7). Specimens from dry contexts presented with both continuous (56.5%) and discontinuous corrosion (43.5%). Low levels of deep, continuous corrosion were also identified across all environmental conditions (GWW=10.9%, GH=22.2%, GWS=8.7%).

A spatial relationship was identified between surface corrosion features, types of staining, and biological agents. Some surface corrosion features were stained green, black, or blackish purple, with sharp margins. Black staining was significantly more common in the aquatic assemblage (*p*<0.001; GH=13.2%, GWW=23.6%, GWS=2.9%), whilst green staining was observed in both dry and wet settings at similar frequencies (*p*=0.090; GH=10.5%, GWW=23.0%, GWS=33.3%). Stains were not always associated with corrosion, particularly in dry, surface conditions where 34.7% of bones presented green stains but only 25.5% experience surface corrosion. In wet contexts, a chi squared test for independence showed significant associations between surface corrosion and biofilms (*p*<0.001), surface and black staining (*p*<0.001), and black staining and biofilms (*p*<0.001).

Microbial biofilms and localised algae were identified as the biological agents responsible for the staining and associated with corrosion in approximately 20 - 60% of specimens (Table 5). In some cases, biological agents were observed on the bone (Fig 9D-F), whereas only continuous black stains with sharp margins remained on bones from submerged contexts. The morphology of the stains was not consistent with manganese staining [86] but consistent with spectra produced by unstained bone surfaces on the same sample, confirmed by pXRF analysis (S2 Fig). Staining did not cover the entirety of the bone but was localised in distinct regions (Fig 9A-C). Collection photos show that staining is not exclusively associated with burial within sediment, or exposure to water, but rather is indicative of the area of bone exposed to light (Fig 9A-B).

**Fig 9.**
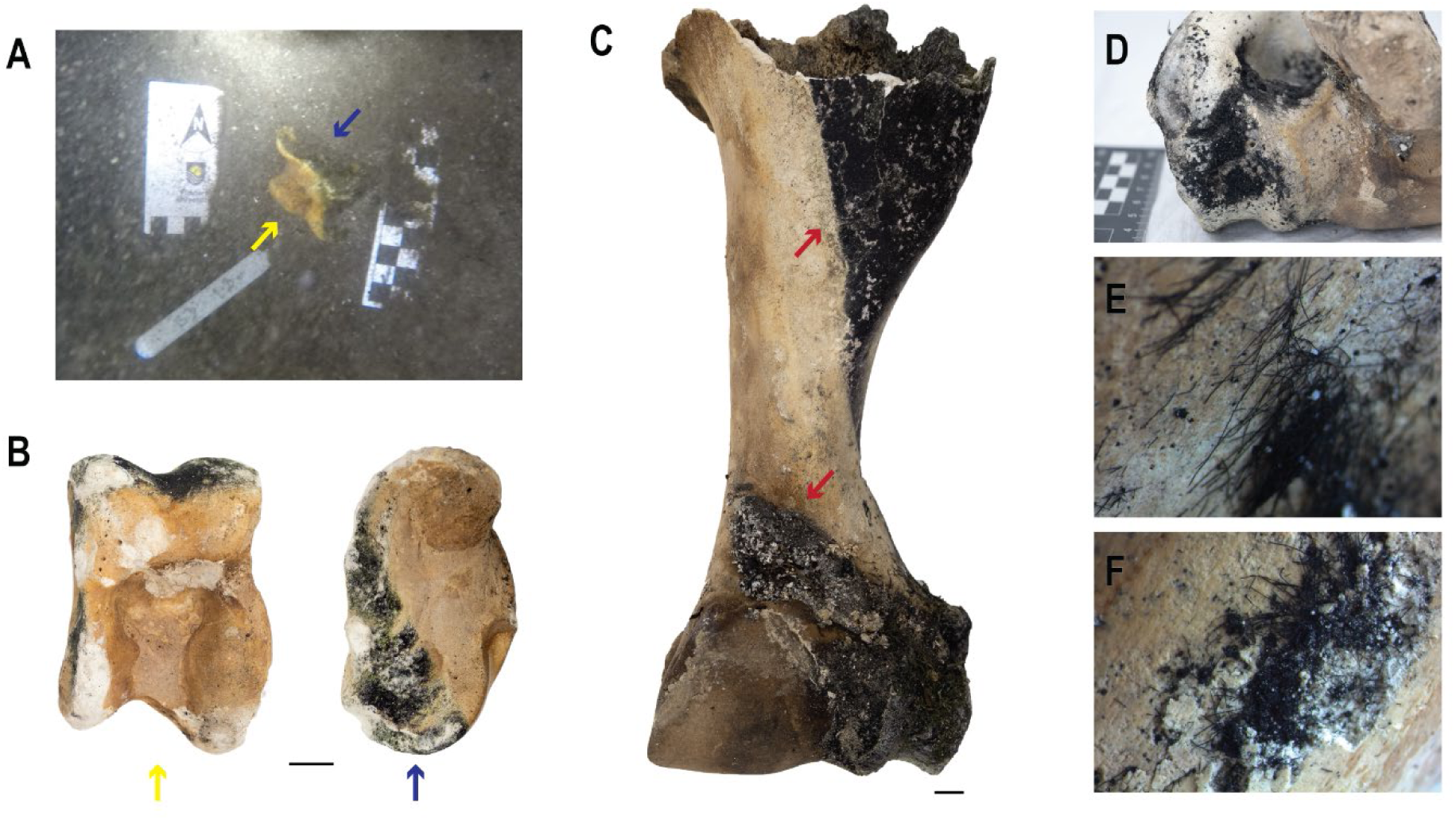
Biofilms, biota, and black staining associated with sunlight and surface corrosion. Scale bar is 1cm. A: In situ collection photo of astragalus (GWW09) with the exposed stained surface (blue arrow) facing upward and exposed to light compared to the unstained surface (yellow arrow) facing the dark zone; B. Astragalus from A, highlighting staining, and showing the unstained surface that was buried; C. Black biofilm staining and corrosion on humerus (GWW50) with sharp margins (red arrows); D-F: black biota across bone associated with corrosion but not staining.

**Fig 10.**
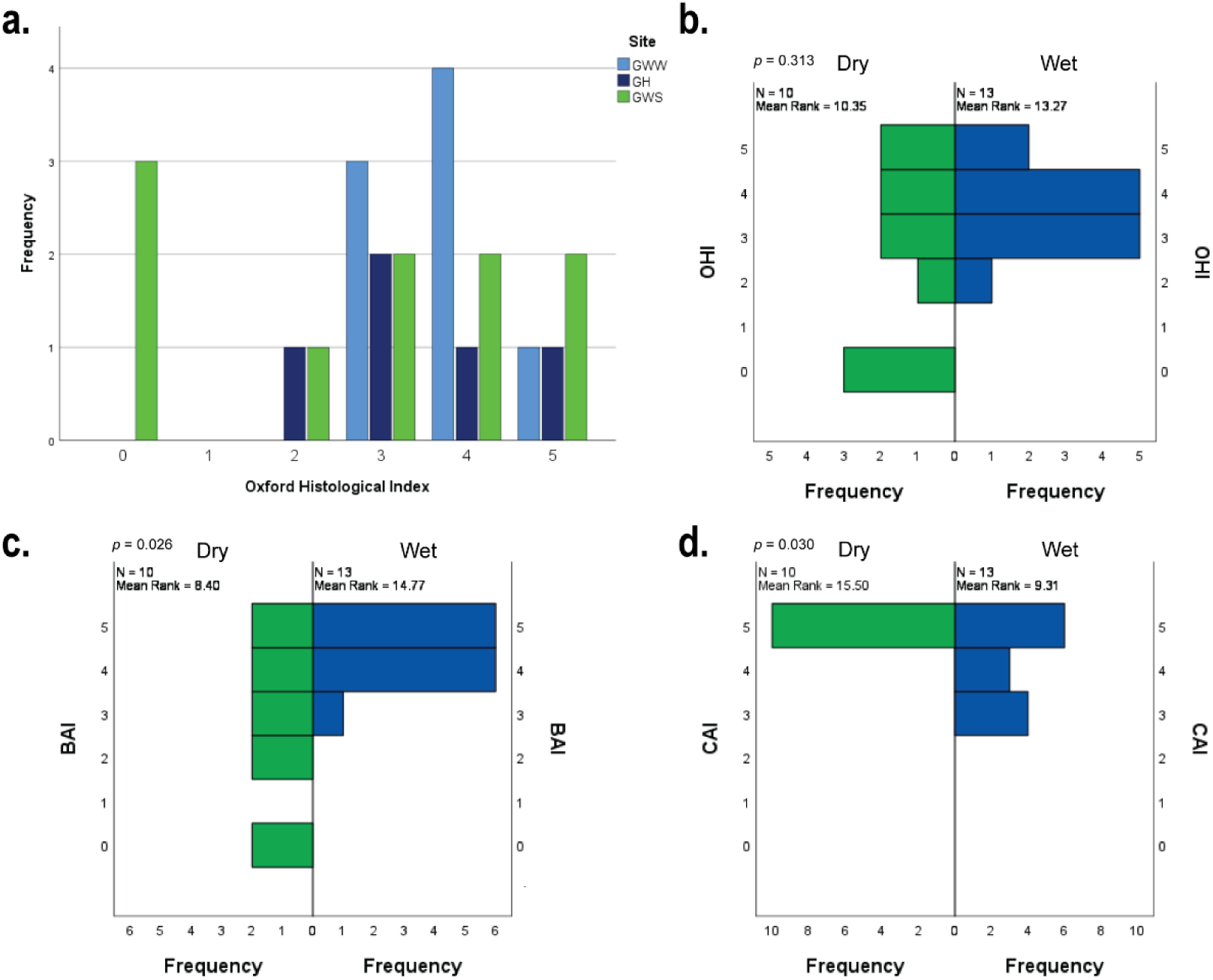
Distribution of diagenesis across each site, measured by the Oxford Histological Index (A)[87], and diagenetic indices recording: B: general; C: bacterial; and D: cyanobacterial degradation across wet and dry burial contexts.

In aquatic settings, the presence of biofilms and algae, surface corrosion, and staining varies with depth. Black stains were not recovered at the deepest sites of the caves, and green staining was limited to shallow regions (GWW: <-4.4 m; GH: <-13.3 m). The deepest example of a biological agent, black or green staining, and surface corrosion together was recovered from -13.3 m at GH. Small sample sizes may influence these results, as samples collected from depth consist of only single or few specimens.

### Histotaphonomy

Bone diagenesis ranged between completely modified (0) and well preserved (5) bone (Fig 9, Table 3 in S1 Appendix). Whilst specimens from dry contexts in GWS tended to exhibit poorer preservation compared to those from wet assemblages, Oxford Histological Index (OHI) distributions did not vary significantly (*p*=0.313). However, possible cyanobacteria tunnelling (CAI) was significantly more prominent in wet versus dry environments (*p*=0.030), whilst bacterial attack was significantly more common in dry bone microstructure (*p*=0.26). The degree of modification was not correlated with time since deposition in wet contexts (OHI: *p=* 0.295, BAI: *p=* 0.445, CAI: *p=* 0.234) or across the pooled whole assemblage (OHI: *p=* 0.089, BAI: *p=* 0.413, CAI: *p=* 0.922). Further, depth below water surface did not significantly impact OHI (*p*=0.913), BAI (*p*=0.288), or CAI (*p*=0.581), but small sample size limits statistical power (Table 4 in S1 Appendix).

#### Patterns of bioerosion and degradation

Bones from wet environments presented with well-preserved bone with isolated areas of bioerosion that were limited to the sub-periosteal, peripheral bone region (GH n=5, GWW n=8, Fig 11). Seven samples from wet burial contexts presented with a type of corrosion that formed a scalloped or bridged structure across the bone surface (Fig 12, A1-A2). These were not identified across any sample from dry contexts. Only a single specimen (ID: GWW_76), found at the deepest collection site at GWW (-5.4m) did not present with any evidence of degradation or bioerosion (Fig 12, C1-C2). Birefringence under polarised light across wet and dry sites was linked to general degradation reflected by the OHI. Regions of preserved bone were consistently birefringent in bones from wet and dry contexts. Bone marrow was still present in the medullary cavity of specimens (GWW, n=4) associated with burial across a decadal time scale.

**Fig 11.**
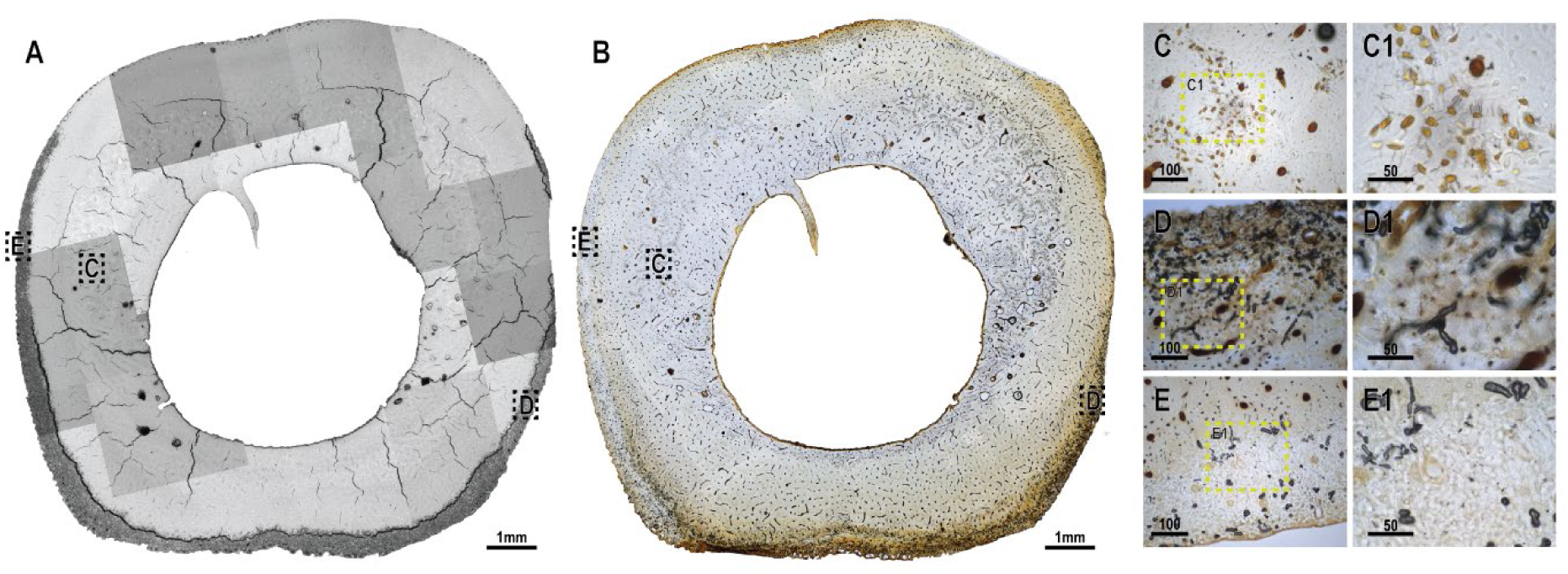
A and B are cross-sections from a sheep metatarsal (ID: GH_19) showing typical bioerosion patterns seen across specimens from Gouldens Sinkhole (ID: GH19) comparing: (A) backscattered scanning electron microscopy (bSEM) with (B) transmitted light histology. Close up images under transmitted light of: C-C1: enlarged canaliculi; D-D1: cyanobacteria tunnelling; E-E1: black and white tunnelling in region of cyanobacterial tunnelling under bSEM (A).

**Fig 12.**
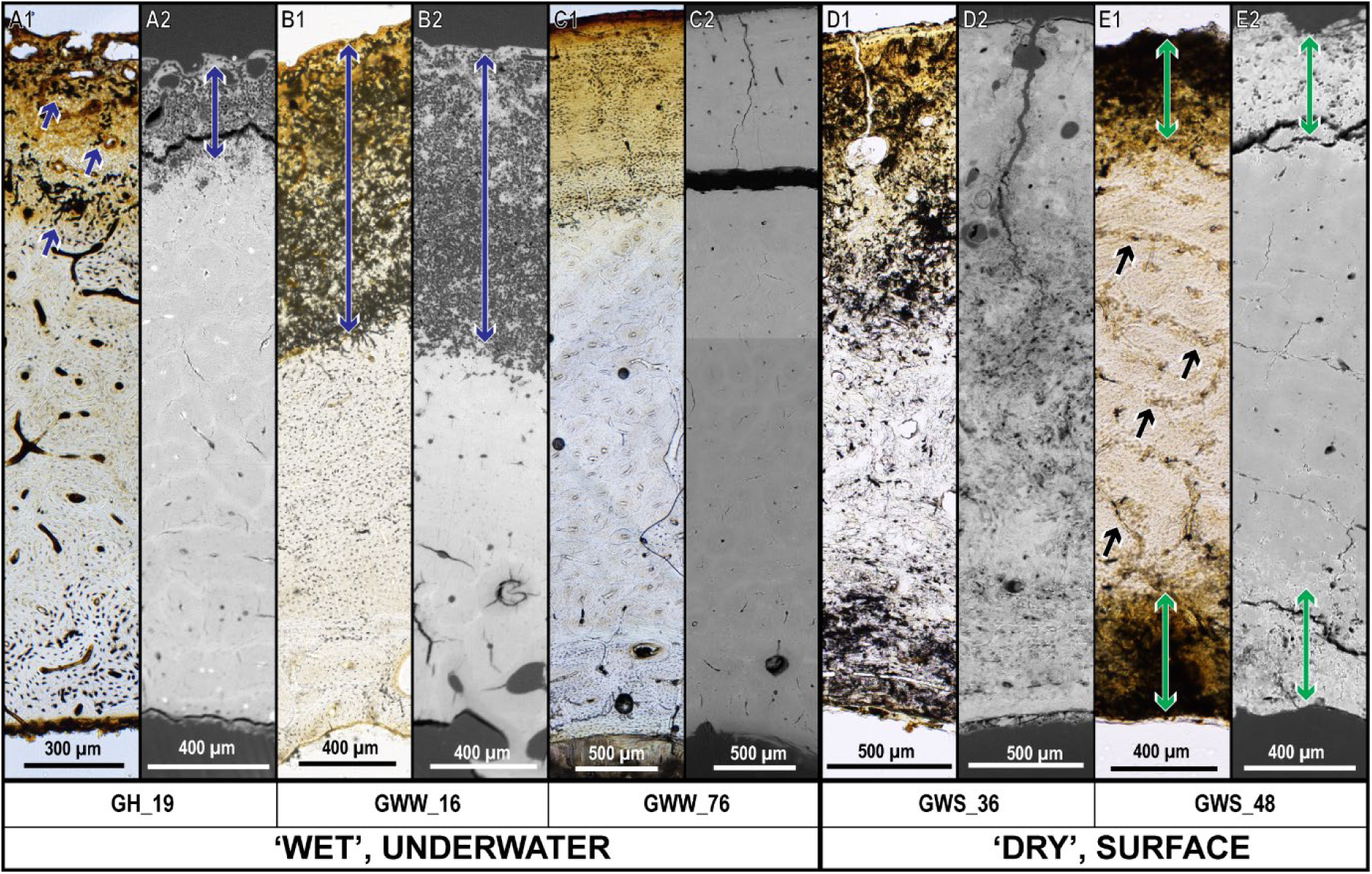
Example of bone histology types across samples from ‘wet’ underwater and ‘dry’ surface burial contexts presenting the same region of interest strip under transmitted light microscopy (A1-E1) and backscatter scanning electron microscopy (A2-B2). In A and B, blue coloured arrows indicate examples of peripheral cyanobacterial tunnelling. In C, well preserved bone with no bioerosion is presented next to complete degradation in D. In E, green coloured arrows indicate non-cyanobacterial degradation of sub-periosteal and sub-endosteal envelopes, and black coloured arrows indicate mid-cortical budded MFD associated with vascular canals only observed through transmitted light microscopy. (note: variation between images is a result of the different histology blocks used for each technique). Images were taken from ovicaprid metatarsi (ID: GH_19, GWS_36, GWS_48), a sheep radius (ID: GWW_76) and a cow rib (GWW_16).

Microscopic features in bone samples from wet sites were identified as tunnelling (possible by cyanobacteria) where resorptive regions approximately 5 - 15 µm in width were not surrounded by a hyper-mineralised border and did not contain bacterial foci (Fig 13). Where present, limited areas of the total bone surfaces showed tunnelling at GWW (n=7, min=0.4%, max=25.0%, *x̅*=10.4%, σ=10.7%) and GH (n=5, min=0.3%, max=19.8%, *x̅*=11.5%, σ=6.3%). Conversely, where other MFD types were present, they impacted cortices to a greater extent (GWW: n=7, min=1.5%, max=100.0%, *x̅*=21.9%, σ=32.1%; GH: n=3, min=1.9%, max=43.3%, *x̅*=16.8%, σ=18.7%). These wet features were also restricted to the peripheral surface in all but one specimen where they extended across the endosteal envelope. In one instance, possibly cyanobacterial tunnelling was layered beneath other MFD, possibly superimposing and reworking old modifications (Fig 14). The chalky bone texture identified to be more frequent in wet assemblages (Table 5) occurs at the same location as possibly cyanobacterial tunnelling (S3 Fig).

**Fig 13.**
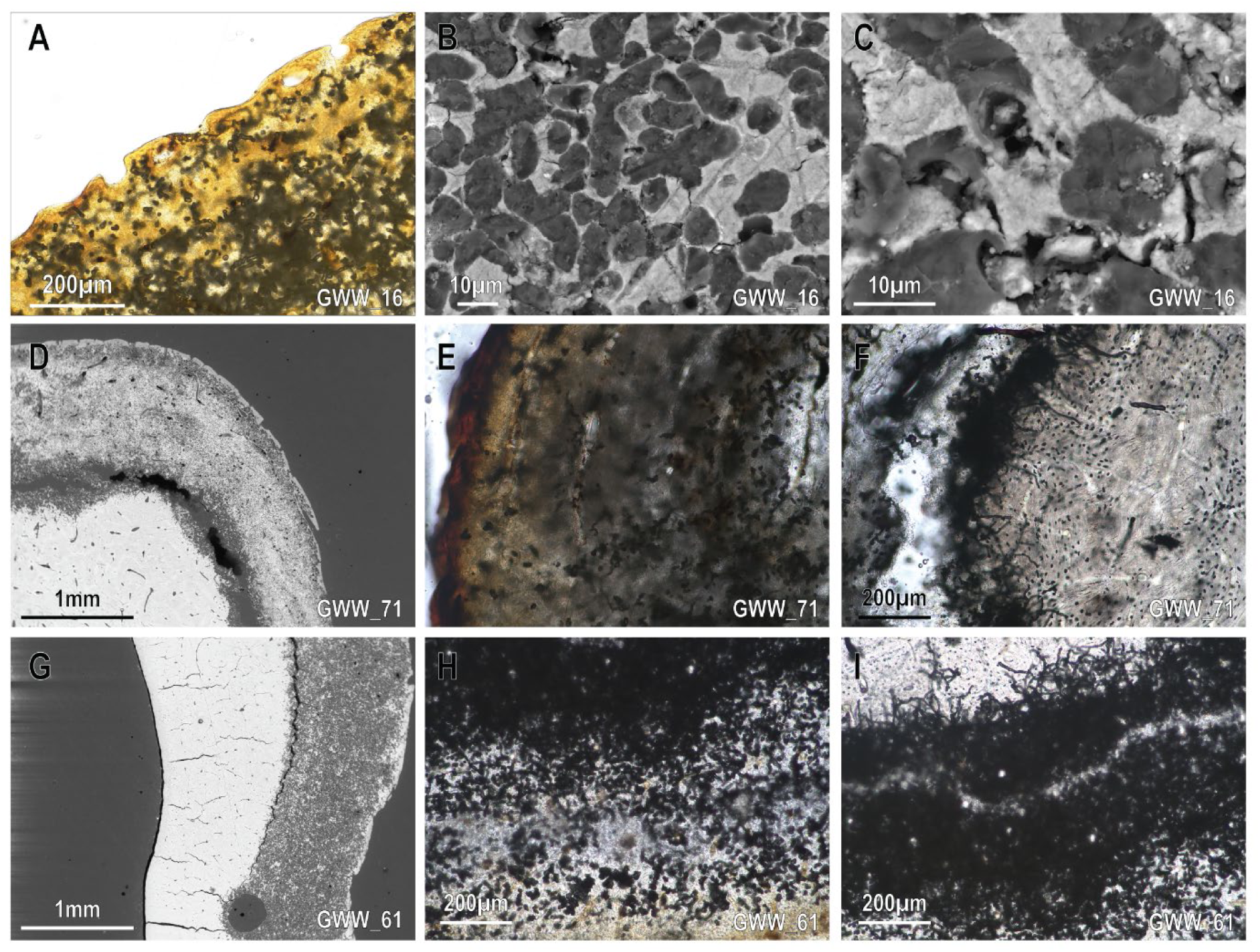
Close up of wet site features at Green Waterhole under backscatter scanning electron microscopy (bSEM) (B,C,D,G) and transmitted light microscopy (A,E,F,H,I). A: peripheral scalloping and subperiosteal bioerosion; B-D: possibly cyanobacterial tunnelling without hypermineralised boundaries; D,G: subperiosteal bioerosion restricted to the exterior bone region; E: close up of bioerosion with unidentifiable features; F: separation of bone at the margin of bioerosion with Wedl tunnelling extending into well preserved bone; H-I: close up of bioerosion features associated with G, with a black mass of Wedl tunnelling located within but not adjacent to the periosteal surface (bottom of images). Images taken from a cow rib (ID: GWW_16; A-C), cow ulna (ID: GWW_71; D-F) and an ovicaprid metacarpal (ID: GWW_61; G-I).

**Fig 14.**
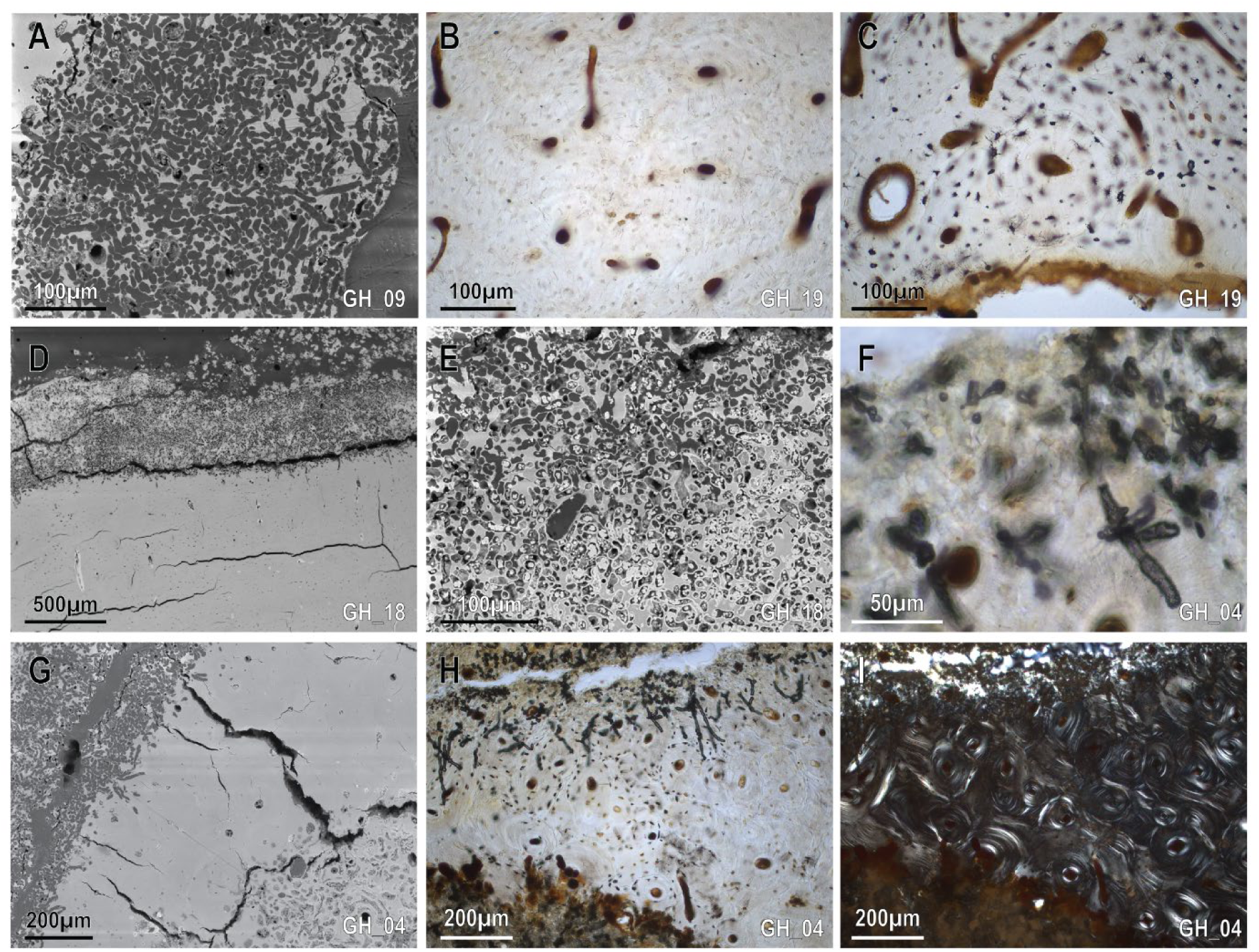
Close up of wet site features at Gouldens Sinkhole under backscattered scanning electron microscope (A,D,E,G) and transmitted optical light, (B,C,F,H), and polarised optical light (I). A: dense region of Wedl tunnelling across bone surface; B-C: well preserved bone with visible primary canals and osteocyte lacunae; D: biotic attack at periosteal boundary with mixed terrestrial and possibly cyanobacterial tunnelling (E); F: close up of Wedl-tunnelling; G-I: tunnelled exterior bone with Wedl-tunnelling extending into well preserved and birefringent matrix, followed by extensive terrestrial biotic attack towards endosteal surface. Images were taken from Ovicaprid metatarsi (ID: GH_09, GH_19; A-C), a kangaroo tibia (ID: GH_18; D-E), kangaroo rib (GH_04; F-I).

Whilst degradation extending from the subperiosteal envelope was identified equally across both wet and dry samples (*p*=0.75), the presence of degradation across only the peripheral region was found to be statistically significantly associated with wet environments (*p=*0.018). In wet environments, the depth of possible cyanobacterial tunnelling from the peripheral surfaces varied within a sample, often only presenting in isolated areas (Fig 11, S3 Fig). Maximum depths of this tunnelling ranged between 109 µm and 2254 µm (Fig 12, B1-B2), with greater average penetration in the shallower regions at GWW (*x̅* = 1114.4 µm, σ = 573.7 µm) compared to the deeper areas of GH (*x̅* = 552.5 µm, σ =175.9 µm). Collection depth was not significantly correlated with the presence of bioerosion (*p*=0.116) or maximum depth of bioerosion (*p*=0.270). In dry settings, specimens presented with an additional sub-endosteal modification, where both regions were dominated by MFD not associated with cyanobacteria (Fig 12, E1-E2). Bones in wet environments that presented with both sub-periosteal and sub-endosteal peripheral degradation also presented with a fractured shaft, thus exposing the medullary cavity to the aquatic burial environment.

Type and location of degradation were highly variable across specimens collected from dry contexts (Figs 15 and 16). Three groups were identified: 1) near complete preservation (>95% preserved; n=2); 2) bioerosion of the sub-endosteal and/or sub-periosteal regions (n=5); and 3) near complete degradation (>99% degraded; n=3). Tunnelling possibly associated with cyanobacteria was identified across 33.33% of the dry surface samples whereas other MFD were present across 90% of specimens. Of the specimens where this tunnelling was present, it was minimally invasive (min.=0.1%, max.=3.0%; *x̅* =1.2%, σ =1.3%) compared to extensive degradation generated by other agents (min.=4.0%, max.=100.0%; *x̅*=47.1%, σ =38.0%). Under transmitted light microscopy, localised budded MFD degradation was identified in dry samples associated with vascular pathways (Fig 12, E1). These mid-cortical MFD features were not picked up by bSEM imaging (Fig 12, E2). Few samples (n=3) from dry burial contexts also featured large, irregular circular empty spaces across the sub-periosteal pocket (Fig 12, D1-D2). These were not associated with natural bone resorption at primary or secondary osteons. No bones from wet environments presented with these features.

**Fig 15.**
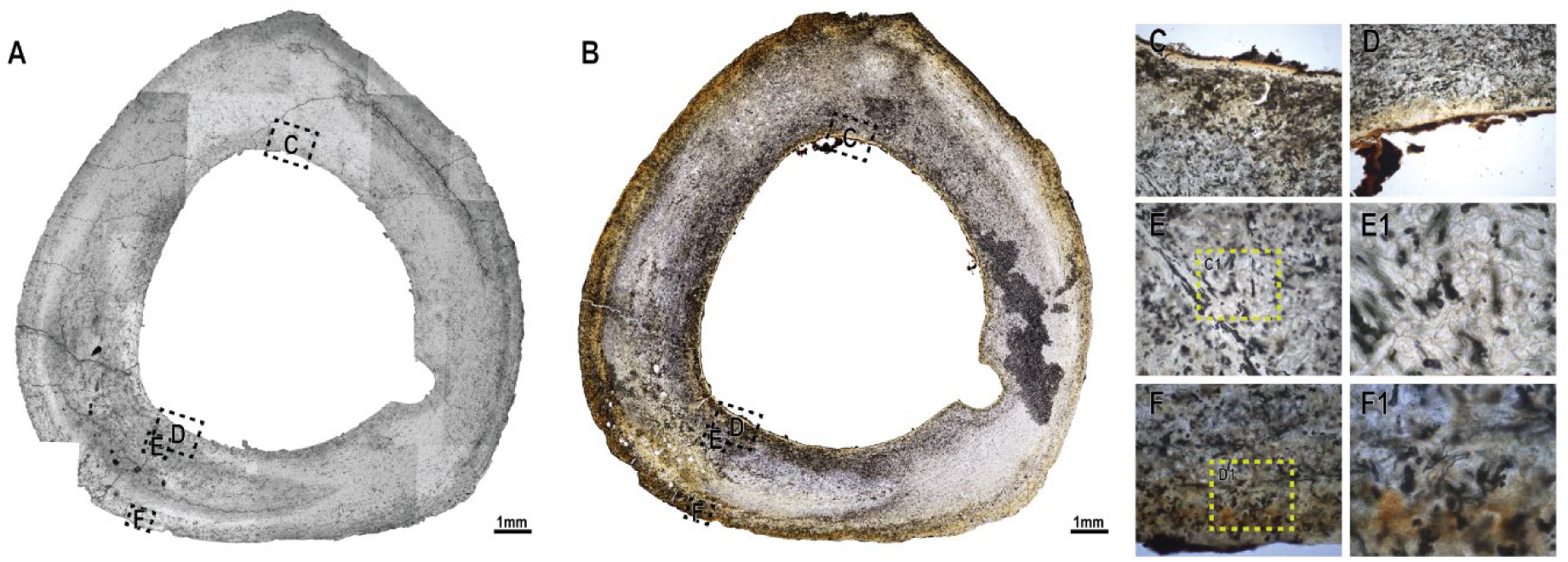
Ovicaprid tibia cross section (ID: GWS_86) from Green Waterhole Surface with extreme degradation. Comparing backscattered scanning electron microscope (A) with transmitted light histology (B). Darkened region on right of B due to sample preparation and not pre-collection staining. Close up images under transmitted light of: C-D: endosteal surfaces; E-E1: budded MFD, F-F1: possibly cyanobacterial tunnelling.

**Fig 16.**
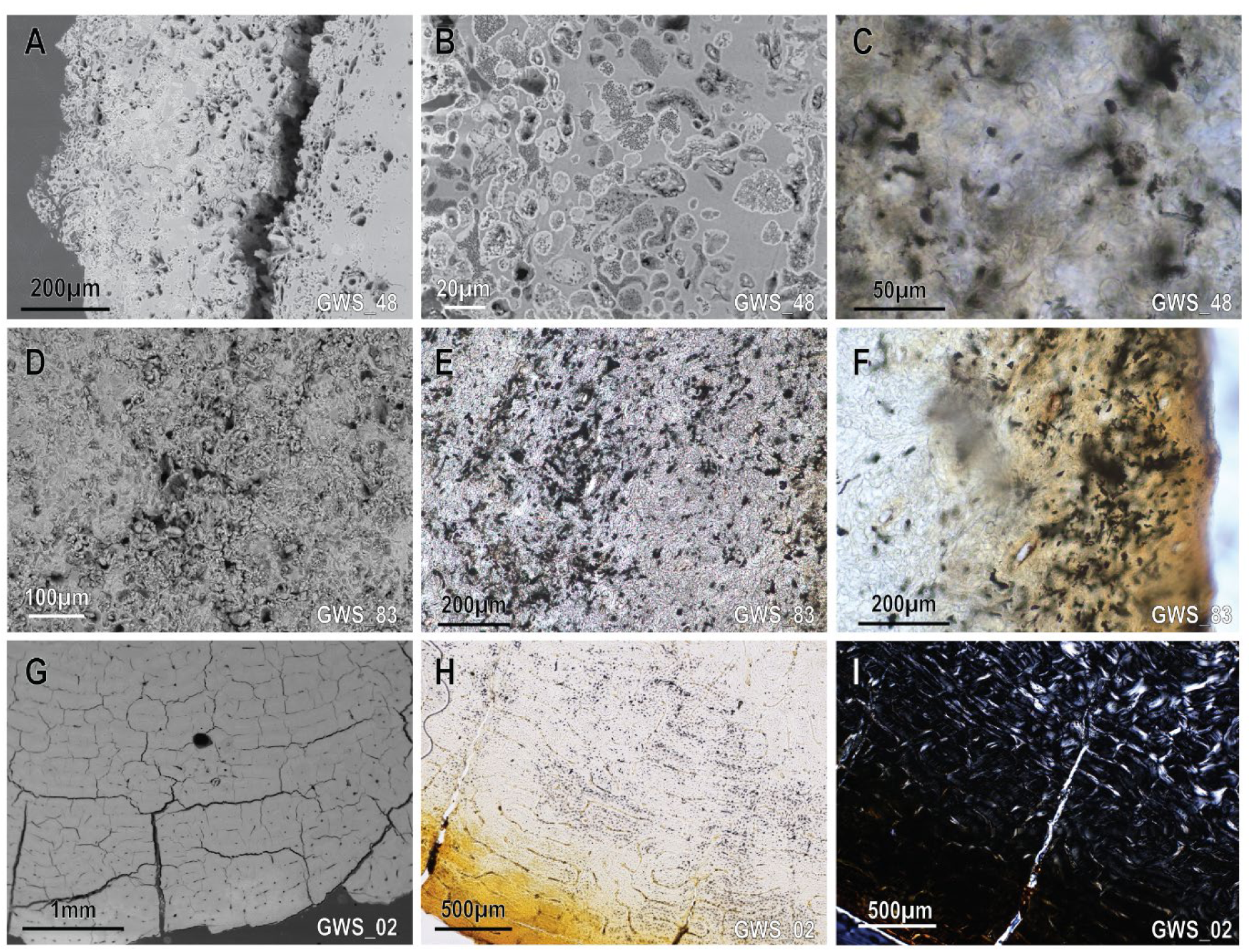
Close up of dry site features at Green Waterhole Surface under backscattered scanning electron microscope (bSEM) (A,B,D,G) transmitted optical light, (C,E, F,H), and polarised optical light (I). A: heavily degraded periosteal surface; B: closeup of bacterial attack with internal foci and hypermineralised boundaries; C: bacterial attack under transmitted light; D-E: increased porosity and degradation across central bone region; F: stained periosteal surface with biotic attack; G: well preserved bone with flaked periosteal surface, internal cracking a product of SEM chamber; H-I: same bone as G, with stained and not birefringent periosteal surface followed by well preserved and birefringent bone. Images were taken from ovicaprid metatarsi (ID: GWS_48, GWS_83; A-F) and a sheep tibia (ID: GWS_02, G-I).

Across wet and dry contexts, microfractures were not associated with histological structures, instead limited to general destruction through the cortex. Radial cracks originating from the periosteal surface and penetrating various depths into the cortex were observed across 70% of the dry assemblage. These did not occur in any sample from wet burial sites. Underwater cave samples presented with fractures that ran parallel to the bones surface (Figs 13 and 14), most frequently at the margin between cyanobacteria tunnelling and unaltered bone matrix. These parallel fractures resulted in delamination of the surface (n=9). Although samples from dry burial environments also experienced surface delamination and weathering, surface fractures were angular and irregular (Fig 15). As removed bone surfaces cannot be measured due to their absence, the true maximum depth of bioerosion, and degradation indices of the original bone cross section cannot be identified.

Radial microfractures across secondary osteonal cement lines were not identified, regardless of burial condition or time since deposition. Although various forms of cracking were observed in bSEM images, these were created by the pressurised chamber where thin sample blocks (<3mm thick) were more susceptible to damage. Samples from wet sites appear more susceptible to SEM cracking compared to those from dry sites.

Raw histology and histotaphonomy measurements, and descriptions for each sample are provided in S2 Dataset.

## Discussion

### Site formation processes

Radiometric dating indicates that domesticate bones accumulated across a decadal (>54, <75 years) and centennial (>81, <184 years) scale. The youngest bones were deposited at Green Waterhole in the 1970s, highlighting that modifications recorded in this study are at least 50 years old. The impact of old, dissolved inorganic carbon in cave waters on terrestrial animal bones deposited in water is not well understood. The freshwater reservoir effect can artificially increase the radiocarbon ages of living animals, plants and sediments from aquatic systems (e.g. fluvial and lacustrine), as well as their consumers, but not of bones or material deposited into freshwater sites [88, 89]. However, dissolved inorganic carbonates in groundwater can contaminate bioapatite during diagenesis through ionic exchange with older carbon from sedimentary rock, effectively aging bones [90]. A Late Pleistocene bone sample collected from an underwater cave in Mexico was identified as possibly contaminated and older that the ^14^C enamel yields and uranium/thorium dating results from the same site [91]. Discrepancies between enamel and bone carbonate ^14^C ages are found across some, but not all, terrestrial sites, and thus differences may be a product of site-specific diagenesis pathways [92]. In this study, enamel was not measured, and further work is required to understand the impact of dissolved inorganic carbon on bone assemblages in underwater caves.

Bone specimens across Green Waterhole reflect culturally and naturally accumulated deposits. The dry and wet assemblages at Green Waterhole are dominated by domesticate species and unarticulated bones, with low numbers of native remains. Butchery marks and the abundance of butchery portions at GW suggest that the remains were anthropogenically dumped at the site after processing, supported by the presence of historic refuse scattered around the margins of the dry doline [93, 94]. This anthropogenic assemblage is comingled with what is likely a natural deposit of native remains that did not exhibit butchery marks, however we can’t rule out that the presence of native animals in this site were also a result of anthropogenic accumulation that did not leave any visible marks.

Almost equal proportions of native and domestic taxa with no evidence of butchery were recovered from Gouldens Sinkhole. This suggests that these bones were naturally deposited at the site, however historic refuse across the talus cone highlights inputs from dumping at some point. Low levels of weathering suggest individuals drowned in the sinkhole, or bones were quickly reworked into the site. Evidence of predation (gnawing) across a few specimens indicates that at least some of the assemblage was re-worked from a terrestrial environment [7].

Signs of primary deposition at Gouldens Sinkhole are further skewed by likely intra-site reworking down the talus cone. Bone specimens with a similar length and breadth were found at deeper deposits, mirroring observations of sorting across fluvial [63, 95] and steep sloped hills [96]. After deposition, bones in underwater caves were trapped in the suspended, fine-grained allochthonous sediment of the talus cone. Whilst further analysis of sphericity and bone weight may provide further insights [95–97], basic measurements of shape highlight patterns of transport in a highly sloped, low energy flow cave setting. This pattern was not observed as strongly at Green Waterhole West due to the presence of a lower gradient slope, indicating that slope angle is strongly correlated with degree of sorting.

### Burial conditions in underwater settings

#### Bone surface modifications

Submerged cave assemblages were generally well preserved, with minimal element breakage. Lower frequencies of well-preserved bones at GWW compared to GH is a product of butchery practices, not an indicator of in situ breakage. Increased fragmentation and fractures across the dry assemblage is likely a result of trampling, a feature observed at other dry cave sites [98, 99]. Well preserved bones are a feature of underwater cave assemblages [3, 5], whereas dry caves and terrestrial aquatic sites are open to predation, trampling and physical transport [18, 19].

Submerged bones were soft, and prone to collection and drying damage. Biomineral and organic analyses of bone matrix were not conducted here, however submersion does impact bone collagen and crystallinity [22, 43]. Modifications to bone mineral and organic structure that results in soft bones can remove prior evidence of abrasion and physical modifications. For example, abrasion modifications formed by cave diving line prior to collection were similar in colour and profile to collection damage.

The weathering scale did not distinguish wet and dry cave settings, or differences between decadal and centennial deposits, highlighting the need for a nuanced approach to understand environmental weathering [18]. Increased flaking and exfoliation across dry sites may be valuable in identifying dry deposited specimens, where wet deposited bones experience chemical desquamation exfoliation instead of physical flaking [21, 100]. Bones from waterlogged soils showed exfoliation post-collection after drying [1], and identifying the presence of an original bone surface may be helpful in identifying bones that have experienced changing hydrological conditions.

Waterlogged bones from phreatic caves do not dry evenly and this produces surface delamination. Differential expansion and contraction pressures eventually led to the separation of external layers of bone, ‘popping’, and finally bone removal [101]. Delamination is common in salt-water conditions, as expanding salt crystals enhance the effect of delamination [102]. Bones from waterlogged soils also present with similar effects [1], and experimental studies highlight that increased exposure to wet-dry cycles will result in cracking that mirror early stages of the weathering scale [103]. Delamination in our samples occurred during drying and thus may also reflect the impact of shifting water tables in older bones from phreatic caves that occur either seasonally or with long term sea-level changes.

The localised corrosion of thin bones resulting in complete loss of material observed in Quaternary subfossil bones from underwater caves [3, 104] was not identified in this study. Those specimens were collected in salt-water, below a halocline, and thus this type of corrosion may be linked to salinity [102], which is not applicable here.

Distinct expressions of corrosive biotic attack were identified in this study, with increased etching in dry settings and increased surface corrosion in wet environments. Pitting was consistent across environments. Corrosion associated with floral attack are common, as organisms excrete chemical compounds, degrading bone for nutrient uptake [105]. Therefore, linking corrosive features to specific flora is difficult due to the similar expressions of corrosions across plant taxa. In this study, however, different frequencies of etching and surface corrosion likely reflect the difference between plants with roots (vascular plants) that are linked to dry landscapes, and rootless non-vascular plants (e.g. bryophytes and thallophytes) and biofilms that are linked to wet settings [106, 107].

In the submerged caves, surface corrosion is the only bone surface modification that increased significantly with time. Previously, shallow, continuous surface corrosion features were attributed to biofilm corrosion across Miocene lake fossils [81, 108]. In these fossilised samples, bone surfaces were not completely degraded, indicating that for the specimens that survived into fossilisation, corrosion ceased at some point prior to complete destruction. Here, surface corrosion was frequently observed with pitting around the edges of bones, suggesting that successive pitting events may result in complete loss of bone. Further investigations into the speed of floral attack, and why it terminates, is required to understand how surface corrosion proceeds through time.

Although etching was mostly associated with a dry environment, a specific form of etching was observed underwater: a circular target etching that occurred in clusters or singly. No agents could be associated with either form. Predation in marine settings by barnacles and molluscs create homing scars [28], and similar circular patterns attributed to *Thatchtelithichnus* have been viewed on subfossil mammal bones from seafloors [109]. Ecological analyses of local biota across the study sites are needed to identify the possible agents responsible for etching expressions. Whilst the distinction between etching forms is beneficial in differentiating depositional environments, they were recorded infrequently.

There is a strong association between corrosion, biological agents, and black staining across bones from underwater environments. A clear taphonomic history outlining the timing and relationship between agent, corrosion, and stain cannot be deduced through this neotaphonomic framework. Stains were not caused by manganese precipitation or manganese-oxidising bacteria like those found in other waterlogged archaeological sites [86]. Instead, they are likely a result of cyanobacteria, fungal attacks, or biofilms that secrete acids capable of eroding bone, stone, and metals [110, 111], stain bone surfaces black [61, 112, 113], and produce pigments in caves where exposed to light [8]. In our samples, continuous pigmented black stains with sharp margins are not associated with burial in sediment or exposure to water but are linked with light availability.

Light in caves is restricted to entrance zones, typically unidirectional, and easily obstructed by objects that cast shadows. Associations between biofilms, flora and light have been identified across cave systems before [8], however microbial communities can also be found deep within dark underwater cave systems [114]. The deepest example of surface corrosion associated with green or black staining and a biofilm or algal agent was at Goulden Hole, 13.3 metres below water surface. This collection context is within the sinkhole opening area, exposed to light, although seasonal algal growth at the surface can restrict light significantly even within this area. Whilst surface corrosion and staining were independently found in deeper regions of the cave, they did not present together. Light variability thus accounts for the seemingly random distribution of this feature across bone surfaces, and different patterns observed with depth.

Bone surface modifications in underwater caves vary relative to dry environments, and those from other aquatic landscapes (Table 7). Walker and Louys [7] identified features that may be attributed to distinct landscapes, and here we confirm the lack of aquatic bleaching outside of saltwater environments, marine aquatic fauna modifications [102], and homogenous or heterogenous rounding under different hydraulic regimes [24, 115, 116]. Diatom presence, pits, or orientation pattern were not assessed in this study [21].

**Table 7.**
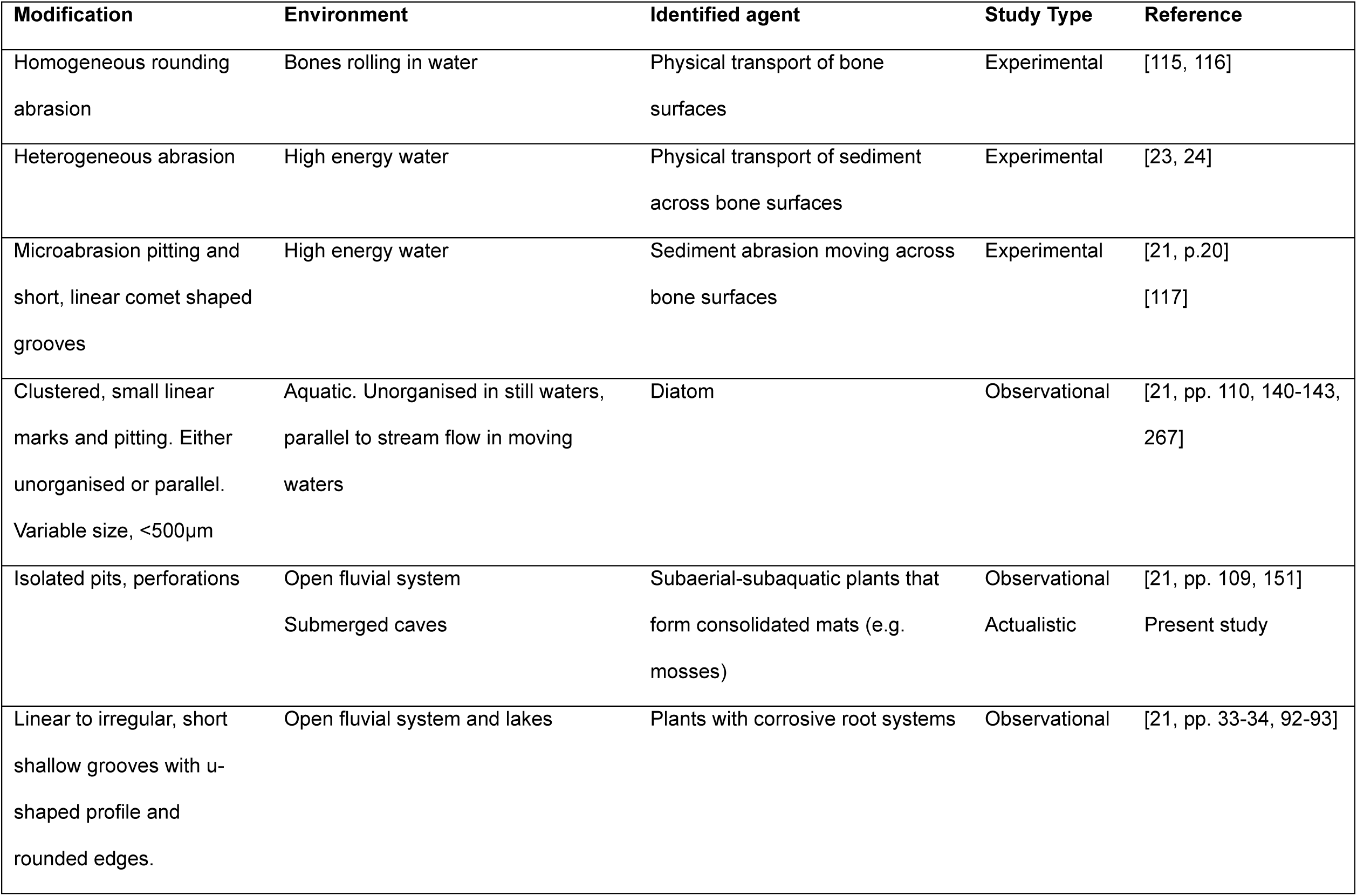
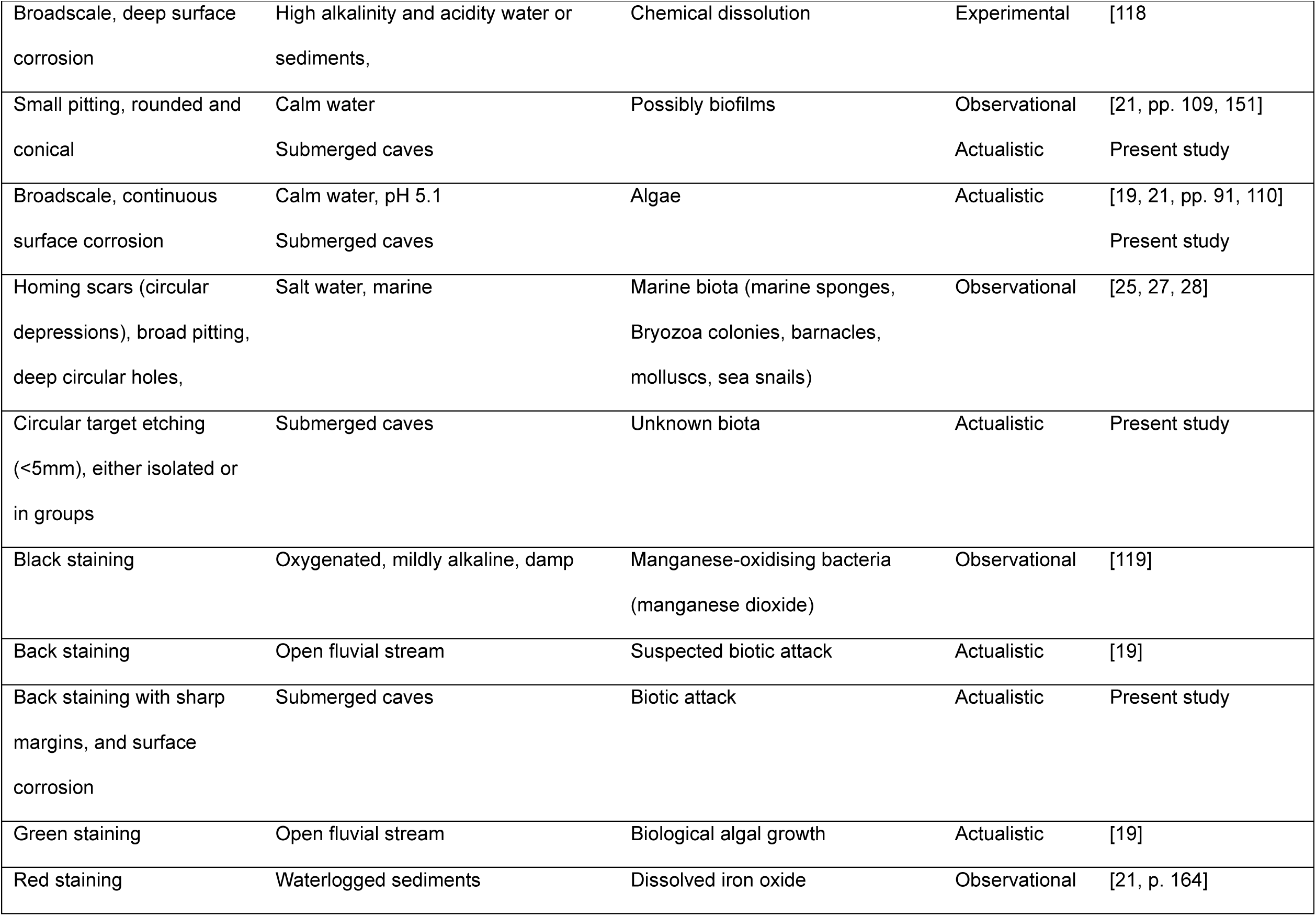
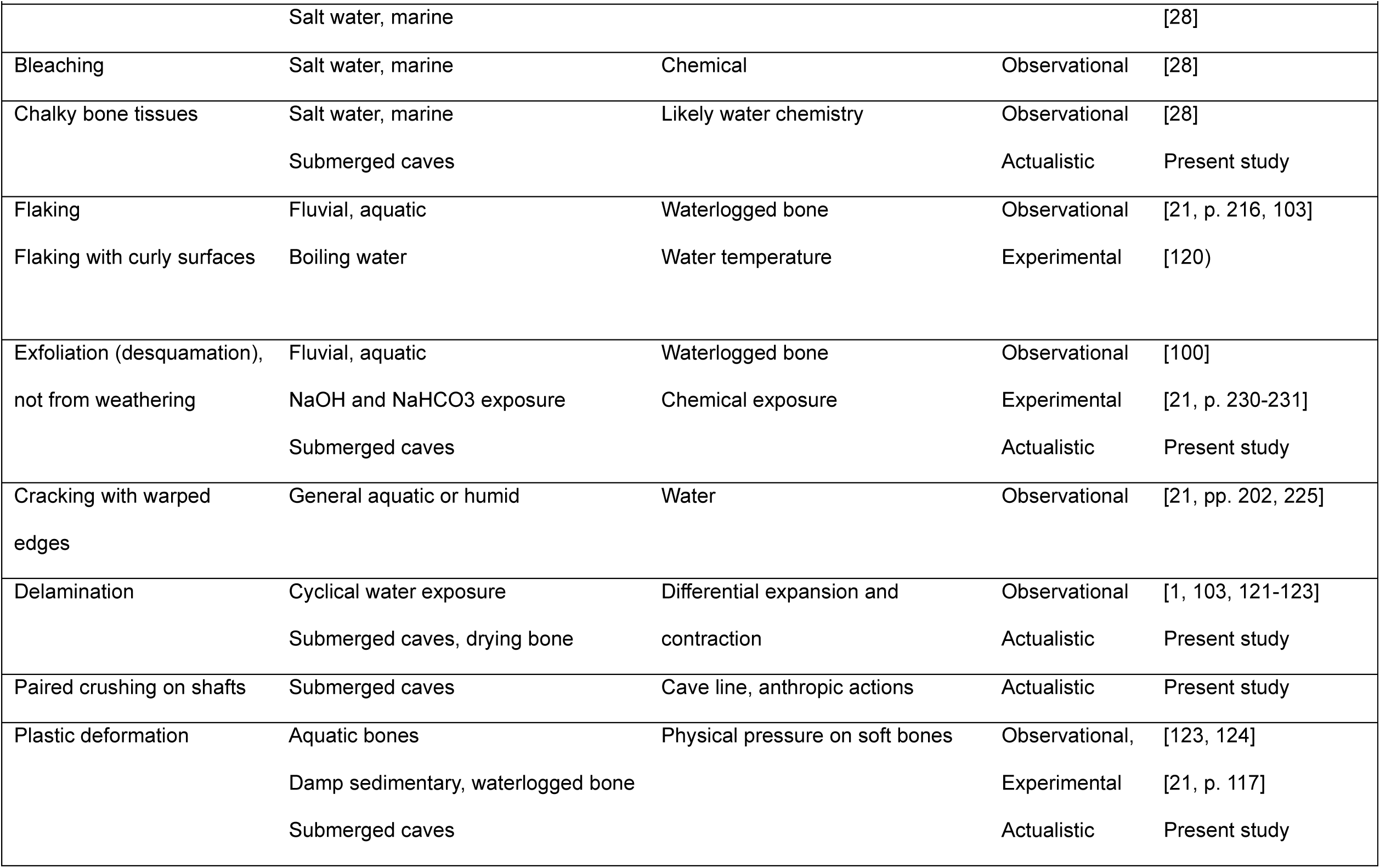
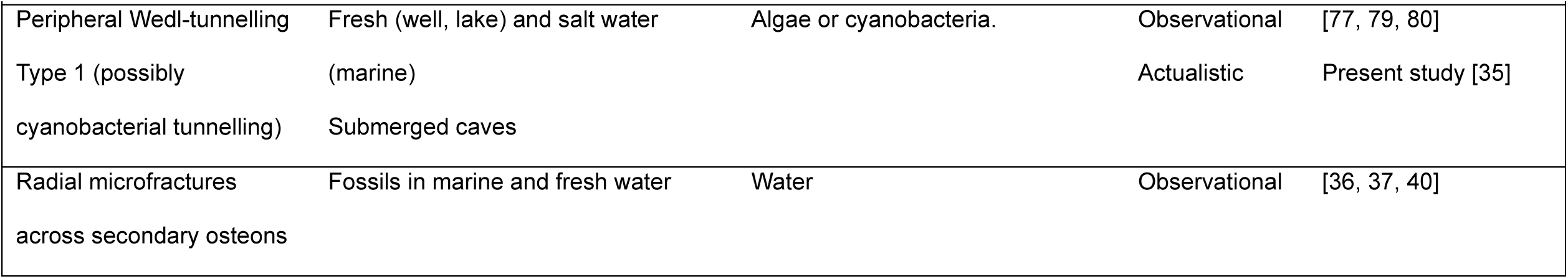
Aquatic bone surface modifications identified through different research frameworks. Modifications listed are restricted to bone surfaces and microstructures. Only modifications identified as significant or unique to the Submerged cave environment are included.

The effect of time on bone modifications was minimal at our scale of observation. Whilst small sample sizes will inevitably bias results, data here suggest that patterns of wet or dry diagenesis will occur within fifty years. Understanding how and why bone matrices are altered can support our understanding of the early and late diagenetic process in underwater cave environments [13], and if taphonomic analyses can be used to distinguish between wet and dry deposition.

#### Microstructural diagenesis

Histotaphonomy presents clear, unique patterns of degradation linked with bones from either wet or dry conditions. Increased tunnelling that may be associated with cyanobacterial extending from exposed peripheral surfaces is significantly linked to the submerged cave environment, whereas other microscopical focal destruction features are more frequently observed in dry cave settings. Wedl-tunnelling Type 1 observed in this study has been previously identified in an actualistic study linking their presence to cyanobacteria euendolithis that bore into bone from peripheral surfaces [35, 80]. Whilst linking features of bioerosion to identify cultural burial practices or post-mortem intervals is debated [12, 76, 125], Wedl-Type 1 tunnelling has been identified in nearshore marine, deep marine, and lacustrine landscapes [35, 79, 81], and is established through a strong relationship between submersion, environmental DNA, and degradation [35].

Wedl-tunnelling Type 1 also occurred in some specimens recovered from surface deposits, but these were located at the lake shore boundary and thus experienced wetting during periods of higher ground water levels. Dry specimens away from the lake shore do not present with this type of tunnelling, mirroring other experimental and actualistic studies [35, 126]. Presence of tunnelling is thus also a feature of epiphreatic zone of caves where ground water fluctuates from wet and dry depending on seasonal rain inputs.

Morphology of Wedl-tunnelling Type 1 remains consistent across marine environments but is suggested to vary in terrestrial bodies of water. The presence of a remineralised, electron dense, tunnel border was suggested to differentiate marine from continental burial environments [82]. Unlike the patterns of tunnelling with hypermineralised boundaries identified on seven-million-year-old fossils from a palaeolake [81], the patterns in our underwater cave samples affected by ground waters were not surrounded by a hypermineralised border, aligning with other marine studies [35]. Different expressions may reflect local conditions or the mineralisation process during fossilisation.

One specimen (GWW76) from the underwater settings did not present with any tunnelling and was recovered from the darkest portion of the cave. No signs of reworking of animal predation were identified on this sample. In Bell and Elkerton [79], tunnelling on bone specimens from the Mary Rose warship was restricted to those found to be reworked and outside the warship exposed to light, compared to those recovered from within the confined, dark space of the ship that did not feature tunnelling features [79]. Areas of caves in complete darkness may thus not present any form of bioerosion. Whilst no statistically significant correlation was found between depth of collection and either presence or maximum penetration of tunnelling, this is likely a product of small sample size across different collection depths.

Estimating the timing of Wedl-MFD degradation is not well resolved. Our data indicate that under certain conditions, Wedl-tunnelling will not be present even 50 years after deposition. In some fresh water and marine settings, tunnelling did not present after 12 and 24 months [12, 43], and in other marine environments they occurred after 24 months [35] and four to five years [126]. Fossils exposed to tap water of varying pH under laboratory conditions did not exhibit microbial tunnelling after three weeks [127], suggesting that either the environment or timespan was not conducive to microbial growth, or that Wedl tunnelling does not occur post-fossilisation, when the organic and inorganic components of bone have mineralised or degraded. That no statistically significant differences in the severity of tunnelling across time scales were found in this study supports the hypothesis that that local conditions impact the degree of bioerosion.

Previous studies identified that the depth of tunnelling associated with cyanobacteria penetration can vary up to 200 - 400 µm [82], whilst the maximum depth of penetration from underwater caves extended up to 2250 µm. Time since burial may contribute to differences between bone analysed here and those in early stages of tunnelling, however, tunnelling is not a continuous process as shown by a peripheral degradation pattern on seven million years old bones [81, 82]. Differences in penetration depth is likely altered by surface bone loss through corrosion and flaking and exfoliation that reduces the observed penetration depth. Recording the presence of intact bone surfaces is thus critical in measuring degree of penetration.

A possible relationship between aquatic settings, crumbly surface textures, fungi and mosses, and bioerosion across the sub-periosteal regions have been previously identified [19], with some considering this a function of surface corrosion by other biotic agents [76]. A chalky sub-periosteal pocket of degradation and peripheral microboring was identified on some of our samples from underwater caves. These regions are associated with tunnelling but not the surface corrosion features linked to staining and biological attack. Tunnelling possibly associated with cyanobacteria thus will not produce surface modification and is only observed through histological analyses.

Whilst radial microfractures across the secondary osteon cement line are theoretically linked to early diagenesis in aquatic environments [36, 40, 83], they were not observed in this study. Time since deposition, or the process of mineralisation in water may play a larger role in producing the microfractures identified in fossils from aquatic environments [37, 38]. Here, neither a decadal or centennial timeframe produced these observations, nor have they been reproduced in other experimental studies [43].

The relationship between collagen preservation and aquatic environments is not well understood, with initial studies suggesting increased degradation underwater compared to on land [43]. Whilst birefringence cannot directly assess collagen preservation, it was consistently high in out samples in addition to ZooMs peptides successfully yielded from the preserved bones from dry and wet environments. This might support preservation of collagen structures up to 180 years after submergence based on our samples. In the thermally cool, and hydrologically stable underwater cave environment, collagen hydrolysis may be limited or slow, and thus the rate of water diffusion across bone may occur faster than collagen hydrolysis, preventing radial microfractures. Differences in vessel structures in bone across taxa, and density of primary and secondary Haversian bone, will further influence water diffusion rates across matrices [128]. Although Haversian bone systems were targeted for this study, some specimens contained laminar and longitudinal structures where the movement of water through the vascular network into the surrounding bone matrix is not impeded by cement lines.

### Cave zonation and taphonomic expressions

Cave systems do not present as a uniform or homogenous site. Karst systems are hydrologically zoned into the saturated (underwater) phreatic zone, the unsaturated (above water) vadose zone, and the epiphreatic interface [129]. Biological communities in caves are further controlled by light, graduating from the light entrance to dark twilight zone [8]. Areas within the cave can be considered taphonomically passive or active [130]: passive regions indicate a low degree of modification, and active regions indicate a high degree of modification (Fig 17).

**Fig 17.**
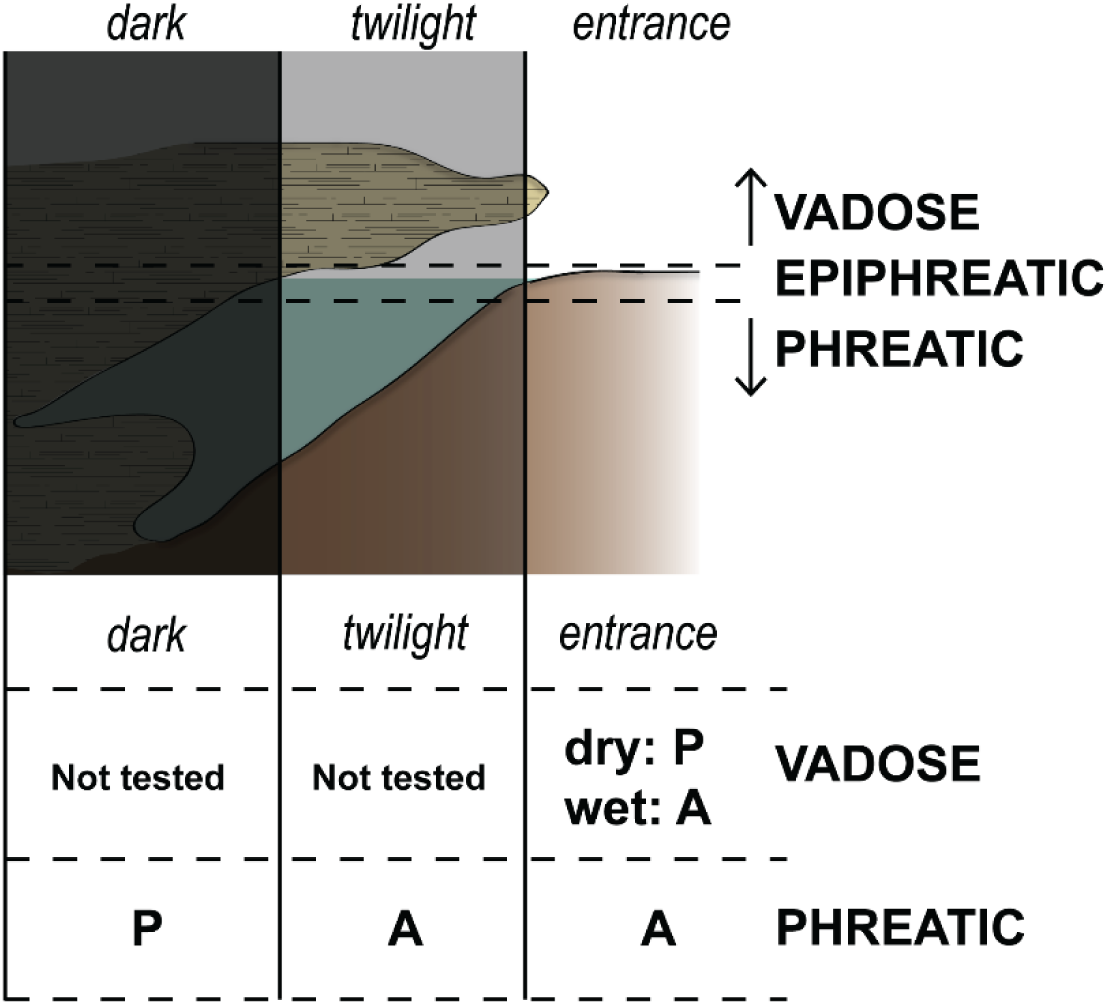
Passive (P) and active (A) regions of taphonomic modification across underwater caves accounting for light availability. The vadose entrance zone can shift between active and passive states depending on the availability of surface water [130]. Further testing across dark and twilight vadose zones is required for a comprehensive understanding of modifications associated with burial conditions.

Light availability in phreatic caves is here suggested to heavily influence bone modifications, shifting from a light, active entrance region to a dark, passive region of the cave system. Aquatic modifications, identified as an increase in surface corrosion, biotic black staining, and peripheral microboring by cyanobacteria, dominate across the entrance and twilight zones of the cave. Deeper specimens in darker regions of the cave presented with limited to no biotic modifications, although sample sizes are small. Light induced staining, corrosion and biota on bone surfaces shows that the ‘dark side’ of bones do not support biological growth. Limited Wedl-tunnelling in deep regions of submerged caves further supports a potential relationship between cyanobacteria and light in caves and deep maritime contexts [8, 79, 131, 132]. Cyanobacteria are photosynthetic prokaryotes that rely on light to grow and reproduce; however, studies highlight their adaptability to darkness [133]. Developing this model for phreatic system will rely on testing changes in oxygen availability, salinity, and hydraulic flow regimes across systems [101, 102, 114, 134].

An essential consideration for deep time assemblages in phreatic caves are water table fluctuations, and cave development that alters the light zones through time [9, 129]. Seasonal and eustatic modifications to the water table will shift hydrological zones, potentially modifying the burial environment to a passive system. A change from wet to dry burial environment may be indicated by the presence of delamination on bone surfaces. Mixed cyanobacteria and terrestrial bacterial tunnelling across bone microstructures can also suggest a mixed taphonomic history. These changes effectively act as reworking events, changing the burial environments and taphonomic histories.

## Conclusion

Underwater environments were characterised by well-preserved bones that show limited sorting by shape across talus cones. Over 68% of the wet assemblages retained over 80% of their original bone element portion compared to 44% in the dry assemblage, and breakage index distributions were significantly different between the two burial contexts where wet assemblages were less fragmented.

There were significant differences in bone surface modification frequencies between wet and dry burial conditions (Table 5). Whilst weathering scale did not separate the two conditions, parameters of the weathering scale did. Dry bone presented with significantly greater flaking and exfoliation and wet bone with more delamination. Wet conditions significantly increased the frequency of abrasion and physical modifications; however this is likely a result of the ‘soft’, chalky bone mineral associated with wet bones, predisposing them to collection damage. Both wet and dry assemblages presented with different types of corrosion, with etching occurring more often on dry bones, and surface corrosion on wet specimens. Certain types of etching were seen solely in wet (circular target etching) conditions. A significant association between surface corrosion, continuous black staining with sharp margins, and algae/biofilms was only observed on submerged samples. Frequency of surface corrosion was the only feature influenced by time, with older specimens having greater corrosion on their surfaces.

Bone microstructural bioerosion distinguished between wet and dry conditions, where wet bones presented with a pattern of cyanobacterial etching focused on the sub-periosteal peripheral margins, whilst dry bones were degraded by a range of bioeroders with no identified pattern of degradation. Time was not seen to influence the pattern of microstructural decay. However, a larger dataset would improve the statistical robusticity of tests across difference time scales and depths.

The interplay between light and water are key determinations of modification expression at the macro and histological levels. Changing light conditions across cave zones modify the biological agents within a submerged cave site, increasing activity and thus modifications across the entrance and twilight zone. Further testing of these zones will support reconstructing burial environments from bones deposited in phreatic cave systems.

## Supporting information

Supplementary BSM and Histotaphonomy Data

Supplementary Appendix Expanded Methods

## Acknowledgements

We thank the Boandik Aboriginal Corporation on whose land this work was conducted, the Department for Environment and Water (DEW) for land access permissions, the Cave Divers Association of Australia specifically Damian Bishop, Kelvyn Ball, Steve Trewavas, Hiro Yoshida, Ellyse Klein and Tanya Yarra for supporting this project, Jody Kruger for additional dive support, Tanya Smith, Kritim Dhakal, Jillian Huntley, Vikram Neelesh Vakil, and Carney Matherson for their support facilitating lab and instrument access, Daniel Kolarich and Arun Everest-Dass for assistance with the ZooMS analysis, and Linda Barry for work extracting isotope data to assess bone quality.

## Supporting information

**S1 Appendix. Expanded methods for Neotaphonomic characteristics of vertebrate site formation in underwater caves.**

**S1 Dataset. Bone surface modification frequency data, and histotaphonomy measurements and descriptions associated with wet and dry burial contexts across submerged caves.**

**S1 Fig.**
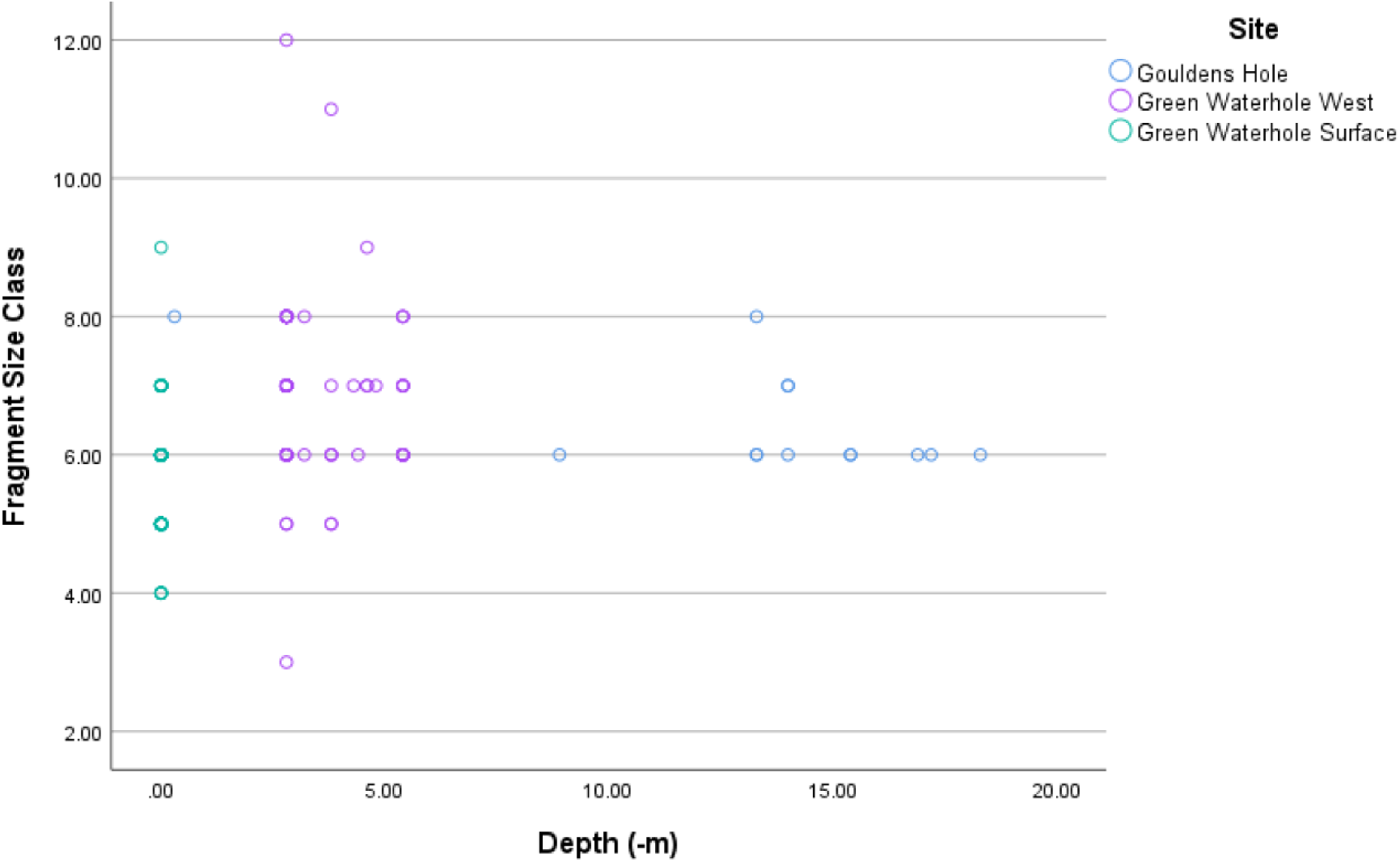
Distribution of bone size class [60] across depth (-m).

**S2 Fig.**
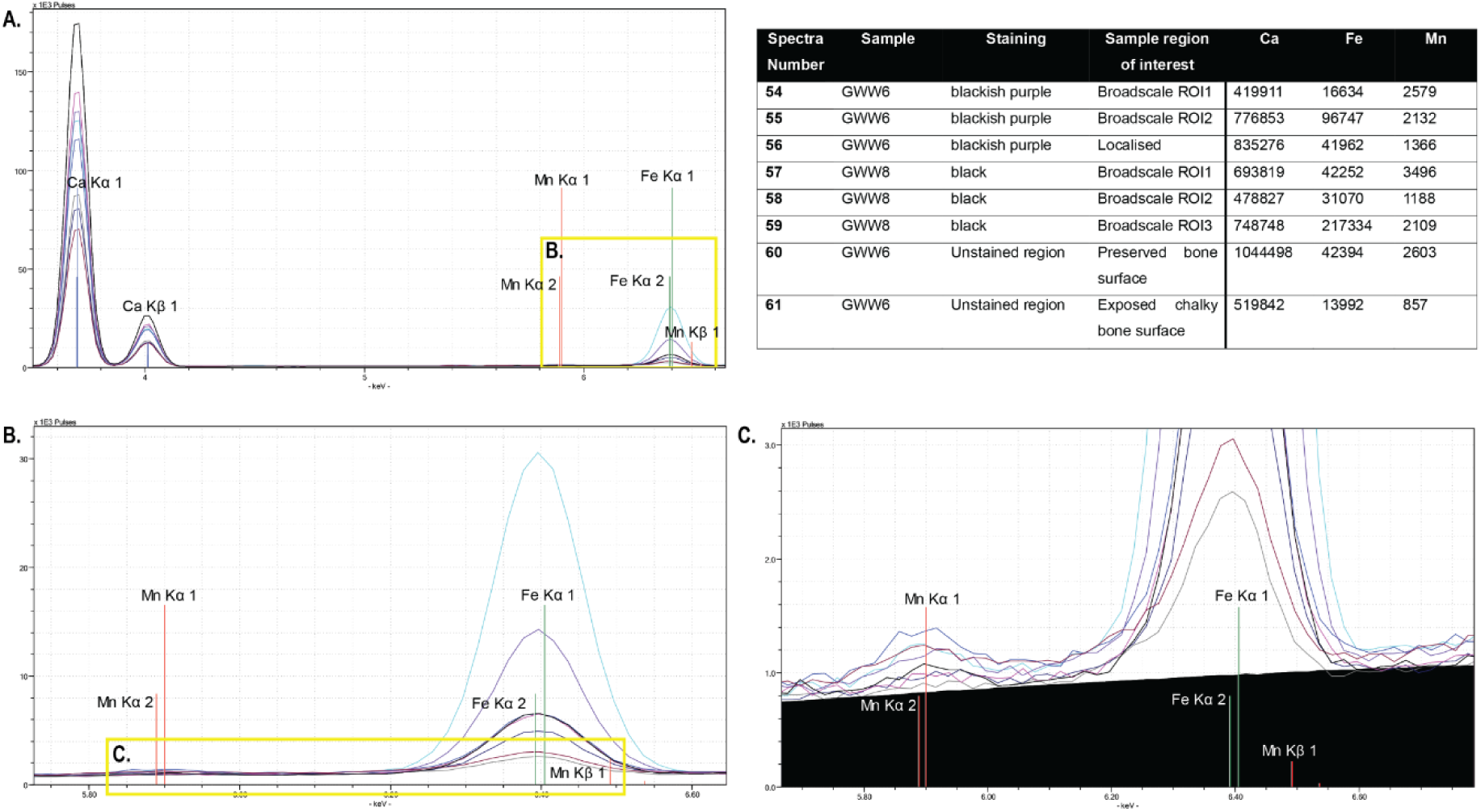
PXRF spectra with calcium (Ca), iron (Fe) and manganese (Mn) indicated. Associated spectral values and data provided in table. A: complete spectra; B: spectra highlighting Mn and Fe; C: focused view of Mn peaks.

**S3 Fig.**
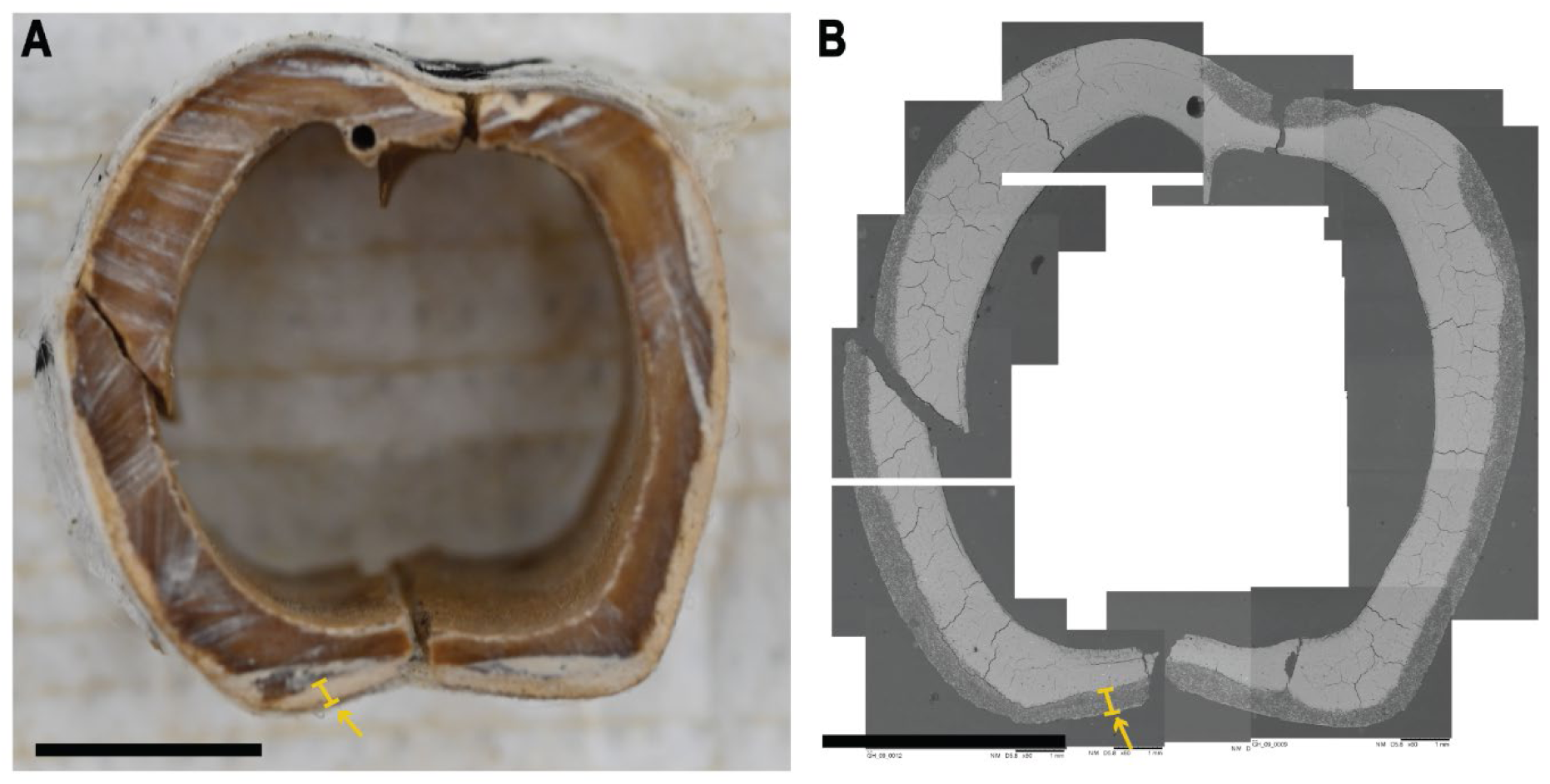
Cross-section of GH_09 sheep metatarsal comparing spatial distribution of chalky white texture (A) and cyanobacterial tunnelling (B). Yellow arrow and indicator identify the area that represents both a chalky degradation and tunnelling. Scale bars represent 5mm.

**S1 Table. ZooMS analysis and taxonomic ID across a sub-sample of taxon.**

**S2 Table. Radiocarbon dating results from ANSTO and ANU.**

## References

1. Stone TT, Dickel DN, Doran GH. The Preservation and Conservation of Waterlogged Bone from the Windover Site, Florida: A Compsrison of Methods. J. Field Archaeol. 1990;17(2):177-86.

2. González González AH, Sandoval CR, Mata AT, Sanvicente MB, Stinnesbeck W, Jeronimo Aviles O, et al. The Arrival of Humans on the Yucatan Peninsula: Evidence from Submerged Caves in the State of Quintana Roo, Mexico. Current Research in the Pleistocene. 2008;25:1–24.

3. Rosenberger AL, Godfrey LR, Muldoon KM, Gunnell GF, Andriamialison H, Ranivoharimanana L, et al. Giant subfossil lemur graveyard discovered, submerged, in Madagascar. J Hum Evol. 2015;81:83–7.

4. Newton CA. A Taphonomic and palaeoecological analysis of the Green Waterhole (5L81), a Submerged Late Pleistocene Bone deposit in the Lower Southeast of South Australia [Honours dissertation]: Adelaide: Flinders University; 1988.

5. Steadman DW, Franz R, Morgan GS, Albury NA, Kakuk B, Broad K, et al. Exceptionally well preserved late Quaternary plant and vertebrate fossils from a blue hole on Abaco, The Bahamas. Proc Natl Acad Sci U S A. 2007;104(50):19897–902.

6. Louys J. Practice and prospects in underwater palaeontology. Palaeontol. electron. 2018:1–14.

7. Walker MM, Louys J. Site formation processes and the taphonomy of vertebrate remains in underwater caves. Earth-Sci. Rev. 2024;256(104883):1–19

8. Kosznik-Kwasnicka K, Golec P, Jaroszewicz W, Lubomska D, Piechowicz L. Into the Unknown: Microbial Communities in Caves, Their Role, and Potential Use. Microorganisms. 2022;10(2).

9. Gillieson DS. Caves: processes, development, and management. 2nd ed. Hoboken: Wiley Blackwell; 2021.

10. Kowalewski M, Labarbera M. Actualistic Taphonomy: Death, Decay, and Disintegration in Contemporary Settings. PALAIOS. 2004;19(5):423–7.

11. Lyman RL. What Taphonomy Is, What it Isn’t, and Why Taphonomists Should Care about the Difference. J. Taphon. 2010;8(1):1–16.

12. Turner-Walker G. Early bioerosion in skeletal tissues: persistence through deep time. Neues Jahrbuch für Geologie und Paläontologie - Abhandlungen. 2012;265(2):165–83.

13. Keenan SW. From bone to fossil: A review of the diagenesis of bioapatite. Am. Mineral. 2016;101(9):1943–51.

14. Smith HE, Price GJ, Duval M, Westaway K, Zaim J, Rizal Y, et al. Taxonomy, taphonomy and chronology of the Pleistocene faunal assemblage at Ngalau Gupin cave, Sumatra. Quat. Int. 2021;603:40–63.

15. Haglund WD. Disappearance of Soft Tissue and the Disarticulation of Human Remains from Aqueous Environments. J. Forensic Sci. 1993;38(4):806–15.

16. Reed EH. Decomposition and Disarticultion of Kangaroo Carcasses in Caves at Naracoorte, South Australia. J. Taphon. 2009;7(4):265–84.

17. Syme CE, Salisbury SW. Patterns of aquatic decay and disarticulation in juvenile Indo-Pacific crocodiles (Crocodylus porosus), and implications for the taphonomic interpretation of fossil crocodyliform material. Palaeogeogr., Palaeoclimatol., Palaeoecol. 2014;412:108–23.

18. Andrews P. Owls, Caves and Fossils. Chicago: The University of Chicago Press; 1990 1990.

19. Fernández-Jalvo Y, Andrews P, Pesquero D, Smith C, Marín-Monfort D, Sánchez B, et al. Early bone diagenesis in temperate environments: Part I: Surface features and histology. Palaeogeogr., Palaeoclimatol., Palaeoecol. 2010;288(1-4):62–81.

20. Brian CK. The Hunters or th Hunted? An Introduction to African Cave Taphonomy. Chicago, London: University of Chicago Press; 1981.

21. Fernández-Jalvo Y, Andrews P. Atlas of Taphonomic Identifications. Delson E, Sargis, EJ. editors. Dordrecht Heidelberg New York London: Springer; 2016.

22. Griffith SJ. Aquatic Bone Taphonomy: Forensic and Archaeological Implications for the Interpretation of Submerged Bone [PhD dissertation]. Southampton: University of Southampton; 2017.

23. Littleton J. Taphonomic Effects of Erosion on Deliberately Buried Bodies. J. Archaeol. Sci. 2000;27:5–18.

24. Fernández-Jalvo Y, Andrews P. Experimental Effects of Water Abrasion on Bone Fragments. J. Taphon. 2003;1(3):145–61.

25. Cartajena I, López P, Carabias D, Morales C, Vargas G, Ortega C. First evidence of an underwater Final Pleistocene terrestrial extinct faunal bone assemblage from Central Chile (South America): Taxonomic and taphonomic analyses. Quat. Int. 2013;305:45–55.

26. Cartajena I, Marin F, Carabias D, Pavez J, Yrarrázaval S. How Underwater Bone Taphonomy Looks Like? Reevaluating Taphonomic Indicators in a Late Pleistocene Drowned Terrestrial Site, Chile 14th International Conference for Archaeozoology; 2023 August 8-12; Cairns, Australia

27. Guareschi EE, Schönberg CHL, Magni PA, Tobe SS, Nicholls PK, Turner-Walker G. Marine sponge bioerosion in the forensic taphonomy of terrestrial bone. Quat. Int. 2023;660:84–94.

28. Pokines JT, Higgs N. Macroscopic taphonomic alterations to human bone in marine environments. J. Forensic Identif. 2015;65(6):953–84.

29. Brönnimann D, Portmann C, Pichler SL, Booth TJ, Röder B, Vach W, et al. Contextualising the dead – Combining geoarchaeology and osteo-anthropology in a new multi-focus approach in bone histotaphonomy. J. Archaeol. Sci. 2018;98:45–58.

30. Bell LS. Histotaphonomy. In: Crowder C, Stout S, editors. Bone Histology: an Anthropological Perspective. Boca Raton, Fla: CRC Press; 2012. p. 241-52.

31. Jans MME. Microscopic Destruction of Bone. In: Pokines JT, Symes SA, editors. Manual of Forensic Taphonomy. Boca Raton: CRC Press; 2014. p. 19-35.

32. Jans MME. Histological Characterisation of Diagenetic Alteration of Archaeological Bone. Amsterdam: Institute for Geo and Bio-archaeology, Vrije Universiteit; 2005.

33. Hedges REM. Bone diagenesis: an overview of processes. Archaeometry. 2002;44(3):319–28.

34. Schotsmans EMJ, Stuart BH, Stewart TJ, Thomas PS, Miszkiewicz JJ. Unravelling taphono-myths. First large-scale study of histotaphonomic changes and diagenesis in bone from modern surface depositions. PLoS One. 2024;19(9):e0308440.

35. Eriksen AMH, Nielsen TK, Matthiesen H, Caroe C, Hansen LH, Gregory DJ, et al. Bone biodeterioration-The effect of marine and terrestrial depositional environments on early diagenesis and bone bacterial community. PLoS One. 2020;15(10):e0240512.

36. Pfretzschner H-U. Fossilization of Haversian bone in aquatic environments. Comptes Rendus Palevol. 2004;3(6-7):605–16.

37. Pfretzschner H-U, Tütken T. Rolling bones – Taphonomy of Jurassic dinosaur bones inferred from diagenetic microcracks and mineral infillings. Palaeogeogr., Palaeoclimatol., Palaeoecol. 2011;310(1-2):117–23.

38. Marin-Monfort D, de Santisteban C, Garrone M, Montalvo CI, Fernández-Jalvo Y, Fernández FJ, et al. Histotaphonomy of a Pleistocene megamammal assemblage from Argentine Pampas. J. South Am. Earth Sci. 2023;129.

39. Guareschi EE, Haig DW, Tobe SS, Nicholls PK, Magni PA. Foraminifera—A new find in the microtaphonomical characterization of bones from marine archaeological excavations. Int. J. Osteoarchaeol. 2021;31(6):1270–5.

40. Pfretzschner HU. Microcracks and fossilization of Haversian bone. Neues Jahrbuch für Geologie und Paläontologie - Abhandlungen. 2000;216(3):413.

41. Guareschi EE, Tobe SS, Nicholls PK, Magni PA. Taphonomy and Diagenesis of Human Bone in Underwater Archaeology: A Review of the Current Status and the Proposal of Post-Mortem Submersion Interval (PMSI) as a Potential Forensic Application. J. Marit. Archaeol. 2021;16(1):57–75.

42. Turner-Walker G. Light at the end of the tunnels? The origins of microbial bioerosion in mineralised collagen. Palaeogeogr., Palaeoclimatol., Palaeoecol. 2019;529:24–38.

43. Guareschi EE, Nicholls PK, Tobe SS, Magni PA. Taphonomy and diagenesis of submerged bone: an experimental approach. Forensic Sci. Int. 2025.

44. Horne P. Project Report No. 1: “Fossil Cave” - 5L81 Underwater Palaeontoloigical and Surveying Project 1987-1988. South Australian Speleological Society Inc.; 2006.

45. Mather EK, Lee MSY, Fusco DA, Hellstrom J, Worthy TH. Pleistocene raptors from cave deposits of South Australia, with a description of a new species of Dynatoaetus (Accipitridae: Aves): morphology, systematics and palaeoecological implications. Alcheringa: An Australasian Journal of Palaeontology. 2023:1–32.

46. Reed EH, Bourne SJ. Pleistocene Fossil Vertebrate Sites of the South East Region of South Australia. Trans. R. Soc. S. Aust. 2000;124:61–90.

47. Horne P. Project Report No. 2: Gouldens Hole - 5L8 - mapping project, 1987/1988. Hove, SA: South Australian Underwater Speleological Society; 2009.

48. Department of Environment and Water. Limestone Coast Prescribed Areas 2020–21 water resources assessment DEW Technical Note 2022/13. Adelaide: Government of South Australia, Department for Environment and Water; 2023.

49. Wilcken K, Hotchkis M, Levchenko V, Fink D, Hauser T, Kitchen R. From carbon to actinides: A new universal 1MV accelerator mass spectrometer at ANSTO. Nucl. Instr. Meth. Phys. Res. B. 2015;361:133–8.

50. Fallon SJ, Fifield LK, Chappell JM. The next chapter in radiocarbon dating at the Australian National University: Status report on the single stage AMS. Nucl. Instr. Meth. Phys. Res. B. 2010;268(7-8):898–901.

51. Wood R, Esmay R, Usher E, Fallon S. SAMPLE PREPARATION METHODS USED AT THE AUSTRALIAN NATIONAL UNIVERSITY RADIOCARBON FACILITY. Radiocarbon. 2023;65(2):573–89.

52. Suess HE. Radiocarbon concentration in modern wood. Science. 1955;122(3166):415–7.

53. Hua Q. Radiocarbon: A chronological tool for the recent past. Quat. Geochronol. 2009;4:378–390.

54. Unknown Author. Early History of Mount Gambier. The Border Watch. 1946 Nov 30:8.

55. Hill LR. Mount Gambier: the city around a cave. Leabrook, SA: Investigator Press; 1972.

56. MacGillivary L. We have found our paradise: The South-East squattocracy, 1840–1870. Journal of the Historical Society of South Australia. 1989;17:25–38.

57. Uno KT, Quade J, Fisher DC, Wittemyer G, Douglas-Hamilton I, Andanje S, et al. Bomb-curve radiocarbon measurement of recent biologic tissues and applications to wildlife forensics and stable isotope (paleo)ecology. Proc Natl Acad Sci U S A. 2013;110(29):11736–41.

58. Hua Q, Turnbull JC, Santos GM, Rakowski AZ, Ancapichún, S.,, De Pol-Holz R, Hammer, S.,, Lehman SJ, et al. Atmospheric radiocarbon for the period 1950-2019. Radiocarbon. 2022;64(4):723–45.

59. Fillios M, Blake N. Animal Bones in Australian Archaeology: A Field Guide to Common Native and Introduced Species. Sydney: Sydney University Press; 2015.

60. O’Connor TP. The Archaeology of Animal Bones. Stroud: Sutton Publishing Ltd; 2008.

61. Andrews P, Molleson T, Boz B. The Human Burials at Çatalhöyük. In: Hodder I, editor. Inhabiting Çatalhöyük reports from the 1995-99 seasons. McDonald Institute for Archaeological Research and British Institute of Archaeology at Ankara Cambridge, London; 2005. p. 261.

62. Blob RW, Fiorillo AR. The significance of vertebrate microfossil size and shape distributions for faunal abundance reconstructions: a Late Cretaceous example. Paleobiology. 1996;22(3):422–35.

63. Voorhies MR. Taphonomy and Population Dynamics of an early Pliocene Vertebrate Fauna, Knox County, Nebraska. Laramie, Wyoming University of Wyoming; 1968.

64. Lyman LR. Available Meat from Faunal Remains: A Consideration of Techniques. Am. Antiq. 1979;44(3):536–46.

65. Lyman RL. Quantitative paleozoology. Cambridge; New York: Cambridge University Press; 2008 2008.

66. Lyman RL. Vertebrate Taphonomy. Cambridge: Cambridge University Press; 1994.

67. Dobney K, Rielly K. A method for recording archaeological animal bones: the use of diagnostic zones. Circaea. 1988;5(2):79–96.

68. Fiorillo AR. Taphonomy of Hazard Homestead Quarry (Ogallala Group), Hitchcock County, Nebraska. Rocky Mountain Geology. 1988;26(2):57–97.

69. Villa P, Mahieu E. Breakage patterns of human long bones. J. Hum. Evol. 1991;21:27–48.

70. Huntley J. Taphonomy Or Paint Recipe: In situ portable x-ray fluorescence analysis of two anthropomorphic motifs from the Woronora Plateau, New South Wales. Aust. Archaeol. 2016;75(1):78–94.

71. Huntley J, Aubert M, Ross J, Brand HEA, Morwood MJ. One Colour, (at Least) Two Minerals: A Study of Mulberry Rock Art Pigment and a Mulberry Pigment ‘Quarry’ from the Kimberley, Northern Australia. Archaeometry. 2015;57(1):77–99.

72. Behrensmeyer AK. Taphonomic and ecologic information from bone weathering. Paleobiology. 1978;4(2):150–62.

73. Walker MM, Louys J, Herries AIR, Price GJ, Miszkiewicz JJ. Humerus midshaft histology in a modern and fossil wombat. Aust. Mammal. 2021;43(1).

74. Turner-Walker G, Nielsen-Marsh CM, Syversen U, Kars H, Collins MJ. Sub-micron spongiform porosity is the major ultra-structural alteration occurring in archaeological boneInt. J. Osteoarchaeol. 2002;12(6):407–14.

75. Hackett CJ. Microscopial Focal Destruction (Tunnels) in Exhumed Human Bones. Med Sci Law. 1981;21(4):243–65.

76. Turner-Walker G. Diagenetic Alternations to Vertebrate Mineralized Tissues - A Critical Review. In: Pollard AM, Armitage RA, Makarewicz CA, editors. Handbook of Archaeological Sciences. 2^nd^ ed. Milton, QLD:John Wiley & Sons; 2023. p. 1117–55.

77. Hackett CJ. Microscopical Focal Destruction (Tunnels) in Exhumed Human Bones. Med Sci Law. 1981;21(4):243–65.

78. Trueman CN, Martill DM. THE LONG-TERM SURVIVAL OF BONE: THE ROLE OF BIOEROSION. Archaeometry. 2002;44(3):371–82.

79. Bell LS, Elkerton A. Unique marine taphonomy in human skeletal material recov-ered from the medieval warship Mary Rose. Int. J. Osteoarchaeol. 2007;18:523–35.

80. Wedl C. Über einen im Zahnbeim und Knochen keimenden Pilz. Sitzungsberichte der Kaiserlichen Akademie der Wissenschaften, mathematisch-Naturwissenschaftliche Classe. 1865;50.

81. Pesquero MD, Ascaso C, Alcalá L, Fernández-Jalvo Y. A new taphonomic bioerosion in a Miocene lakeshore environment. Palaeogeogr., Palaeoclimatol., Palaeoecol. 2010;295(1-2):192–8.

82. Pesquero MD, Bell LS, Fernández-Jalvo Y. Skeletal modification by microorganisms and their environments. Hist. Biol. 2017;30(6):882–93.

83. Pfretzschner HU. Collagen gelatinization: the key to understand early bone-diagenesis. Palaeontographica Abteilung A: Palaozoologie. 2006;278:135–48.

84. Hedges REM, Millard AR. Measurements and relationshops of diagenetic alteration of bone from three archaeological sites. J. Archaeol. Sci. 1995;22:201–9.

85. Prittard CR. Reconstructing the breed and introduction of South Australia’s first sheep from ancient DNA [Honours dissertation]. Nathan, QLD: Griffith University; 2024.

86. Stathopoulou E, Phoca Cosmetatou N, Theodoropoulou T, Mallouchou M, Margariti E, Psycharis V. Origin of archaeological black bones within a waterlogged context: A multidisciplinary approach. Palaeogeogr., Palaeoclimatol., Palaeoecol. 2019;534.

87. Hedges REM, Millard AR, Pike AWG. Measurements and Relationships of Diagenetic Alteration of Bone from Three Archaeological Sites. J. Archaeol. Sci. 1995;22(2):201–9.

88. Philippsen B. The freshwater reservoir effect in radiocarbon dating. Herit. Sci. 2013;1(24):1–19.

89. Keaveney EM, Reimer PJ. Understanding the variability in freshwater radiocarbon reservoir offsets: a cautionary tale. J. Archaeol. Sci. 2012;39(5):1306–16.

90. Zazzo A, Saliège JF. Radiocarbon dating of biological apatites: A review. Palaeogeogr., Palaeoclimatol., Palaeoecol. 2011;310(1-2):52–61.

91. Chatters JC, Kennett DJ, Asmerom Y, Kemp BM, Polyak V, Nava Blank A, et al. Late Pleistocene Human Skeleton and mtDNA Link Paleoamericans and Modern Native Americans. Sci. Rep. 2014;344:750–4.

92. Zazzo A. Bone and enamel carbonate diagenesis: A radiocarbon prospective. Palaeogeogr., Palaeoclimatol., Palaeoecol. 2014;416:168–78.

93. Lewis ID. Interpreting the Mount Gambier cenotes (sinkholes) within the Kanawinka Geopark. ACKMA Journal. 2007;17:75–82.

94. Eberhard R. Conservation Column No. 2: Use and abuse of sinkholes. ACKMA Journal. 2005;61:22–4.

95. Aslan A, Behrensmeyer AK. Taphonomy and Time Resolution of Bone Assemblages in a Contemporary Fluvial System: The East Fork River, Wyoming. PALAIOS. 1996;11(5):411–21.

96. Frostick L, Reid I. Taphonomic significance of sub-aerial transport of vertebrate fossils on steep semi-arid slopes. Lethaia. 1983;16:157–64.

97. Boaz NT, Behrensmeyer AK. Hominid taphonomy: transport of human skeletal parts in an artificial fluviatile environment. Am J Phys Anthropol. 1976;45(1):53–60.

98. Reed EH. In Situ Taphonomic Investigation of Pleistocene Large Mammal Bone Deposits from The Ossuaries, Victoria Fossil Cave, Nacoorte, South Australia. Helictite. 2006;39(1):5–15.

99. Reed EH. Vertebrate Taphonomy of Large Mammal Bone Deposits, Naracoorte Caves World Heritage Area. South Australia [PhD dissertation]: Adelaide: The Flinders University of South Australia; 2003.

100. Fernández-Jalvo Y, Denys C, Andrews P, Williams T, Dauphin Y, Humphrey L. Taphonomy and palaeoecology of Olduvai Bed-I (Pleistocene, Tanzania). J. Hum. Evol. 1998;34:137–72.

101. Evans T. Fluvial Taphonomy. In: Pokines JT, Symes SA, editors. Manual of Forensic Taphonomy. Boca Raton: CRC Press; 2014. p. 115–41.

102. Higgs ND, Pokines JT. Marine Environmental Alterations to Bone. In: Pokines JT, Symes SA, editors. Manual of Forensic Taphonomy. Boca Raton: CRC Press; 2014. p. 143–79.

103. Pokines JT, Faillace K, Berger J, Pirtle D, Sharpe M, Curtis A, et al. The effects of repeated wet-dry cycles as a component of bone weathering. J. Archaeol. Sci: Reports. 2018;17:433–41.

104. Stinnesbeck SR, Frey E, Olguín JA, Stinnesbeck W, Zell P, Mallison H, et al. Xibalbaonyx oviceps, a new megalonychid ground sloth (Folivora, Xenarthra) from the Late Pleistocene of the Yucatán Peninsula, Mexico, and its paleobiogeographic significance. PalZ. 2017;91(2):245–71.

105. Pokines JT, Baker JE. Effects of Burial Environment on Osseous Remains. In: Pokines JT, Symes SA, editors. Manual of Forensic Taphonomy. Boca Raton: CRC Press; 2014. p. 74–114.

106. Entwisle TJ, Skinner S, Lewis SH, Foard HJ. Algae of Australia: Batrachospermales, Thoreales, Oedogoniales and Zygnemaceae. Clayton: Australian Biological Resources Study, CSIRO Publishing; 2007.

107. McCarthy PM, editor. Flora of Australia Volume 51 (Mosses). Canberra & Melbourne: Australia Biological Resource Study, CSIRO; 2006.

108. Pesquero MD, Fernández-Jalvo Y. Bioapatite to calcite, an unusual transformation seen in fossil bones affected by aquatic bioerosion. Lethaia. 2014;47(4):533–46.

109. Collareta A, Tsai C-H, Coletti G, Bosselaers M. Thatchtelithichnus on a Pliocene grey whale mandible and barnacles as possible tracemakers. Neues Jahrbuch für Geologie und Paläontologie - Abhandlungen. 2021;302(1):53–61.

110. Kip N, van Veen JA. The dual role of microbes in corrosion. ISME J. 2015;9(3):542–51.

111. Mulec J, Kosi G, Vrhovsek D. Characterization of Cave Aerophytic Algal Communities and Effects of Irradiance Levels on Production of Pigments. Journal of Cave and Karst Studies. 2008;70(1).

112. Cappitelli F, Salvadori O, Albanese D, Villa F, Sorlini C. Cyanobacteria cause black staining of the National Museum of the American Indian Building, Washington, DC, USA. Biofouling. 2012;28(3):257–66.

113. Maestri C, Hébert RL, Di Martino P. Biofilm associated with pigmented areas on a waterproofing coating surface. AIMS Microbiology. 2025;11(1):74–86.

114. Brankovits D, Pohlman JW, Niemann H, Leigh MB, Leewis MC, Becker KW, et al. Methane-and dissolved organic carbon-fueled microbial loop supports a tropical subterranean estuary ecosystem. Nat. Commun. 2017;8(1):1835.

115. Griffith SJ, Thompson CEL, Thompson TJU, Gowland RL. Experimental abrasion of water submerged bone: The influence of bombardment by different sediment classes on microabrasion rate. J. Archaeol. Sci: Reports. 2016;10:15–29.

116. Thompson CEL, Ball S, Thompson TJU, Gowland R. The abrasion of modern and archaeological bones by mobile sediments: the importance of transport modes. J. Archaeol. Sci. 2011;38(4):784–93.

117. d’Errico F, Giacobini G, Puech PF. Varnish replicas: A new method for the study of worked bone surfaces. International Journal of Skeletal Research. 1984;9–11:29-51.

118. Christensen AM, Myers SW. Macroscopic observations of the effects of varying fresh water pH on bone. J Forensic Sci. 2011;56(2):475–9.

119. López-González F, Grandal-d’Anglade A, Ramón Vidal-Romaní J. Deciphering bone depositional sequences in caves through the study of manganese coatings. J. Archaeol. Sci. 2006;33(5):707–17.

120. Fernández-Jalvo Y, Monfort MDM. Experimental taphonomy in museums: Preparation protocols for skeletons and fossil vertebrates under the scanning electron microscopy. Geobios. 2008;41(1):157–81.

121. Barron-Ortiz CI, Sawchuk MR, Li C, Jass CN. Conservation of subfossil bones from a lacustrine setting: uncontrolled and controlled drying of Late Quaternary vertebrate remains from Cold Lake, Western Canada. Collection Forum. 2018;32(1):1–13.

122. Eric M, Puhar, E.G., Jaklic, A. and Solina, F. The Necessity of Changing the Methodology of Preserving Waterlogged Wooden Objects: The Case of a Palaeolithic Wooden Point from the Ljubljanica River. Skyllis. 2018;2:174–85.

123. Grant T. Conservation of Wet Faunal Remains: Bone, Antler, and Ivory. 2007.

124. Wescott DJ. Postmortem change in bone biomechanical properties: Loss of plasticity. Forensic Sci Int. 2019;300:164–9.

125. Turner-Walker G, Jans M. Reconstructing taphonomic histories using histological analysis. Palaeogeogr., Palaeoclimatol., Palaeoecol. 2008;266(3-4):227–35.

126. Yoshino M, Kimijima T, Miyasaka S, Sato H, Seta S. Microscopical study on estimation of time since death in skeletal remains. Forensic Sci. Int. 1991;49:143–58.

127. Sullivan CA, Keenan SW. Experimental dissolution of fossil bone under variable pH conditions. PLoS One. 2022;17(10):e0274084.

128. Francillon-Vieillot H, de Buffrénil V, Castanet JD, Géraudie J, Meunier FJ, Sire JY, et al. Microstructure and mineralization of vertebrate skeletal tissues. In: Carter JG, editor. Skeletal Biomineralization: Patterns, Processes and Evolutionary Trends. New York: Van Nostrand Reinhold; 1990. p. 471–530.

129. Ford DC, Williams PW. Karst Geomorphology and Hydrology. London, England: Unwin Hyman; 2007.

130. Woodward JC, Goldberg P. The sedimentary records in Mediterranean rockshelters and caves: Archives of environmental change. Geoarchaeology. 2001;16(4):327–54.

131. Cennamo P, Marzano C, Ciniglia C, Pinto G, Cappelletti P, Caputo P, et al. A survey of the algal flora of anthropogenic caves of Campi Flegrei (Naples, Italy) archeological district. Journal of Cave and Karst Studies. 2012;74(3):243–50.

132. Popović S, Nikolić N, Jovanović J, Predojević D, Trbojević I, Manić L, et al. Cyanobacterial and algal abundance and biomass in cave biofilms and relation to environmental and biofilm parameters. Int. J. Speleol. 2019;48(1):49–61.

133. Hood RD, Higgins SA, Flamholz A, Nichols RJ, Savage DF. The stringent response regulates adaptation to darkness in the cyanobacterium Synechococcus elongatus. Proc Natl Acad Sci U S A. 2016;113(33):E4867–76.

134. Pokines JT, Nicholas DH. Marine Environmental Alterations to Bone. In: Pokines JT, L’abbé E, Symes SA, editors. Manual of Forensic Taphonomy. 2nd Edition ed. Boca Raton: CRC Press; 2021. p. 193–249.

