## Supplementary Table 2 for "Neotaphonomic characteristics of vertebrate site formation in underwater caves"

**S2 Table. Radiocarbon dating results from ANSTO and ANU**

| Lab | Lab Sample Code | Sample ID | Depth (m) | Taxa | Element | % yield | δ^13^C  (‰) | δ^15^N  (‰) | C:N | F14C  ± 1σ | ^14^C age (BP)  ± 1σ | Calibrated Age Ranges (cal AD (pr%)) | Time period | Defined  Age Period |
| --- | --- | --- | --- | --- | --- | --- | --- | --- | --- | --- | --- | --- | --- | --- |
| ANU | SANU-68813 | GH_01 | -10.1 | *Ovis aries* | Cranium | 14.04 | -20.88 | 8.22 | 2.97 | 0.9770 ± 0.0016 | 184 ± 18 | 1671 - 1712 (29.18) 1718 - 1744 (16.66) 1753 - 1765 (4.16) 1772 - 1781 (2.49) 796 - 1814 (12.73) 1834 - 1890 (21.54) 1923 - 1949* (13.22) | Centennial | 1841-1955 |
| ANU | SANU-70526 | GH_01 repeat | -10.1 | *Ovis aries* | Cranium | As above | As above | As above | As above | 0.9780 ± 0.0014 | 181 ± 16 | 1673 - 1712 (29.49) 1718 - 1740 (15.71) 1755 - 1764 (02.03) 1774 - 1779 (01.24) 1798 - 1813 (12.27) 1834 - 1890 (24.15) 1923 - 1949* (15.11) | Centennial | 1841-1955 |
| ANU | SANU-68814 | GH_02 | -10.1 | *Ovis aries* | Cranium | 7.99 | -20.18 | 8.65 | 2.98 | 0.9790 ± 0.0015 | 174 ± 17 | 1674 - 1737 (38.46) 1758 - 1761 (0.41) 1799 - 1817 (9.50) 1831 - 1892 (33.08) 1922 - 1949* (18.55) | Centennial | 1841-1955 |
| ANU | SANU-70527 | GH_02 repeat | -10.1 | *Ovis aries* | Cranium | As above | As above | As above | As above | 0.9800 ± 0.0014 | 159 ± 16 | 1687 - 1730 (27.70) 1806 - 1894 (52.00) 1921 - 1949* (20.30) | Centennial | 1841-1955 |
| ANU | SANU-76430 | GH_04 | -13.3 | *Macropus* sp. | Rib | 12.75 | -9.70 | 2.68 | 3.10 | 0.9774 ± 0.0021 | 184 ± 22 | 1671 - 1748 (42.72) 1749 - 1767 (05.45) 1771 - 1782 (03.35) 1796 - 1815 (11.69) 1833 - 1892 (23.52) 1922 - 1949* (13.26) | Centennial | 1841-1955 |
| ANU | SANU-76405 | GH_06 | -18.3 | Indet | Vertebra | 8.96 | -22.05 | 6.25 | 3.16 | 0.9797 ± 0.0020 | 165± 21 | 1677 – 1735 (29.6) 1801 - 1897 (46.7) 1905 - cal AD (19.2) | Centennial | 1841-1955 |
| ANU | SANU-76407 | GH_09 | -15.4 | *Ovis aries* | Metatarsal | 8.96 | -22.05 | 6.25 | 3.16 | 0.9885 ± 0.0020 | 93 ± 21 | 1705 - 1721 (4.5) 1811 - 1838 (19.8) 1849 - 1868 (4.6) 1878 - 1929 (66.5) | Centennial | 1841-1955 |
| ANU | SANU-76431 | GH_10 | -14 | Ovicaprid | Metatarsal | 10.10 | -13.13 | 5.33 | 3.13 | 0.9801 ± 0.0022 | 161 ± 23 | 1678 - 1733 (28.67) 1802 - 1896 (50.38) 1903 - 1915 (02.57) 1918 - 1949* (18.38) | Centennial | 1841-1955 |
| ANU | SANU-76432 | GH_16 | -16.9 | *cf. Macropus* sp. | Vertebra | 12.78 | -9.65 | 3.31 | 3.14 | 0.7971 ± 0.0022 | 1821 ± 27 | 140 - 153 (2.50) 165 - 177 (2.15) 202 - 341 (94.57) 353 - 359 (0.79) | Millennial | 140-359 |
| ANU | SANU-76433 | GH_18 | -0.3 | *Macropus* sp*.* | Tibia | 11.24 | -8.32 | 2.05 | 3.14 | 0.9679 ± 0.0022 | 262 ± 23 | 1640 - 1675 (49.00) 1736 - 1799 (51.00) | Centennial | 1841-1955 |
| ANU | SANU-76435 | GH_19 | -8.9 | Ovicaprid | Metatarsal | 11.98 | -22.34 | 7.28 | 3.16 | 0.9838 ± 0.0022 | 131 ± 23 | 1697 - 1724 (16.06) 1810 - 1872 (38.24) 1874 - 1947 (45.70) | Centennial | 1841-1955 |
| ANU | SANU-76436 | GH_22 | -17.2 | *Bos sp.* | Vertebra | 12.04 | -21.92 | 4.98 | 3.11 | 0.9785 ± 0.0022 | 175 ± 23 | 1672 - 1742(36.62) 1754 - 1765 (2.09) 1773 - 1780 (1.38) 1797 - 1894 (43.61) 1920 - 1949 (16.30) | Centennial | 1841-1955 |
| ANU | SANU-75425 | GH_23 | -10.1 | *Canis lupus dingo* | Mandible | 11.51 | -16.90 | 7.98 | 3.15 | 0.895 ± 0.0021 | 889 ± 24 | 1155 - 1230(92.34) 1248 - 1267 (07.66) | Centennial | 1155-1267 |
| ANU | SANU-76437 | GH_24 | surface | *Ovis aries* | Cranium | 7.70 | -21.42 | 6.00 | 3.18 | 0.9739 ± 0.0024 | 213 ± 24 | 1659 - 1699 (23.10) 1722 - 1811 (72.93) 1837 - 1845 (1.12) 1869 - 1876 (1.02) 1928 - 1938 (1.09) 1944 - 1949* (0.73) | Centennial | 1841-1955 |
| ANU | SANU-76521 | GH_26 | -10 | *Canis familiaris* | Cranium | 12.31 | -16.95 | 8.80 | 3.15 | 0.9513 ± 0.0022 | 401 ± 23 | 1456 - 1513 (51.61) 1544 - 1625 (48.39) | Centennial | 1456-1625 |
| ANU | SANU-76520 | GH_27 | -10 | *Trichosorus vulpecta* | Cranium | 8.35 | -20.76 | 3.18 | 3.22 | 0.9731 ± 0.0023 | 219 ± 24 | 1654 - 1696 (22.81) 1724 - 1810 (76.40) 1840 - 1842 (0.31) 1872 - 1873 (0.21) 1947 - 1949 (0.26) | Centennial | 1841-1955 |
| ANU | SANU-76519 | GH_28 | -10 | *Macropus* sp*.* | Cranium | 6.08 | -17.54 | 11.94 | 3.17 | 0.8254 ± 0.0019 | 1542 ± 24 | 529 - 637 (100) | Millennial | 529-637 |
| ANSTO | OZBP59 | GWS_47 | Surface | *Ovis aries* | Metatarsal | 7.85 | -20.5 | 5.8 | 3.2 | 0.9749 ± 0.0032 | 200 ± 30 | 1663 - 1710 (22.6) 1719 - 1812 (57.1) 1835 - 1861 (4.4) 1866 - 1880 (3.1) 1925 - 1942 (3.6) 1943 - 1945 (0.4) 1946 - 1955 (4.3) | Centennial | 1841-1955 |
| ANU | SANU-76507 | GWS_60 | Surface | *Bovidae* | Tibia | 12.94 | -21.90 | 5.84 | 3.09 | 0.9834 ± 0.0021 | 135 ± 22 | 1697 - 1725 (16.79) 1809 - 1872 (38.71) 1873 - 1947 (44.50) | Centennial | 1841-1955 |
| ANSTO | OZBU47 | GWS_66 | Surface | *Ovis aries* | Femur | 11.6 | -18.3 | 6.4 | 3.3 | 0.9893 ± 0.0030 | 90 ± 25 | 1701 - 1721 (5.7) 1811 - 1837 (18.9) 1848 - 1868 (5.4) 1878 - 1927 (58.8) 1939 - 1947 (4.1) 1954 - 1956 (2.5) | Centennial | 1841-1955 |
| ANSTO | OZBU46 | GWS_69 | Surface | *Ovis aries* | Metatarsal | 10.69 | -18.5 | 6.3 | 3.3 | 0.9842 ± 0.0029 | 130 ± 25 | 1697 - 1724 (14.6) 1810 - 1871 (35.7) 1874 - 1947 (44.8) 1948 - 1949 (0.2) 1954 - 1955 (0.2) | Centennial | 1841-1955 |
| ANU | SANU-76512 | GWS_86 | Surface | Ovicaprid | tibia | 8.84 | -22.03 | 7.07 | 3.12 | 0.9918 ± 0.0022 | 66 ± 22 | 1711 - 1719 (1.5) 1812 - 1836 (15.0) 1860 - 1866 (0.7) 1822 - 1925 (78.4) | Centennial | 1841-1955 |
| ANU | SANU-76438 | GWS-02 | Surface | *Ovis aries* | Tibia | 13.30 | -17.15 | 5.94 | 3.14 | 0.9797 ± 0.0021 | 165 ± 22 | 1676 - 1735 (30.9) 1800 - 1895 (49.2) 1904 - 1913 (1.6) 1919 - 1949* (18.3) | Centennial | 1841-1955 |
| ANU | SANU-76503 | GWS-19 | Surface | Ovicaprid | Tiba | 11.10 | -21.45 | 8.01 | 3.13 | 0.9819 ± 0.0021 | 147 ± 22 | 1692 - 1727 (21.17) 1808 - 1897 (54.09) 1901 - 1949* (24.74) | Centennial | 1841-1955 |
| ANU | SANU-76504 | GWS-32 | Surface | Ovicaprid | Ulna | 8.99 | -13.10 | 4.84 | 3.17 | 1.5212 ± 0.0033 | MODERN | 1963 - 1964 (9.4) 1964 - 1971 (86.0) | Decadal | 1963-1971 |
| ANU | SANU-76505 | GWS-36 | Surface | *Ovis aries* | Metatarsal | 8.11 | -22.37 | 6.17 | 3.10 | 0.9835 ± 0.0021 | 133 ± 23 | 1697 - 1725 (16.42) 1809 - 1872 (38.39) 1873 - 1947 (45.18) | Centennial | 1841-1955 |
| ANU | SANU-76506 | GWS-48 | Surface | Ovicaprid | Metatarsal | 2.57 | -21.63 | 5.25 | 3.10 | 0.9815 ± 0.0021 | 150 ± 22 | 1689 - 1729 (22.95) 1807 - 1897 (53.48) 1901 - 1916 (05.09) 1917 - 1949* (18.48) | Centennial | 1841-1955 |
| ANU | SANU-76509 | GWS-80 | Surface | Ovicaprid | Molar | 9.04 | -18.64 | 6.49 | 3.14 | 0.9898±0.0021 | 82 ± 22 | 1708 - 1720 (3.1) 1811 - 1838 (18.3) 1850 - 1867 (3.2) 1879 - 1927 (70.8) | Centennial | 1841-1955 |
| ANU | SANU-76510 | GWS-83 | Surface | Ovicaprid | Metatarsal | 6.37 | -21.84 | 6.35 | 3.17 | 0.9822 ± 0.0024 | 144 ± 25 | 1692 - 1727 (20.15) 1808 - 1949* (79.85) | Centennial | 1841-1955 |
| ANU | SANU-76511 | GWS-84 | Surface | Ovicaprid | Ulna | 2.54 | -22.19 | 5.96 | 3.11 | 0.9847±0.0021 | 124 ± 22 | 1698 - 724 (14.04) 1810 - 1839 (22.48) 1843 - 1871 (14.17) 1875 - 1946 (49.30) | Centennial | 1841-1955 |
| ANSTO | OZBU45 | GWW_04 | -2.8 | Ovicaprid | Tibia | 16.35 | -22.0 | 6.0 | 3.2 | 1.5536 ± 0.0045 | MODERN | 1964 - 1964 (9.0) 1968 - 1969 (86.4) | Decadal | 1964-1969 |
| ANSTO | OZBU44 | GWW_16 | -2.8 | *Bos* sp. | Rib | 17.34 | -22.9 | 6.0 | 3.3 | 1.5491 ± 0.0052 | MODERN | 1964 - 1964 (11.2) 1968 - 1969 (84.3) | Decadal | 1964-1969 |
| ANU | SANU-76513 | GWW_56 | -2.8 | *Sus domesticus* | Maxilla | 2.73 | -21.93 | 7.51 | 3.10 | 1.5276 ± 0.0032 | MODERN | 1964 - 1964 (7.8) 1969 - 1970 (87.6) | Decadal | 1964-1970 |
| ANSTO | OZBU41 | GWW_57 | -3.2 | Ovicaprid | Scapula | 15.3 | -22.6 | 6.1 | 3.2 | 1.5409 ± 0.0047 | MODERN | 1964 - 1964 (10.5) 1968 - 1968 (17.6) 1969 – 1970 (67.4) | Decadal | 1964-1970 |
| ANSTO | OZBP58 | GWW_58 | -3.8 | *Bos* sp. | Femur | 15.1 | -22.6 | 8.0 | 3.3 | 1.5731 ± 0.0052 | MODERN | 1964 - 1964 (5.8) 1967 - 1968 (89.6) | Decadal | 1964-1968 |
| ANSTO | OZBU43 | GWW_59 | -3.2 | *Bos* sp. | Humerus | 17.1 | -22.9 | 7.3 | 3.3 | 1.5503 ± 0.0052 | MODERN | 1964 - 1964 (11.3) 1968 - 1969 (84.2) | Decadal | 1964-1969 |
| ANSTO | OZBU40 | GWW_61 | -4.3 | Ovicaprid | Metacarpal | 15.51 | -22.2 | 9.1 | 3.2 | 0.9769 ± 0.0030 | 190 ± 25 | 1669 - 1715 (23.9) 1715 - 1784 (28.4) 1794 - 1816 (11.1) 1833 - 1892 (19.5) 1922 - 1942 (7.7) 1943 - 1945 (0.8) 1946 - 1950 (2.3) 1951 - 1955 (1.6) | Centennial | 1841-1955 |
| ANSTO | OZBP57 | GWW_62 | -3.8 | Ovicaprid | Humerus | 15.0 | -22.9 | 6.2 | 3.3 | 1.5301 ± 0.0042 | MODERN | 1964 - 1964 (8.9) 1969 - 1970 (86.6) | Decadal | 1964-1970 |
| ANU | SANU-76523 | GWW_70 | -4.6 | *Macropus* sp*.* | Tibia | 3.71 | -9.28 | 2.36 | 3.17 | 0.7827 ± 0.0020 | 1969 ± 25 | cal BC 38 - cal BC 23 (3.92) 21 - 128 (93.70) 185 - 196 (2.37) | Millennial | 38 cal BC-196 cal AD |
| ANSTO | OZBU39 | GWW_71 | -5.4 | *Bos* sp. | Ulna | 17.5 | -20.5 | 5.8 | 3.2 | 1.5616 ± 0.0051 | MODERN | 1964 - 1964 (14.0) 1967 - 1969 (81.4) | Decadal | 1964-1969 |
| ANSTO | OZBU42 | GWW_76 | -5.4 | Ovicaprid | Radius | 16.15 | -22.9 | 5.7 | 3.2 | 1.5758 ± 0.0052 | MODERN | 1964 - 1964 (5.4) 1967 - 1968 (90.0) | Decadal | 1964-1968 |
