## Supplementary Table 1 for "Neotaphonomic characteristics of vertebrate site formation in underwater caves"

**S1 Table ZooMS analysis and taxonomic ID across a sub-sample of taxon**

| **Condition** | **Site** | **No.** | **Zooms ID** | **Grid** | **Depth (-m)** | **Element** | **Morph. ID** | **ZooMS ID** | **Age Category** |
| --- | --- | --- | --- | --- | --- | --- | --- | --- | --- |
| Wet | GH | GH_09 | MTGZ1 | C6 | 15.4 | Metatarsal | Ovicaprid | *Ovis aries* | 1841-1955 CE |
| Wet | GH | GH_4 | MTGZ2 | C4 | 13.3 | Rib | Indet | *Macropus* sp. | 1841-1955 CE |
| Wet | GH | GH_18 | MTGZ3 | D6 | 0.3 | Tibia | Indet | *Macropus* sp. | 1841-1955 CE |
| Wet | GH | GH_26 | MTGZ4 | NA | 10 | Cranium | Canid | *Canis* sp. | 1456-1625 CE |
| Wet | GH | GH_28 | MTGZ5 | NA | 10 | Cranium | Macropod | *Macropus* sp. | 529-637 CE |
| Dry | GWS | GWS_36 | MTGZ6 | 4 | 0 | Metatarsal | Ovicaprid | *Ovis aries* | 1841-1955 CE |
| Dry | GWS | GWS_47 | MTGZ7 | 5 | 0 | Metatarsal | Ovicaprid | *Ovis aries* | 1841-1955 CE |
| Dry | GWS | GWS_60 | MTGZ8 | 6 | 0 | Femur | Unknown | *Ovis aries* | 1841-1955 CE |
| Dry | GWS | GWS_66 | MTGZ9 | 8 | 0 | Femur | Ovicaprid | *Ovis aries* | 1841-1955 CE |
| Dry | GWS | GWS_69 | MTGZ10 | 10 | 0 | Metatarsal | Ovicaprid | *Ovis aries* | 1841-1955 CE |
| Wet | GWW | GWW_16 | MTGZ11 | B3 | 2.8 | Rib | Ovicaprid | *Ovis aries* | 1964-1969 CE |
| Wet | GWW | GWW_56 | MTGZ12 | B3 | 2.8 | Cranium | Sus sp. | *Sus domesticus* | 1964-1970 CE |
| Wet | GWW | GWW_58 | MTGZ13 | B3 | 3.8 | Femur | Bos sp. | *Bos taurus* | 1964-1968 CE |
| Wet | GWW | GWW_59 | MTGZ14 | B3 | 3.2 | Humerus | Bos sp. | *Bos taurus* | 1964-1969 CE |
| Wet | GWW | GWW_71 | MTGZ15 | B3 | 5.4 | Ulna | Bos sp. | *Bos taurus* | 1964-1969 CE |
