## Supplementary Appendix Expanded Methods for "Neotaphonomic characteristics of vertebrate site formation in underwater caves"

**S1 Appendix: Expanded methods for *Neotaphonomic characteristics of vertebrate site formation in underwater caves***

### Chronological framework

Domesticate species were targeted to constrain chronologies to the European colonisation period in the region [1, 2]. For each burial context, a representative sheet/goat (*Ovis*/*Capra*) and cow (*Bos taurus*) were selected (Table 4). These species differ in in size, shape, microstructure, and composition, which allows for the assessment of whether degradation varies based on size classes. Other native and introduced taxa were analysed if they occupied areas of the cave not represented by domesticate species. Native taxa provide the capacity to extend the temporal scale by hundreds of years. By spatially controlling the samples within the same burial context, the study also aims to determine whether animals from the same collection locality were deposited at the same time.

At ANSTO, bone collagen was extracted using a modified Longin method and purified using ultrafiltration [3, 4]. At the ANU Radiocarbon Laboratory, samples were prepared following the protocol described in Wood et al. [5], with collagen extracted from bone and teeth purified using an ultrafiltration protocol based on Brock et al. [4]. Samples were not treated with solvent to remove lipids prior to demineralisation [6]. The preservation of collagen and suitability for dating were assessed using criteria described by van Klinken [7], and Wood [5], and we report percent collagen yield (wt%), δ^13^C (‰) and δ^15^N (‰), and the carbon to nitrogen atomic ratio (S2 Table). The carbon to nitrogen ratio and stable isotope composition were analysed using an ANCA GSL connected to a Sercon 20-22 IRMS at the ANU (for a description of standards and calculation methods see Wood et al. [5]). At ANSTO these were analysed by CF-EA/IRMS using an Elementar vario ISOTOPE select EA with an Elementar EcovisION IRMS following Beck et al. [8] To account for fractionation between carbon isotopes, the δ^13^C correction at ANU was made using an AMS derived value, and at ANSTO a correction was made using the IRMS analysis on the graphite from the sample. All samples report F14C [9] and the Conventional Radiocarbon age, rounded and reported in radiocarbon years using the Libby half-life of 5568 years [10].

Dating material from modern, historic and pre-1650s period required tailored approaches to address historical radiocarbon events. Radiocarbon ages with greater than 1.000 F14C were considered modern and calibrated against the Southern Hemisphere Bomb Curve Zone 1-2 (Bomb 21 SH1-2) using the online OxCal 4.4 online calibration software [11], measuring the period after 1955 [12]. Older radiocarbon ages were calibrated against the Southern Hemisphere calibration curve (SHCal20) [13] using Calib 8.10 software [14]. Calibrated age (cal BP) ranges are presented using the two-sigma range (95.4%) as this has the highest probability of the true age falling within this range.

The larger calibrated age ranges created by the Suess Effect was considered for older samples deposited during the industrialisation period, and before the intense period of nuclear weapons testing [15, 16]. During this period, a plateau region in the calibration curve occurs due to the influence of cosmic rays, and ^14^C depleted emissions from the Industrial Revolution that diluted atmospheric radiocarbon levels, resulting in multiple calibrated dates for a single radiocarbon age [17-20]. Taphonomic and diagenetic modifications within this period were identified as occurring after the first recorded introduction of domesticate species to the Mount Gambier region (1841 cal AD) [1, 2, 21] defining the lower chronological age. The upper age for historic specimens across each context was estimated as the beginning of the Bomb 21 SH1-2 (1955). Although this method refines the dates for older domesticate species for each location associated with European settlement, it is only an estimate of deposition in their current burial context. It is possible that these samples were deposited throughout this period, and bones were not deposited shortly after the death of the animal.

#### ZooMS

Fifteen specimens were selected for Zooarchaeology by Mass Spectrometry (ZooMS) analysis, using the acid insoluble protocol (following [22-24]). These included five sheep/goat, three cattle, and one each of pig, canid, and macropod, identified morphologically, as well as four unidentified remains (Tables 1 and 4, S1 Table). The specimens included a mix of dense limb bone (n=10), crania (n=3), and rib (n=3) fragments.

An approximately 20 mg of bone chip from each specimen was demineralized in 500 µL of 0.6 M hydrochloric acid (HCl) and stored in the fridge at 4 °C overnight, at which point the bones had become spongy/flexible and no bubbling was visible. After centrifuging, the acid supernatant was removed, and the samples were washed three times with 200 µl of 50 mM Ammonium Bicarbonate (AmBic) solution. The samples were then gelatinized in 100 µl AmBic at 65 °C for one hour. Following this, 50 µl of the solution was transferred to a new Eppendorf tube, with 1 µl of trypsin (0.4 µg/µl Pierce^TM^ Trypsin Protease, Thermo Scientific), and incubated overnight (~16 hours) at 37 °C. After incubation, 1 µl of 5% trifluoroacetic acid (TFA) was added to stop tryptic digestion. The samples were purified and desalted using a C18 Ziptip (Piece^TM^ C18 Tips, Thermo Scientific) into 10 µl of conditioning solution. A 0.5 µl aliquot was spotted in triplicate onto a MALDI 384 ground steel target plate (Bruker Daltonics) and mixed with 0.5 µl of matrix solution (α-Cyano*-*4*-*hydroxycinnamic acid).

Analysis was conducted at the Institute for Biomedicine and Glycomics, Griffith University, Australia, using a Bruker rapifleX (Bruker Daltonics) in reflector mode, positive polarity, with a mass-to-charge range 800–3500 m/z, laser intensity between 20–40%, and 2000 shots per sample. External mass calibration was performed using a peptide calibration standard (#8206195, Bruker Daltonics) containing a mixture of seven known peptides.

Raw MALDI spectra were converted to text and mzML formats in MSConvert (ProteoWizard). For each specimen, the triplicate spectra were then processed in R following the workflow outlined in Mylopotamitaki et al. [25] and using the MALDIquant and MALDIquantForeign packages [26]. Processing steps involved smoothing with a moving-average algorithm (half-window size = 2), baseline correction using the TapHat method (half-window size = 14), and spectra alignment using the SuperSmoother method (half-window size = 7, signal-to-noise cut off = 3). After alignment, the triplicates were merged, and a second TopHat baseline correction was applied. The resulting spectra were exported as .msd files and imported into mMass [27] for peak picking using a signal-to-noise threshold of 3.5. Taxonomic identifications were made by comparing peaks to those in a custom database of known peaks from Australia endemic and introduced fauna [22, 28].

ZooMS analysis identified all samples to at least the genus level, in the case of *Macropus* and *Canis*, or to the species level in the case of introduced domesticates. All specimens identified morphologically as ovicaprid were identified as sheep (*Ovis aries*). The four morphologically unidentifiable specimens were identified using ZooMS, two as sheep, and the other two as kangaroo (*Macropus*), distinguishable from other macropodids based on peaks at m/z 2989 (COL1a2 10–42) and m/z 2897 and 2913 (COL1a1 586–618).

**Table 1. Results of the ZooMS analysis with the 11 collagen peptide markers used for identifications**

| Sample ID | Morph. ID | ZooMS ID | COL1a1  508–519 | COL1a2  978–990 | COL1a2  484–498 | COL1a2  502–519 | COL1a2  292–309 | COL1a2  889–906 | COL1a2  793–816 | COL1a2  454–483 | COL1a1  586–618 | COL1a2  757–789 | COL1a2  10-42 |
| --- | --- | --- | --- | --- | --- | --- | --- | --- | --- | --- | --- | --- | --- |
| GH_09 | Indet. | *Ovis aries* | 1105 | 1180 + 1196 | 1427 | 1580 | 1648 | - | 2131 | 2792 | 2883 + 2899 | 3017 + 3033 | - |
| GH_4 | Indet. | *Macropus sp.* | 1162 | 1150 + 1166 | 1453 | 1598 | 1680 | 1652 | 2145 | - | 2897 + 2913 | 2943 + 2959 | 2989 |
| GH_18 | Indet. | *Macropus sp.* | 1162 | 1150 + 1166 | 1453 | 1598 | 1680 | 1652 | 2145 | - | 2897 + 2913 | 2943 + 2959 | 2989 |
| GH_26 | Canid | *Canis* sp. | 1105 | 1210 + 1226 | 1453 | - | - | - | 2131 | 2820 | 2853 + 2869 | 2983 + 2899 | - |
| GH_28 | Macropod | *Macropus sp.* | 1162 | 1150 + 1166 | 1453 | 1598 | 1680 | 1652 | 2145 | - | 2897 + 2913 | 2943 + 2959 | 2989 |
| GWS_36 | *Ovis/Capra* | *Ovis aries* | 1105 | 1180 + 1196 | 1427 | 1580 | 1648 | - | 2131 | 2792 | 2883 + 2899 | 3017 + 3033 | - |
| GWS_47 | *Ovis/Capra* | *Ovis aries* | 1105 | 1180 + 1196 | 1427 | 1580 | 1648 | - | 2131 | 2792 | 2883 + 2899 | 3017 + 3033 | - |
| GWS_60 | Indet. | *Ovis aries* | 1105 | 1180 + 1196 | 1427 | 1580 | 1648 | - | 2131 | 2792 | 2883 + 2899 | 3017 + 3033 | - |
| GWS_66 | *Ovis/Capra* | *Ovis aries* | 1105 | 1180 + 1196 | 1427 | - | 1648 | - | 2131 | 2792 | 2883 + 2899 | 3017 + 3033 | - |
| GWS_69 | *Ovis/Capra* | *Ovis aries* | 1105 | 1180 + 1196 | 1427 | 1580 | 1648 | - | 2131 | 2792 | 2883 + 2899 | 3017 + 3033 | - |
| GWW_16 | *Ovis/Capra* | *Ovis aries* | 1105 | - | 1427 | 1580 | 1648 | - | 2131 | 2792 | 2883 + 2899 | 3017 + 3033 | - |
| GWW_56 | *Sus sp.* | *Sus domesticus* | 1105 | 1180 + 1196 | 1453 | 1550 | 1647 |  | 2131 | 2820 | 2883 + 2899 | 3017 + 3033 | - |
| GWW_58 | *Bos sp.* | *Bos taurus* | 1105 | 1192 + 1208 | 1427 | 1580 | 1648 | - | 2131 | 2792 | 2853 + 2689 | 3017 + 3033 | - |
| GWW_59 | *Bos sp.* | *Bos taurus* | 1105 | 1180 + 1196 | 1453 | 1550 | 1647 |  | 2131 | 2820 | 2883 + 2899 | 3017 + 3033 | - |
| GWW_71 | *Bos sp.* | *Bos taurus* | 1105 | 1192 + 1208 | 1427 | 1580 | 1648 | - | 2131 | 2792 | 2853 + 2689 | 3017 + 3033 | - |

#### Bone surface modifications

Bone surface modifications were recorded based on previously developed systems, recording the quantitative and qualitative data to understand local agents. Previously published qualitative methodologies are summarised in Table 2. Presence and absence of features were further recorded to understand bone surface modifications, alterations to shape, penetration into bone tissue, and loss of matrix [29]. These data are provided in S1 Dataset.

Corrosion features were further described by the depth and spread of corrosion. Superficial corrosion is the light removal of periosteal bone surface, approximately less than 0.5 mm, whilst deep corrosion penetrated the bone matrix approximately more than 0.5 mm. Spread is distinguished by a continuous corrosion, where a single uniform corrosion feature spread across bone surfaces, compared to discontinuous corrosion where a feature is mottled and disjointed. Combination of these attributes across corrosion types (surface, pitting, etching, bone loss) aimed to understand the impact and characteristics of corrosion features.

Images of surface modifications were taking using a Nikon Z50 camera and Nikkor Z MC 50mm macro lens, and a Dino-lite AM7915MZT.

**Table 2. Bone modification analytical methods**

| **Measure** | **Data type** | **Measure** | **Reference** |
| --- | --- | --- | --- |
| Completeness | Nominal | Complete (C), Shaft (S), Distal (D), Proximal (P) | [30] |
| Fragment size class | Ordinal | 1) <2mm; 2) 2-4mm; 3) 4-8mm; 4) 8-16mm; 5) 16-32mm; 6) 32-128mm; 7) 128-256mm; 8) 256-512mm; 9) 512-1024; 10) 1024-2048; 11) >2048mm | [31] |
| Breakage index (BI) | Ordinal | 1) <10%; 2) 10-20%; 3) 20-30%; 4) 30-40%; 5) 40-50%; 6) 50-60%; 7) 60-70%; 8) 70-80%; 9) 80-90%; 10) 90-100% bone remaining | [32] |
| Shape | Nominal | Class 1: Tabular  Class 2: Elongate  Class 3: quidimensional/compact  Class 4: conical | [33] |
| Shape (Aspect Ratio) | Continuous | Maximum length: maximum breadth | [31] |
| Shaft circumference (Circ) | Ordinal | 1) Bone circ. <1/2  2) Circ. >1.2 in at least a portion  3) Complete in at least a portion | [34] |
| Shaft fragmentation | Ordinal | 1) <1/4 the original length  2) 1/4-1/2 the original length  3) 1/2-3/4 the original length  4) >3/4 the original length | [34] |
| Weathering stages | Ordinal | 1) Bone surface presents no cracking or flaking  2) Surface shows cracking, typically parallel to the orientation of collagen fibres. A mosaic cracking pattern may be observed on the articular surfaces  3) Surface shows flaking, typically along the edge of cracks. No rounding is observed on the cracked edges  4) Surfaces present roughened patches from flaking of the periosteal surface, only to 1.0-1.5mm depth. Cracks typically rounded  5) Rough bone surfaces with splinters. Wide cracks, rounded or actively splintering edges  6) Bone disintegrating into splinters, original shape may be obscured. | [35] |
| Abrasion scale | Ordinal | 0) fresh and unabraded/ Sharp and well-defined processes and edged  1) Slight abrasion where edges and processes have slight rounding  2) Moderate abrasion with well rounded edges. Processes are recognised as protrusions on the bone. Possible presence of a polish  3) All edges are extremely well rounded, and processes are no longer present or at best are remnants. Barely recognisable as a one fragment | [36] |

#### Portable X-Ray Fluorescence

Following standard protocols [37, 38] assays were collected by a Bruker Tracer 5i portable XRF spectrometer with a silicon drift detector, Rh target (35kv max voltage, 50µA). Beam-phase parameters collected assays between consecutive 23 - 27 second live count, collecting 32 elements. Each spectra represents an average elemental composition across an ovid shaped area approximately 7x5 mm in size. Certified Reference Material (CRM) 45d (SiO_2_) was selected to calibrate the instrument to ensure precision could be quantified, data accuracy was monitored, and instrumental factors could be quantified [39].

A total of 11 spectra were collected: three spectra were assayed from CRM401, six spectra tested areas of localised and broadscale black staining across the two specimens, and two spectra identify the composition of exposed bone surfaces.

#### Histological preparation

Twenty-four bone specimens were selected for histological analysis (Table 4). Standard protocols were followed to make epoxy embedded blocks and thin sections for backscatter scanning electron microscopy and histological analysis [40, 41].

Using a Dremel® Variable-Speed Rotary Tool with a Dremel® cut off wheel, 1cm transverse cortical bone samples were extracted from thick cortical bone across different regions of the shaft depending on prior breakage points and bone surface modifications (BSMs). For complete and unmodified bones, samples were taken from the midshaft. This was so that the sampling location would capture taphonomic BSMs and the Haversian system in cortical bone. After embedding in Buehler EpoxiCure^TM^ epoxy resin, samples were further reduced transversely using an Allied Techcut 4^TM^ low-speed saw and Kemet Diamond Wheel to approximately 3 mm. The remaining embedded sample was retained for scanning electron microscopy (SEM) analysis.

Surfaces of the reduced sections were polished using 600/P1200 grit wet/dry grinding pads and mounted onto glass microscope slides using Selleys® ultra-clear Araldite epoxy adhesive. Samples were manually ground to approximately 100 µm +/- 90 µm thickness. Once histological features were visible under the microscope, the sections were further polished using Buehler Micropolish II 0.3 µm powder and Beuhler polishing cloth. Samples for SEM imaging followed the same polishing process. Slides were cleaned in an ultrasonic water bath, dehydrated in a series of ethanol baths, and cleared using xylene before the cover slip was applied using DPX mounting medium.

Samples for microscopy were analysed under transmitted and polarised optical light using an Olympus BX63 high powered microscope, with 10x eyepiece magnification, to assess collagen preservation and histotaphonomy. Complete composite images were taken at 10x, with regions of interest captured at additional magnifications as required. Backscatter SEM analysis was undertaken using a Hitachi TM3030 SEM_EDS. Samples were imaged in BSE mode (standard or reduced charge up), under 15kv voltage, with high charge. Composite images were developed at either X40 and X60 magnification, combined using Photoshop 2026, and targeted images taken at various magnification.

Measures of the Oxford Histological Index (OHI) [42, 43] and its derivatives to assess specific microscopic focal destruction features [44] are outlined in Table 3. Descriptions of degradation indices provide a summary of what is typically seen across bone sections for each category (Table 3). Percentages following Booth [45] were used to calculate OHI, Bacterial Attack Index (BAI) and Cyanobacterial Attack Index (CAI) as other studies did not account for degradation between 33 – 67% [44]. S1 Dataset reports on the total cortical area measured (Ct.Ar), percentage preserved used to calculate the OHI %BAI and %CIA, peripheral degradation penetration data measurements, and general descriptions of the histotaphonomy of each sample.

**Table 3. Histological Index followed to record the Oxford Histological Index (OHI), Bacterial Attack Index (BAI), and Cyanobacteria Attack Index (CAI) [44, 45].**

| **Index** | **%** | **OHI (general), BAI (bacterial Microscopic Focal Destruction only), CAI (cyanobacteria degradation only)** |
| --- | --- | --- |
| 0 | <5% | *Very well preserved, virtually indistinguishable from fresh bone* |
| 1 | <15% | *Only minor amounts of destructive foci, otherwise generally well preserved* |
| 2 | <50% | *Clear preservation of some osteocyte lacunae* |
| 3 | >50% | *Clear lamellate structure preserved between destructive foci* |
| 4 | >85% | *Small areas of well-preserved bone, or some lamellar structure preserved by pattern of destructive foci* |
| 5 | >95% | *No original features identifiable, other than Haversian canals* |

**Table 4. Specimen list for radiocarbon dating (14C), ZooMS and Histological analysis.**

| **Site** | **Wet/Dry** | **Sample No.** | **Depth** | **Taxon** | **Element** | **14C** | **14C Group** | **ZooMs** | **Histology** | **SEM Code*** |
| --- | --- | --- | --- | --- | --- | --- | --- | --- | --- | --- |
| GH | Wet | GH_1 | -10.1 | *Ovis aries* | Cranium | Y | Centennial | - | - | - |
| GH | Wet | GH_04 | -13.3 | *Macropus sp.* | Rib | Y | Centennial | Y | Y | HM |
| GH | Wet | GH_06 | -18.3 | Mammal | Vertebra | Y | Centennial | - | - | - |
| GH | Wet | GH_09 | -15.4 | Ovis *aries* | Metatarsal | Y | Centennial | Y | Y | NM |
| GH | Wet | GH_10 | -14 | Ovicaprid | Metatarsal | Y | Centennial | - | Y | NM |
| GH | Wet | GH_16 | -16.9 | Mammal | Vertebra | Y | Millennial | - | - | - |
| GH | Wet | GH_18 | -0.3 | *Macropus sp* | Tibia | Y | Centennial | Y | Y | NM |
| GH | Wet | GH_19 | -8.9 | Ovicaprid | Metatarsal | Y | Centennial | - | Y | HL |
| GH | Wet | GH-22 | -17.2 | Bos. sp | Vertebra | Y | Centennial | - | - | - |
| GH | Wet | GH-23 | -10.1 | Canis (Dingo) | Mandible | Y | Centennial | - | - | - |
| GH | Dry | GH-24 | surface | Ovis aries | Cranium | Y | Centennial | - | - | - |
| GH | Wet | GH-26 | -10 | Canid/ Carnivore | Cranium | Y | Centennial | Y | - | - |
| GH | Wet | GH-27 | -10 | Possum | Cranium | Y | Centennial | - | - | - |
| GH | Wet | GH-28 | -10 | *Macropus sp* | Cranium | Y | Millennial | Y | - | - |
| GWS | Dry | GWS_02 | 0 | Ovis aries | Tibia | Y | Centennial | - | Y | HM |
| GWS | Dry | GWS_19 | 0 | Ovicaprid | Tibia | Y | Centennial | - | Y | HL |
| GWS | Dry | GWS_32 | 0 | Ovicaprid | Ulna | Y | Decadal | - | - | - |
| GWS | Dry | GWS_36 | 0 | *Ovis aries* | Metatarsal | Y | Centennial | Y | Y | HL |
| GWS | Dry | GWS_47 | 0 | Ovicaprid | Metatarsal | Y | Centennial | Y | Y | HM |
| GWS | Dry | GWS_48 | 0 | Ovicaprid | Metatarsal | Y | Centennial | - | Y | HM |
| GWS | Dry | GWS_60 | 0 | Ovicaprid | Tibia | Y | Centennial | Y | Y | HM |
| GWS | Dry | GWS_66 | 0 | Ovicaprid | Femur | Y | Centennial | Y | Y | HM |
| GWS | Dry | GWS_69 | 0 | Ovicaprid | Metatarsal | Y | Centennial | Y | Y | HL |
| GWS | Dry | GWS_80 | 0 | Ovicaprid | Molar (dentine) | Y | Centennial | - | - | - |
| GWS | Dry | GWS_83 | 0 | Ovicaprid | Metatarsal | Y | Centennial | - | Y | HL |
| GWS | Dry | GWS_84 | 0 | Ovicaprid | Ulna | Y | Centennial | - | - | - |
| GWS | Dry | GWS_86 | 0 | Ovicaprid | Tibia | Y | Centennial | - | Y | HL |
| GWW | Wet | GWW_4 | -2.8 | Ovicaprid | Tibia | Y | Decadal | - | Y | HL |
| GWW | Wet | GWW_16 | -2.8 | *Bos sp.* | Rib | Y | Decadal | Y | Y | HL |
| GWW | Wet | GWS_56 | -2.8 | *Sus domesticus* | Maxilla | Y | Decadal | Y | - | - |
| GWW | Wet | GWS_57 | -3.2 | Ovicaprid | Scapula | Y | Decadal | - | - | - |
| GWW | Wet | GWW_58 | -3.8 | *Bos sp.* | Femur | Y | Decadal | Y | Y | HL |
| GWW | Wet | GWW_59 | -3.2 | *Bos sp.* | Humerus | Y | Decadal | Y | Y | HL |
| GWW | Wet | GWW_61 | -4.3 | Ovicaprid | Metacarpal | Y | Centennial | - | Y | HL |
| GWW | Wet | GWW_62 | -3.8 | Ovicaprid | Humerus | Y | Decadal | - | Y | HM |
| GWW | Wet | GWW_70 | -4.6 | *Macropus sp.* | Tibia | Y | Millennial | - | Y | HL |
| GWW | Wet | GWW_71 | -5.4 | *Bos sp.* | Ulna | Y | Decadal | Y | Y | HL |
| GWW | Wet | GWW_74 | -5.4 | *Macropus sp.* | Humerus | Y | FAILED | - | - | - |
| GWW | Wet | GWW_76 | -5.4 | Ovicaprid | Radius | Y | Decadal | - | Y | HL |
| *SEM Codes refer to beam condition and charge up. HL: EDX (15Kv large current), charge up reduction mode; HM: EDX, standard mode; NM: 15Kv, standard mode. | | | | | | | | | | |

### S1 Appendix References

1.	Hill LR. Mount Gambier: the city around a cave. Leabrook, SA: Investigator Press; 1972.

2.	MacGillivary L. We have found our paradise: The South-East squattocracy, 1840–1870. J. Hist. Soc. S. Aust.. 1989;17:25-38.

3.	Brock F, Bronk Ramsey C, Higham T. Quality Assurance of Ultrafiltered Bone Dating. Radiocarbon. 2007;49(2):187-92.

4.	Brock F, Higham T, Ditchfield P, Bronk Ramsey C. Current Pretreatment Methods for AMS Radiocarbon Dating at the Oxford Radiocarbon Accelerator Unit (Orau). Radiocarbon. 2010;52(1):103-12.

5.	Wood R, Esmay R, Usher E, Fallon S. SAMPLE PREPARATION METHODS USED AT THE AUSTRALIAN NATIONAL UNIVERSITY RADIOCARBON FACILITY. Radiocarbon. 2023;65(2):573-89.

6.	Krause MB, Vang N, Ahlmeyer L, Tung TA. Effects of lipid extraction on human bone collagen: Comparing stable carbon and nitrogen isotope values with and without lipid extraction. J. Archaeol. Sci. Rep. 2023;51:104196:.

7.	van Klinken GJ. Bone Collagen Quality Indicators for Palaeodietary and Radiocarbon Measurements. J. Archaeol. Sci.. 1999;26(6):687-95.

8.	Beck KK, Fletcher M-S, Kattel G, Barry LA, Gadd PS, Heijnis H, et al. The indirect response of an aquatic ecosystem to long-term climate-driven terrestrial vegetation in a subalpine temperate lake. J Biogeogr. 2018;45:713–725.

9.	Reimer PJ, Brown TA, Reimer RW. Discussion: Reporting and Calibration of Post-Bomb 14C Data. Radiocarbon. 2016;46(3):1299-304.

10.	Stuiver M, Polach HA. Discussion: Reporting of 14C data. Radiocarbon. 1977;19(3):355-63.

11.	Bronk Ramsay C. Bayesian analysis of radiocarbon dates. Radiocarbon. 2009;51(1): 337–60.

12.	Hua Q, Turnbull JC, Santos GM, Rakowski AZ, Ancapichún S, De Pol-Holz R et al. Atmospheric radiocarbon for the period 1950-2019. Radiocarbon. 2022;64(4):723-45.

13.	Hogg AG, Heaton TJ, Hua Q, Palmer JG, Turney CSM, Southon J, et al. SHCal20 Southern Hemisphere Calibration, 0–55,000 Years cal BP. Radiocarbon. 2020;62(4):759 - 78.

14.	Stuiver M, Reimer PJ. Extended 14C Data Base and Revised CALIB 3.0 14C Age Calibration Program. Radiocarbon. 1993;35(1):215 - 30.

15.	Jull AJT, Burr GS, Hodgins GWL. Radiocarbon dating, reservoir effects, and calibration. Quaternary International. 2013;299:64-71.

16.	Hua Q. Radiocarbon: A chronological tool for the recent past. Quaternary Geochronology. 2009;4:378-90.

17.	Thompson VD, Jefferies RW, Moore CR. The Case for Radiocarbon Dating and Bayesian Analysis in Historical Archaeology. Hist. Archaeol. 2018;53(1):181-92.

18.	Paterne M, Michel É, et Jean-Claude Dutay CH. Carbon-14. In: Ramstein G, Bouttes N, Govin A, Landais A, Sepulchre P, editors. Paleoclimatology. Switzerland: Springer; 2020. p. 51-71.

19.	Suess HE. The Radiocarbon Record in Tree Rings of the Last 8000 years. Radiocarbon. 1980;22(2):200-9.

20.	Suess HE. Radiocarbon concentration in modern wood. Science. 1955;122(3166):415-7.

21.	Unknown Author. Early History of Mount Gambier. The Border Watch. 1946 Vov 30:8

22.	Buckley M, Collins M, Thomas-Oates J, Wilson JC. Species identification by analysis of bone collagen using matrix-assisted laser desorption/ionisation time-of-flight mass spectrometry. Rapid Commun. Mass Spectrom. 2009;23(23):3843-54.

23.	Welker F, Soressi M, Rendu W, Hublin J-J, Collins M. Using ZooMS to identify fragmentary bone from the Late Middle/Early Upper Palaeolithic sequence of Les Cottés, France. J. Archaeol. Sci.. 2015;54:279-86.

24.	Zooarchaeology by Mass Spectrometry (ZooMS)- Pretreatment protocols for bone material. [Internet]. 2020.

25.	Mylopotamitaki D, Weiss M, Fewlass H, Zavala EI, Rougier H, Sumer AP, et al. Homo sapiens reached the higher latitudes of Europe by 45,000 years ago. Nature. 2024;626(7998):341-6.

26.	Gibb S, Strimmer K. MALDIquant: a versatile R package for the analysis of mass spectrometry data. Bioinformatics. 2012;28(17):2270-1.

27.	Strohalm M, Kavan D, Novák P, Volný M, Havlíček V. mMass 3: a crossplatform software environment for precise analysis of mass spectrometric data. Anal.l Chem. 2010;82:4648–51.

28.	Peters C, Richter KK, Manne T, Dortch J, Paterson A, Travouillon K, et al. Species identification of Australian marsupials using collagen fingerprinting. R. Soc. Open Sci. 2021;8:211229.

29.	Fernández-Jalvo Y, Andrews P. Atlas of Taphonomic Identifications: Springer; 2016.

30.	Andrews P. Owls, Caves and Fossils. Chicago: The University of Chicago Press; 1990.

31.	O'Connor TP. The Archaeology of Animal Bones. Stroud: Sutton Publishing Ltd; 2008.

32.	Andrews P, Molleson T, Boz B. The Human Burials at Çatalhöyük. In: Hodder I, editor. Inhabiting Çatalhöyük reports from the 1995-99 seasons. McDonald Institute for Archaeological Research and British Institute of Archaeology at Ankara Cambridge, London; 2005. p. 261.

33.	Blob RW, Fiorillo AR. The significance of vertebrate microfossil size and shape distributions for faunal abundance reconstructions: a Late Cretaceous example. Paleobiology. 1996;22(3):422-35.

34.	Villa P, Mahieu E. Breakage patterns of human long bones. J. Hum. Evol. 1991;21:27-48.

35.	Behrensmeyer AK. Taphonomic and ecologic information from bone weathering. Paleobiology. 1978;4(2):150-62.

36.	Fiorillo AR. Taphonomy of Hazard Homestead Quarry (Ogallala Group), Hitchcock County, Nebraska. Rocky Mountain Geology. 1988;26(2):57-97.

37.	Huntley J. Taphonomy Or Paint Recipe: In situ portable x-ray fluorescence analysis of two anthropomorphic motifs from the Woronora Plateau, New South Wales. Aust. Archaeol. 2016;75(1):78-94.

38.	Huntley J, Aubert M, Ross J, Brand HEA, Morwood MJ. One Colour, (at Least) Two Minerals: A Study of Mulberry Rock Art Pigment and a Mulberry Pigment ‘Quarry’ from the Kimberley, Northern Australia. Archaeometry. 2015;57(1):77-99.

39.	Huntley J, Wallis LA, Stephenson B, Corporation KNA, Davis A. A multi-technique approach to contextualising painted rock art in the Central Pilbara of Western Australia: Integrating in-field and laboratory methods. Quat. Int. 2021;572:52-73.

40.	Walker MM, Louys J, Herries AIR, Price GJ, Miszkiewicz JJ. Humerus midshaft histology in a modern and fossil wombat. Aust. Mammal. 2021;43(1).

41.	Schotsmans EMJ, Stuart BH, Stewart TJ, Thomas PS, Miszkiewicz JJ. Unravelling taphono-myths. First large-scale study of histotaphonomic changes and diagenesis in bone from modern surface depositions. PLoS One. 2024;19(9):e0308440.

42.	Hedges REM, Millard AR, Pike AWG. Measurements and Relationships of Diagenetic Alteration of Bone from Three Archaeological Sites. J. Archaeol. Sci.. 1995;22(2):201-9.

43.	Millard A. The deterioration of bone. In: Brothwell D, Pollard AM, editors. Handbook of archaeological sciences. Chichester: John Wiley & Sons Ltd; 2001. p. 637-47.

44.	Brönnimann D, Portmann C, Pichler SL, Booth TJ, Röder B, Vach W, et al. Contextualising the dead – Combining geoarchaeology and osteo-anthropology in a new multi-focus approach in bone histotaphonomy. J. Archaeol. Sci.. 2018;98:45-58.

45.	Booth T. The Rot Sets In: Low-Powered Microscopic Investigation of Taphonomic Changes to Bone Microstructure and its Application to Funerary Contexts. In: Errickson DaT, T, editor. Human Remains Another Dimension: The Application of Imaging to the Study of Human Remains. London, United Kingdom: Elsevier; 2017. p. 7-27.
